# Neuro-Immune Crypt-Associated Cells and REST-Mediated Reprogramming: Pathogen-Driven Stromal Activation, HERVs Induction, and Abortive Antiviral Signaling in Colorectal Carcinoma

**DOI:** 10.1101/2025.10.21.682573

**Authors:** David Díaz-Carballo, Adrien Noa-Bolaño, Udo Rahner, Ali H. Acikelli, Sahitya Saka, Jacqueline Klein, Flevy D’Souza, Sascha Malak, Anne Höppner, Annabelle Kamitz, Carla Casula, Lalitha Kamada, Andrea Tannapfel, Jens Christmann, Sarah Albus, Enrico Fruth, Daniela Gerovska, Marcos J. Araúzo-Bravo, Metin Senkal, Crista Ochsenfarth, Sven P. Berres, Julian Uszkoreit, Dirk Strumberg

## Abstract

Colorectal cancer (CRC) pathogenesis remains linked to poorly defined cellular origins and microenvironmental drivers. We identify Neuro-Immune Crypt-Associated (NICA) cells as a privileged pathogen portal and a plausible epithelial cell of origin. During carcinogenesis, the repressor REST drives a neuroendocrine-to-epithelial transition by silencing NICA-associated neuroendocrine markers, a lineage loss reversible via epigenetic modulation or genetic ablation in CRC models. This landscape is further shaped by EBV-infected B-lineage cells (BLEICS), which transactivate HERV-H/F elements in CRC cells through paracrine signaling, establishing a niche-restricted ‘viral scar’. While tumor crypts sense this retroviral pressure, they exhibit an abortive antiviral signature marked by RNase L downregulation, creating a functional execution gap. We propose a pathogen-first model where chronic inflammatory pressure triggers ‘mutation-silencing’ -the simultaneous genotoxic and epigenetic inactivation of tumor suppressors. Together, the convergence of NICA plasticity, BLEICS-mediated HERV induction, and compromised antiviral surveillance redefines CRC as a pathogen-driven disruption of the neuro-immune niche.

## Introduction

Colorectal cancer (CRC) is a leading global malignancy with rising incidence in younger patients^1^. While the canonical model emphasizes mutations in crypt stem cells, emerging evidence highlights additional cellular and microenvironmental drivers^2^.

The classical Vogelstein model has long framed colorectal tumorigenesis, linking recurrent *APC*, *KRAS*, and *TP53* mutations in LGR5 intestinal stem cells to defined stages of progression. While foundational, this paradigm is inferred from cross-sectional enrichment rather than direct clonal tracking, limiting its universality. Mutation frequencies highlight this: *APC* mutations occur in ≈70-80% of CRCs, *TP53* in ≈50-60%, and *KRAS* in ≈30-40%. Moreover, MSI-high tumors (≈15%) bypass *APC*/*KRAS*/*TP53* altogether through mismatch repair deficiency, *BRAF* mutation, or epigenetic changes^3^. The predominance of *KRAS*-wildtype tumors indicates that malignant transformation can proceed without KRAS activation, underscoring pathway deregulation rather than single-node mutation as the critical driver^4,5^.An alternative scenario is that pathway deregulation precedes mutation, with chronic activation of growth signals creating selective pressure for stabilizing lesions. Mutations then act as maladaptive locks on deregulated states rather than true initiators^6,7^. Functionally equivalent hits converge on persistent β-catenin signaling, including *APC* truncation, *CTNNB1* stabilization, *AXIN1*/2 loss, *RNF43*/*ZNRF3 inactivation*, or *RSPO* fusions. Viral proteins further contributes to modulate WNT/β-catenin pathways: EBV^8,9^, HBV^10^, HPV^11^, HTLV 1^12^, and KSHV^13^, while *TP53* inactivation may result from *MDM2* amplification, epigenetic silencing^14^, or viral proteins from HPV, adenoviruses, EBV, HBV, or Merkel cell polyomavirus^15,16^. Beyond viral modulation, several microorganisms also contribute to Wnt/β-catenin deregulation^17^. Salmonella enterica and Fusobacterium nucleatum activate β-catenin signaling to promote epithelial proliferation and survival^17^, whereas pks^+^ Escherichia coli produce the genotoxin colibactin, which induces DNA double strand breaks and generates a characteristic mutational signature also detected in human CRC, including truncating APC mutations^18^. S. enterica, Mycobacterium tuberculosis, and certain viruses are preferentially recognized by intestinal Microfold cells (M-cells), which mediate pathogen engulfment^19^. Consistent with this model, F. nucleatum is enriched in colorectal tumors compared to adjacent mucosa^20^. Together, these observations suggest viral and microbial infections can deregulate oncogenic pathways such as Wnt/β-catenin before stabilizing mutations occur. While genetic and viral mechanisms underscore the role of pathway deregulation, they do not explain CRC’s anatomical distribution, which correlates with regions enriched in neuroendocrine cells and lymphoid follicles^21,22^. This spatial pattern suggests lineage-specific contributors, particularly neuroendocrine cells, may participate in tumor initiation^23^.

Neuroendocrine (NE) cells are a specialized epithelial lineage producing neuronal markers and bioactive peptides that regulate endocrine and paracrine signaling^24^. They are present in most organs, with highest density in the gastrointestinal^25^ and respiratory tracts^26^. Approximately 70% of neuroendocrine neoplasms (NENs) arise within the gastrointestinal tract -predominantly in the ileum and rectum-while about 25% originate in the respiratory system^27^. In CRC, NE cells are detected in ≈34% of cases and may be even more frequent in preneoplastic lesions, underscoring their role in carcinogenesis^28,29^. In the colon, the distribution of neuroendocrine cells is regionally heterogeneous. Enterochromaffin (EC) cells are more abundant in the right colon and decrease toward the left side. In contrast, L-, D-, and I-type enteroendocrine cells are enriched in the left colon. Neuroendocrine cells do not follow the 3-5 day epithelial turnover cycle; they are long lived, slow renewing cells that persist for weeks to months. Overall, the left colon contains a higher total number of neuroendocrine cells compared with the right colon. Along the gut associated lymphoid tissue (GALT), the small intestine contains more diffuse and extensive Peyer’s patches, particularly in the ileum. In contrast, isolated lymphoid follicles are numerically more abundant in the colon, with a higher density in the left colon compared with the right. Notably, CRC occurs most often in the descending colon, sigmoid, and rectum -enriched in NE cells and isolated lymphoid follicles-suggesting that local microenvironments contribute to malignant transformation^30,31^.

The colonic mucosa is among the most rapidly regenerating tissues, sustained by the dynamic activity of Lieberkühn crypts. These crypts comprise distinct compartments: the LGR5 crypt base columnar (CBC) stem cell niche, an intermediate transit amplifying zone, and differentiated mature cells^32^. Emerging evidence shows stemness is not confined to the crypt base; under stress or injury, nearly all crypt cells can acquire facultative stem cell properties^33,34^. In line with the stem cell theory of cancer^35^, such plasticity suggests that multiple cell types -including those beyond the CBC niche-may initiate malignancy^33^.

We report for the first time that a subset of colonic neuroendocrine (NE) cells displays a distinct neuroimmune phenotype, designated Neuro Immune Crypt Associated (NICA) cells. These cells co-express neuronal markers [CHGA, Tyrosine hydroxylase (TH), Peripherin, Somatostatin, Neuron specific enolase (NSE)], stemness determinants (LGR5, CD133, GLUT3, ASCL2, β-catenin), and epithelial Keratin 80 (KRT80). They also express innate immune molecules (TLR4, REG4, PigR, TNFαIP2, Viperin, DEFA6, SDF 1) and antigen presenting proteins (CD22, CD80), while lacking CD45. Their localization near follicle associated epithelia suggests integration of neural, regenerative, and immune functions in mucosal homeostasis. By analogy, NICA cells share features with M-cells yet remain distinct in their neuroimmune identity, co expressing neuronal markers (CHGA, TH), immune sensors (TLR4), and stemness determinants (LGR5).

Tumor crypts show a marked loss of neuronal gene expression (≈70-75%) with concomitant REST upregulation. REST (*NRSF*)^36^ is a direct transcriptional target of Wnt/β-catenin signaling in the developing spinal cord^37^, providing a mechanistic rationale for its activation in CRC. We propose that REST acts as a molecular switch that enforces the loss of NICA-specific neuroendocrine signatures during the transition to a malignant epithelial state. The Wnt REST axis thus bridges pathway deregulation and transcriptional reprogramming. Despite dedifferentiation, tumors retain neuronal and stemness markers plus KRT80, supporting derivation from an original NICA lineage. NICA cells also express viral entry receptors (GLUT1, EPHA2, CD80, NgR1), rendering them susceptible to oncogenic infection. Such epigenetic relaxation and viral susceptibility create conditions permissive for the activation of endogenous retroviral elements.

Human endogenous retroviruses (HERVs) constitute a large class of paleoviral insertions acquired through ancient germ line infections and vertically transmitted in a Mendelian fashion. Although most loci have accumulated, disabling mutations or are epigenetically repressed, their canonical 5′-LTR-gag-pol-env-LTR 3′ structure remains recognizable, and some families retain transcriptional competence under conditions that erode heterochromatic control. In particular, some envelope proteins HERV-derived are implicated in fusogenic and immunosuppressive effects^38^. HERV promoters can be transactivated by exogenous viral infections and are highly responsive to chromatin modifying agents such as histone deacetylase inhibitors (HDACi). This pharmacologic inducibility has been exploited in latency reversal strategies, including the use of TLR7 agonists such as vesatolimod, which activate endogenous retroviral elements as part of a broader innate immune stimulation. Aberrant HERV expression is a recurrent feature of epithelial malignancies, including colorectal carcinoma, where epigenetic deregulation and inflammatory signaling create permissive conditions for retroelement activation^39,40^. However, how the colorectal niche senses these induced viral elements -and why the resulting innate immune response fails to achieve viral clearance-remains a fundamental and unresolved question in CRC immune evasion.

Beyond NICA cells, we identify B lymphocyte, EBV Infected, Calamari Shaped (BLEICS) intermediate cells that may fuse with crypt cells and fibroblasts, driving clonal heterogeneity and stromal expansion. Human endogenous retrovirus (HERV) envelope proteins facilitate these fusion events and promote immune tolerance. BLEICS cells interact with early tumors and stimulate cancer associated fibroblasts via the CXCR4 SDF 1 axis, leading to stromal remodeling and crypt deformation.

In summary, we propose an integrative model of CRC pathogenesis: NICA cells, positioned at the intersection of neuronal, stem, and immune functions, undergo malignant transformation under pathogen infection and fusion events. BLEICS amplify stromal expansion through CAF (cancer-associated fibroblasts) interactions, while immune remodeling facilitates evasion. This framework redefines the cellular origins of CRC and highlights potential therapeutic opportunities in REST inhibition, EBV directed vaccination, and fusion blocking strategies.

## Results

### Argentaffin Cells in Colonic Epithelium: Unveiling Neuronal and Stemness Phenotypes

Enterochromaffin (argentaffin) cells have long been recognized in the gastrointestinal tract^41^. In the colon, they localize to crypts, lamina propria, and submucosa, forming part of the subepithelial neuroendocrine complex (**Figure 1A-I**). Within crypts, argentaffin cells typically display a balalaika shape with elongated extensions into the lumen, without specific spatial distribution.

**Figure 1:**
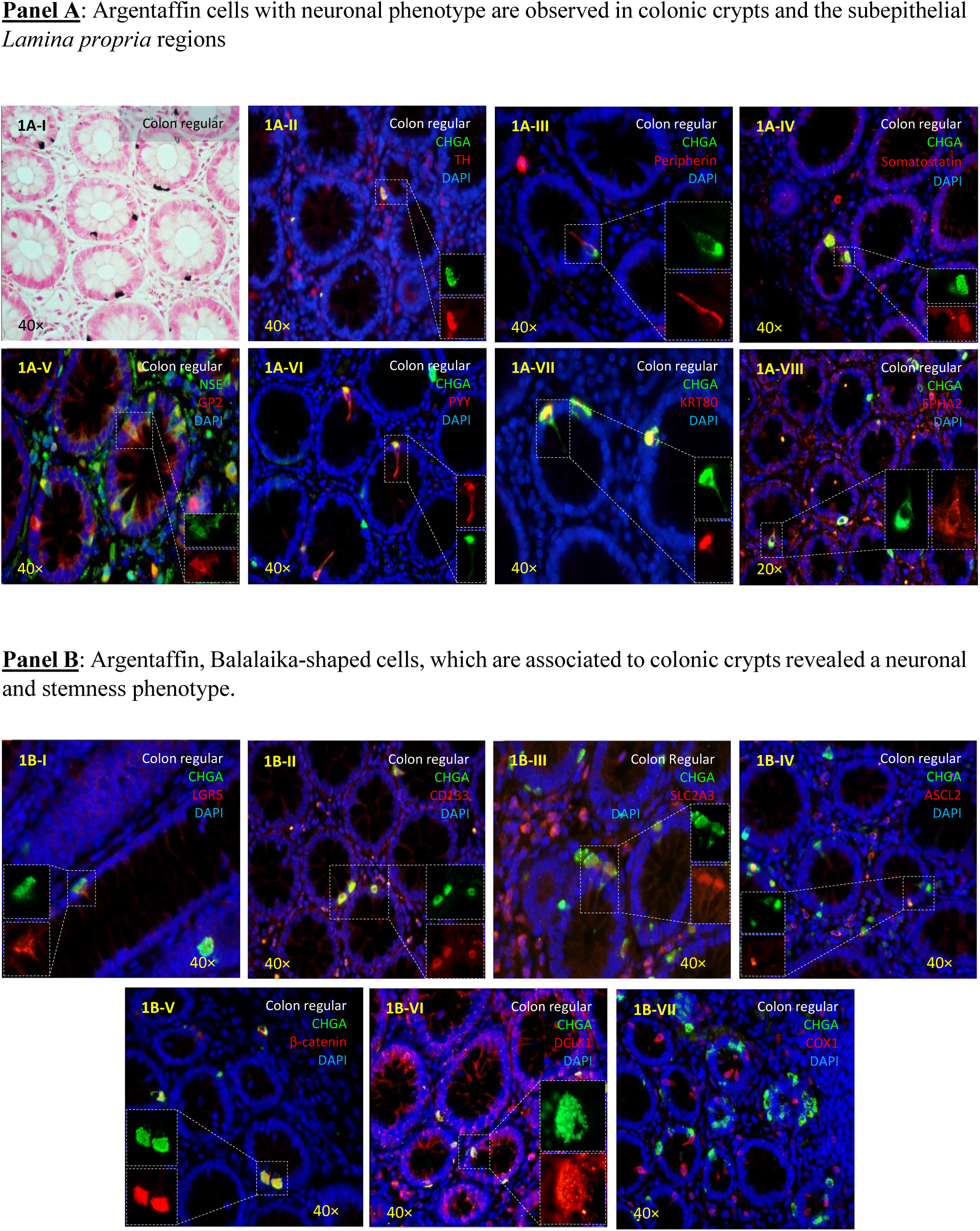
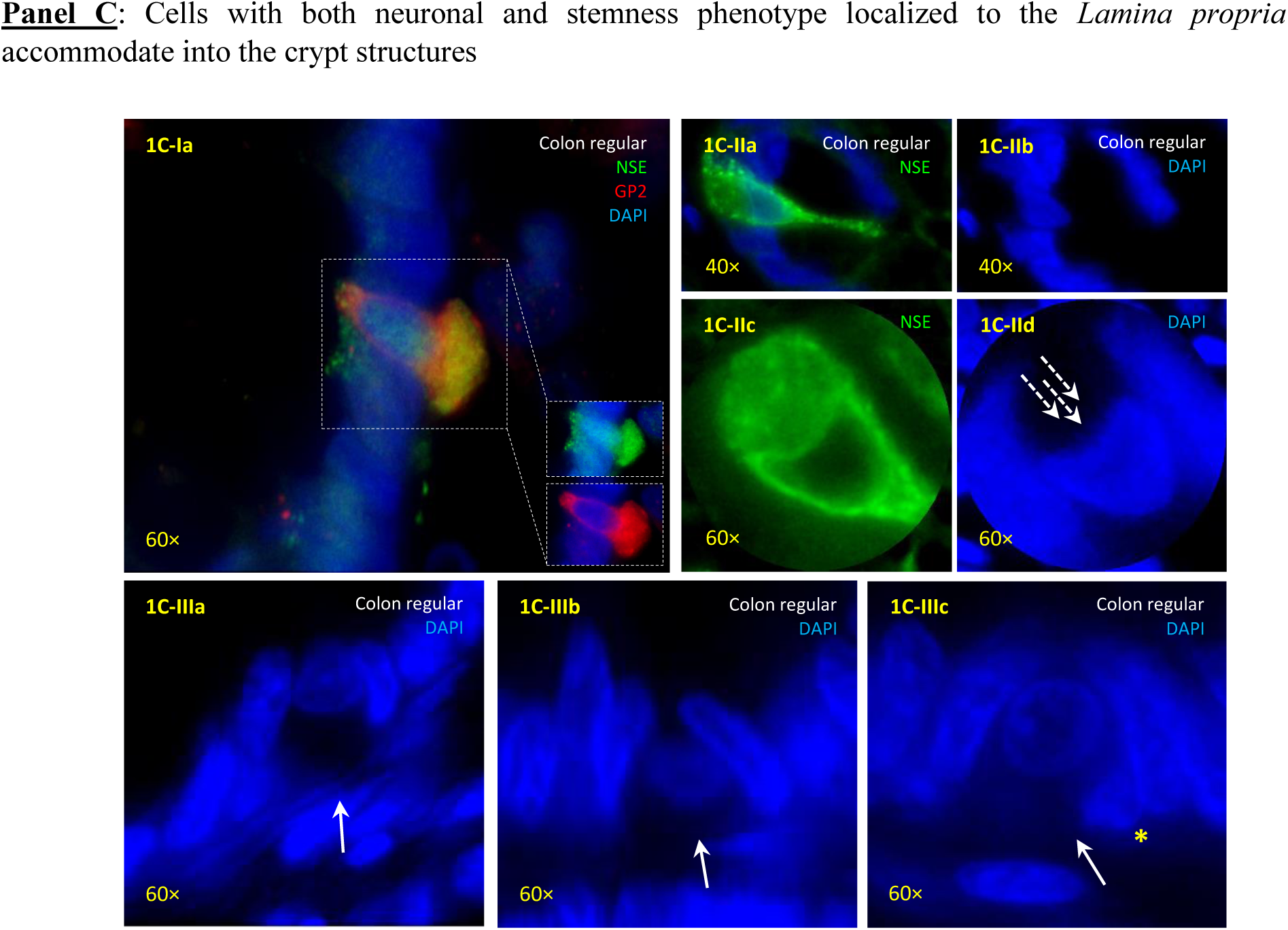
Characterization of argentaffin cells with stemness phenotype in human colonic tissue by IHC. **Panel A**: Argentaffin cells with neuronal phenotype are observed in crypts and subepithelial lamina propria, labeled by Fontana-Masson (1A-I). Crypt-localized Balalaika-shaped cells contain basal granules and co-express Chromogranin A (CHGA), Tyrosine hydroxylase (TH) (1A-II), Peripherin (1A-III), Somatostatin (1A-IV), Neuron-specific enolase (NSE) (1A-V), PYY (1A-VI), epithelial marker KRT80 (1A-VII), and Ephrin receptor EPHA2 (1A-VIII). **Panel B**: Balalaika-shaped cells show neuronal and stemness traits, expressing adult stem markers LGR5 (1B-I) and CD133 (1B-II), neuronal marker SLC2A3 (1B-III), transcription factor ASCL2 (1B-IV), Wnt pathway component β-catenin (1B-V), and the neuronal microtubule-associated kinase DCLK1 (1B-VI). They lack tuft cell marker COX1 (1B-VII). **Panel C**: CHGA-positive cells from lamina propria integrate into crypts (1C-Ia), with nuclei adopting hyperbolic shapes and reorganizing within crypt structures (1C-IIa-d). Some cells align along crypt bases (1C-IIIa-c), permitting docking at the basolateral pole (1C-IIIc, yellow asterisk; SI: NICA basolateral pocket). IHC was performed on paraffin sections of normal colon (>20 patients); experiments are representative.

These cells show neuronal features, expressing CHGA, TH (**Figure 1A-II**), Peripherin (**Figure 1A-III**), Somatostatin (**Figure 1A** I**V**), NSE (**Figure 1A-V**), PYY (**Figure 1A-VI**), the structural protein KRT80 (**Figure 1A-VII**), and EPH receptor A2 (EPHA2) (**Figure 1A-VIII**), which serves as an entry receptor for EBV and KSHV gamma herpesviruses^42^. Cells occur in two states: an “activated” CHGA form and a “quiescent” CHGA form, a duality also observed for other markers (data not shown).

Stemness analysis revealed expression of LGR5 (**Figure 1B**-**I**), CD133 (**Figure 1B-II**), GLUT3 (**Figure 1B-III**), ASCL2 (**Figure 1B-IV**), and β-catenin (**Figure 1B-V**), linked to stemness regulation and colorectal carcinoma biology^43^. DCLK1 marks stem-like cells in human tissues associated with progression and recurrence^44^. DCLK1 was detected in balalaika shaped epithelial cells, lamina propria cells (**Figure 1B-VI)**. Lymphatic follicle cells also stained positive (SI: DCLK1 in subepithelial colon). Notably, balalaika shaped cells lacked COX1 immunoreactivity (**Figure 1B-VII**), excluding them from being canonical human tuft cells, a cell type also considered a source of CRC^45,46^.

Cells residing in the lamina propria expressed markers similar to crypt located argentaffin cells, raising the possibility that balalaika shaped cells derive from LGR5 crypt stem cells or migrate from subepithelial compartments. Supporting this, neuronal phenotype cells were observed drilling into colonic crypts, with hyperbolic nuclear deformation during epithelial accommodation (**Figure 1C-Ia**, **IIa c**, **IIIa c**).

### Neuro-Immune Crypt-Associated (NICA) Cells: Bridging Innate and Adaptive Immune Systems

Following the characterization of the neuronal phenotype of argentaffin balalaika-shaped cells, we investigated their distribution in the colonic epithelium and proximal subepithelial compartments. **Figures 2A**-**Ia**-**c** illustrate their presence in the epithelial lining and their potential interaction with follicle-associated epithelium (FAE). Notably, a gradient of distribution was observed, with a greater concentration of balalaika-shaped cells near FAE, suggesting a possible functional link to subepithelial lymphoid structures (**Figure 2A**-**Ia**).

**Figure 2:**
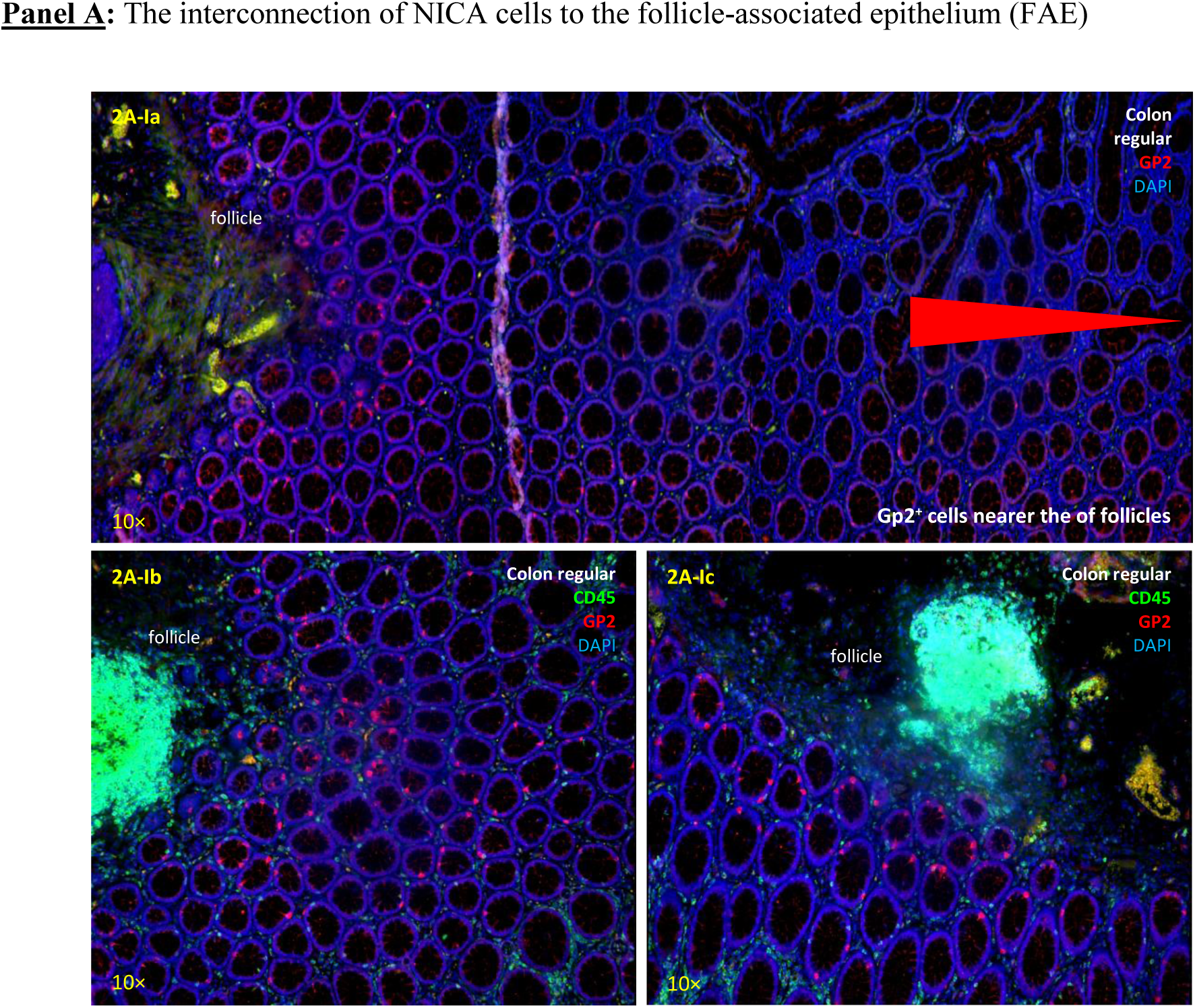
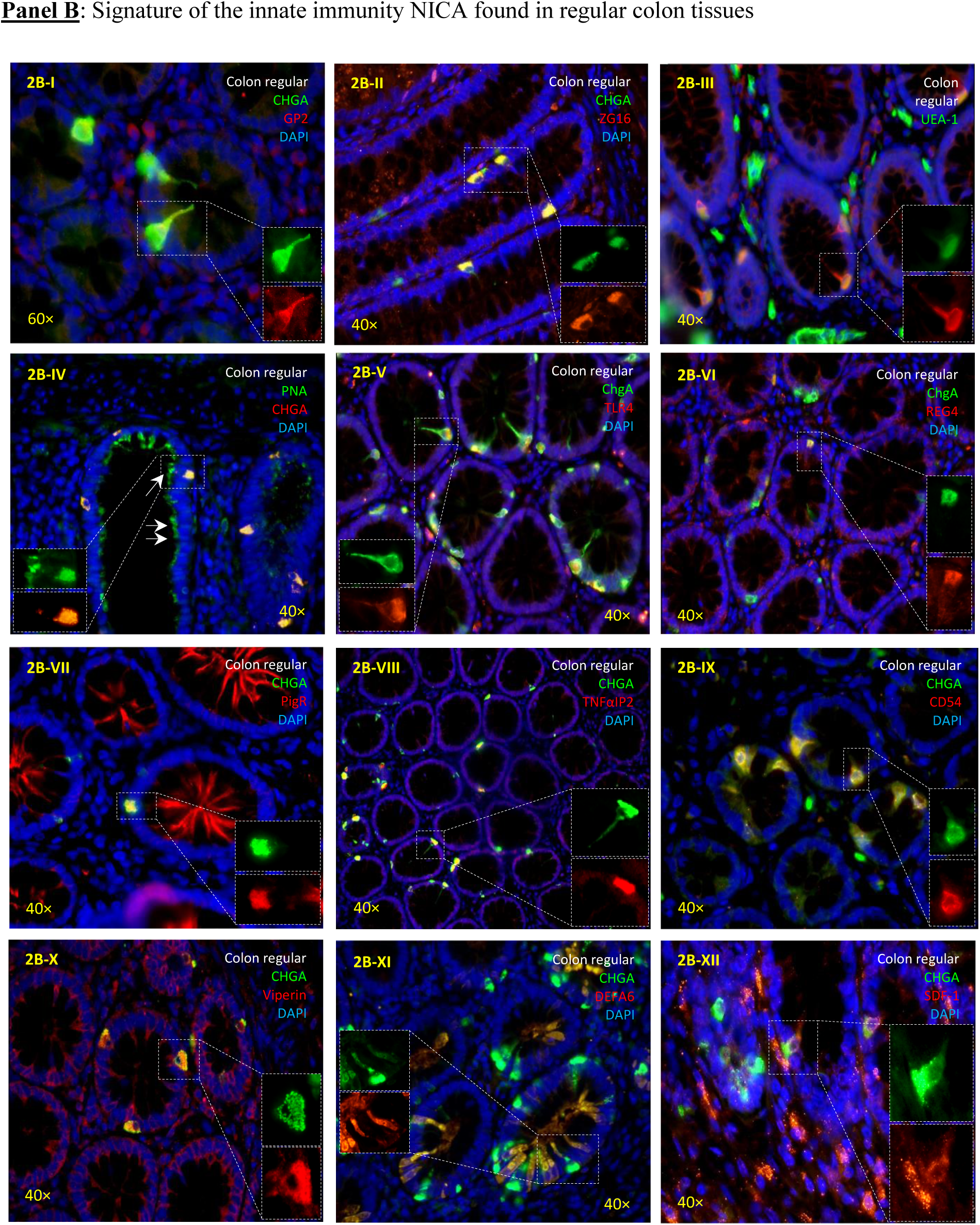
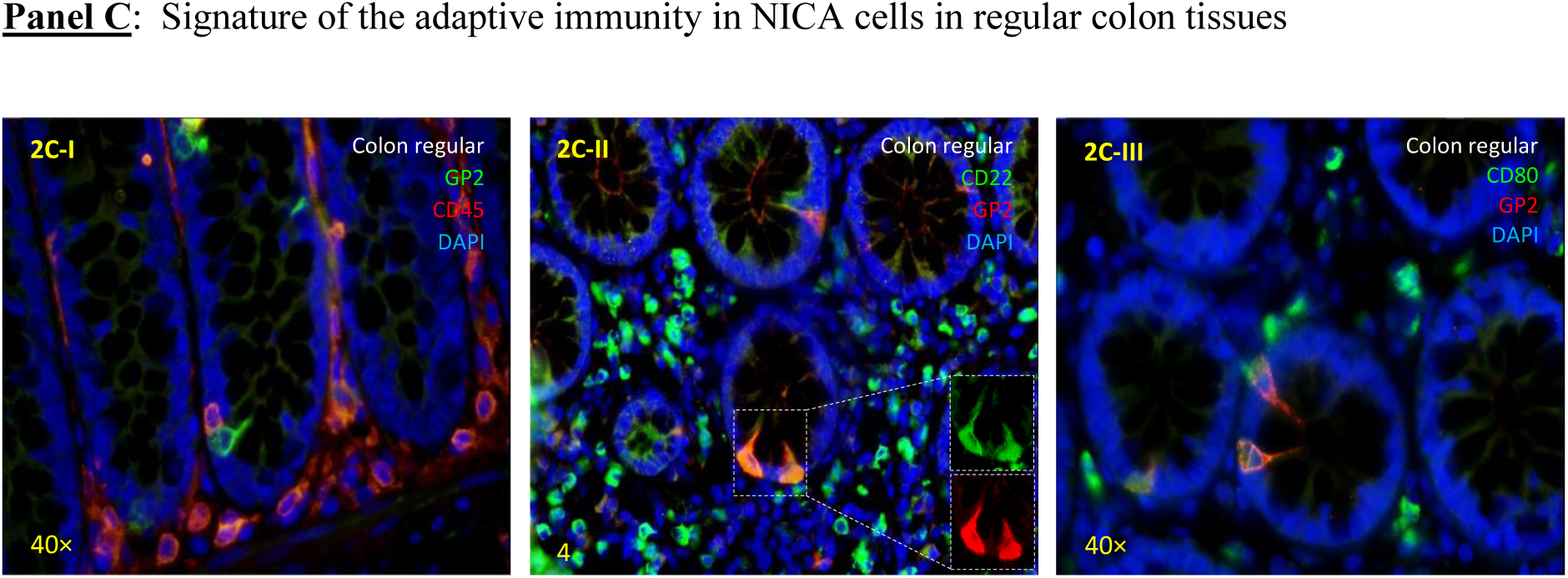
Balalaika-shaped Neuro-Immune Crypt-Associated (NICA) cells integrate innate and adaptive immune features, as demonstrated by immunohistochemical analysis. **Panel A**: Balalaika-shaped argentaffin cells are enriched around subepithelial lymphoid follicles in normal colon, forming a spatial gradient with highest density adjacent to them (2A-Ia-Ic). **Panel B**: NICA cells show innate immune signatures. They express the zymogen-granule–associated secretory proteins GP2 (2B-I) and ZG16 (2B-II), bind FITC-lectins UEA-1 (2B-III), PNA (2B-IV) and express the pattern-recognition receptor TLR4 (2B-V). Inflammatory proteins like REG4 (2B-VI), PigR (2B-VII), TNFaIP2 (2B-VIII), as well as the adhesion molecule CD54 (2B-IX) are detected. NICA cells additionally express antiviral viperin (2B-X), antibacterial DEFA6 (2B-XI), and the chemokine SDF-1/CXCL12 (2B-XII). **Panel C**: Although NICA cells exhibit adaptive immune features, they do not express the pan-leukocyte marker CD45, indicating a non-hematopoietic origin (2C-I). They express CD22 (2C-II) and CD80 (2C-III), markers typically associated with B cells and antigen-presenting cells. Additional viral entry receptors, including CAR (SI: CAR and E1B-55K localization in NICA and colon tissues), were detected in NICA cells. IHC was performed on paraffin sections of normal colon (>20 patients); figures are representative.

### Immune Features of Balalaika-Shaped Cells

An early indication of immunological properties was expression of Glycoprotein 2 (GP2), a bacterial transcytotic receptor typical of M-cells. This suggested a specialized M cell variant. Supporting this, some cells formed basolateral pockets housing immune cells (SI: NICA basolateral pocket). However, the neuronal phenotype exhibited by these cells ruled out their identity as M-cells. Unlike M-cells, GP2 expression extended throughout the cell body, and a soluble form is secreted into the crypt lumen, forming a protective epithelial layer (**Figure 2B-I**). Additional immune markers included besides GP2^47^, the Zymogen granule protein 16 (ZG16), which binds pathogens and prevents barrier penetration^48^ (**Figure 2B-II**). Balalaika shaped cells show weak labeling with Ulex europaeus agglutinin 1 (UEA 1) (**Figure 2B-III**) but display strong reactivity with peanut agglutinin (PNA), which specifically binds the oncofetal Thomsen Friedenreich antigen^49^ (**Figure 2B-IIa**/**IIb**). Balalaika shaped cells expressed Toll-Like Receptor 4 (TLR4) (**Figure 2B-V**)^50^, REG4 (**Figure 2B-VI**)^51^, PigR (**Figure 2B-VII**)^52^, TNFαIP2 (**Figure 2B-VIII**)^53,54^, and ICAM 1 (CD54) (**Figure 2B-IX**), a viral receptor for rhinoviruses and influenza^55^, which presence suggests crypt associated viral entry points analogous to M-cells in Peyer’s patches.

Two potent antimicrobial effectors, Viperin (**Figure 2B-X**)^56^ and DEFA6 (**Figure 2B-XI**)^57^, were expressed, along with stromal cell derived factor 1 (SDF 1/CXCL12)^58^ (**Figure 2B-XII**). Bioactive peptides derived from chromogranin A (CHGA) also exert antipathogen activity^59^. Collectively, lectins, antimicrobial peptides, innate receptors, and immunomodulators define balalaika shaped cells as a distinct neuroimmune effector population.

Based on their neuroimmune unique marker combinations and conserved morphology, we designated the Balalaika shaped cells as Neuro Immune Crypt Associated (NICA) cells.

### Adaptive Immune Properties of NICA Cells

To assess adaptive features, we examined key markers. CD45, a leukocyte common antigen, was absent in NICA cells (**Figure 2C-I**), though CD45 immune cells were present within crypts. Despite lacking CD45, NICA cells expressed CD22 (**Figure 2C-II**), typically found on B lymphocytes and APCs^60^. CD80 (**Figure 2C-III**), a co stimulatory molecule induced by microbial infections^61^ and parasites^62^, was also detected. CD80 serves as a receptor for clade B adenoviruses^63^, some with oncogenic potential^64^.

The adenovirus type 5 receptor CAR was expressed in crypt epithelium, lymphatic follicle immune cells, and lamina propria patrol cells, with polarized basolateral distribution^65^. Notably, CHGA NICA cells showed strong basolateral CAR expression, exceeding surrounding crypt epithelium, which. This enrichment positions NICA cells as potential viral entry points for adenovirus type 5. Importantly, adenovirus type 5 can hijack the p53 pathway, disrupting cell cycle control and conferring malignant potential, underscoring the relevance of CAR^+^ NICA cells in early colorectal carcinogenesis. Moreover, the adenoviral protein E1B-55K is detected in a mosaic distribution within colonic tissues, specifically localizing to a subset of CHGA cells that are morphologically indistinguishable from NICA cells (SI: CAR and E1B-55K localization in NICA and colon tissues).

### Colon Carcinoma Progression Leads to a Loss of Neuronal Identity

A transcriptome covering all 18 566 genes was analyzed in the study, revealing a “Big Picture” view where the tumor crypt transformation is characterized by more widespread downregulation and loss of identity than the gain of new features. With 338 genes significantly increasing (FC > 2, p < 0.05) and 898 genes significantly decreasing (FC < 0.5, p < 0.05), the tumor crypt effectively loses nearly three times more functional features than it gains, while 17 330 genes remain relatively stable. Globally, the transcriptome reveals that tumor crypt transformation is defined by a statistically robust liquidation of the native neuroendocrine identity.

Transcriptomic analysis revealed a loss of neuronal identity in tumor crypts compared to regular crypts. Gene Set Enrichment Analysis (GSEA) demonstrated a negative enrichment of multiple neuronal-related pathways in tumor crypts (NES < 0, p < 0.05). (**Figure 3A**-**I**). Differential expression analysis showed a significant reduction (≈72%) in neuronal gene expression in tumor crypts versus regular crypts (**Figure 3A**-**II)**. A similar pattern was observed when comparing primary and commercial colon cancer cell cultures with normal crypts (**Figure 3A**-**III** & SI: Neuronal genes in crypts), indicating the suppression of the neuronal phenotype in tumor cells.

**Figure 3:**
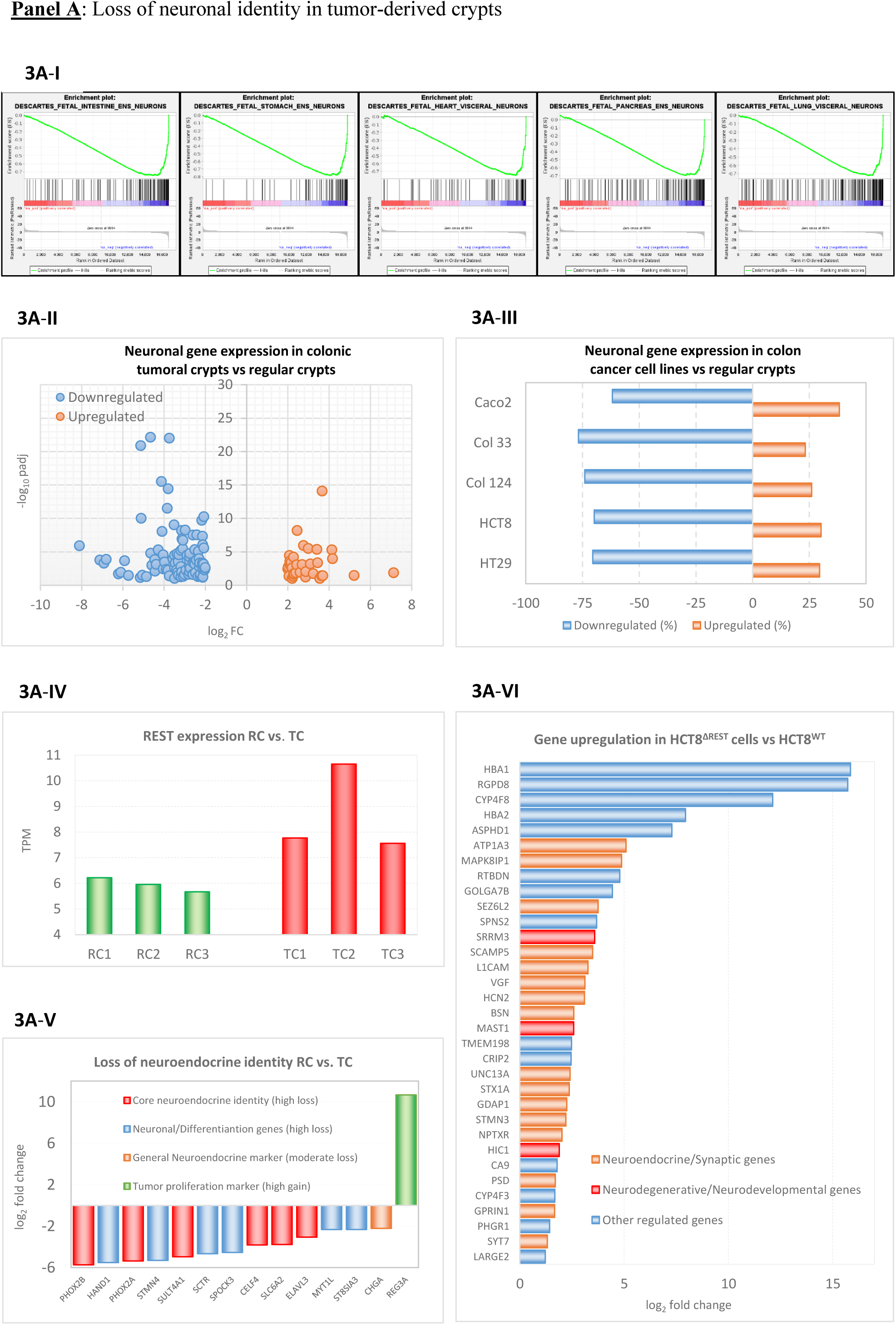
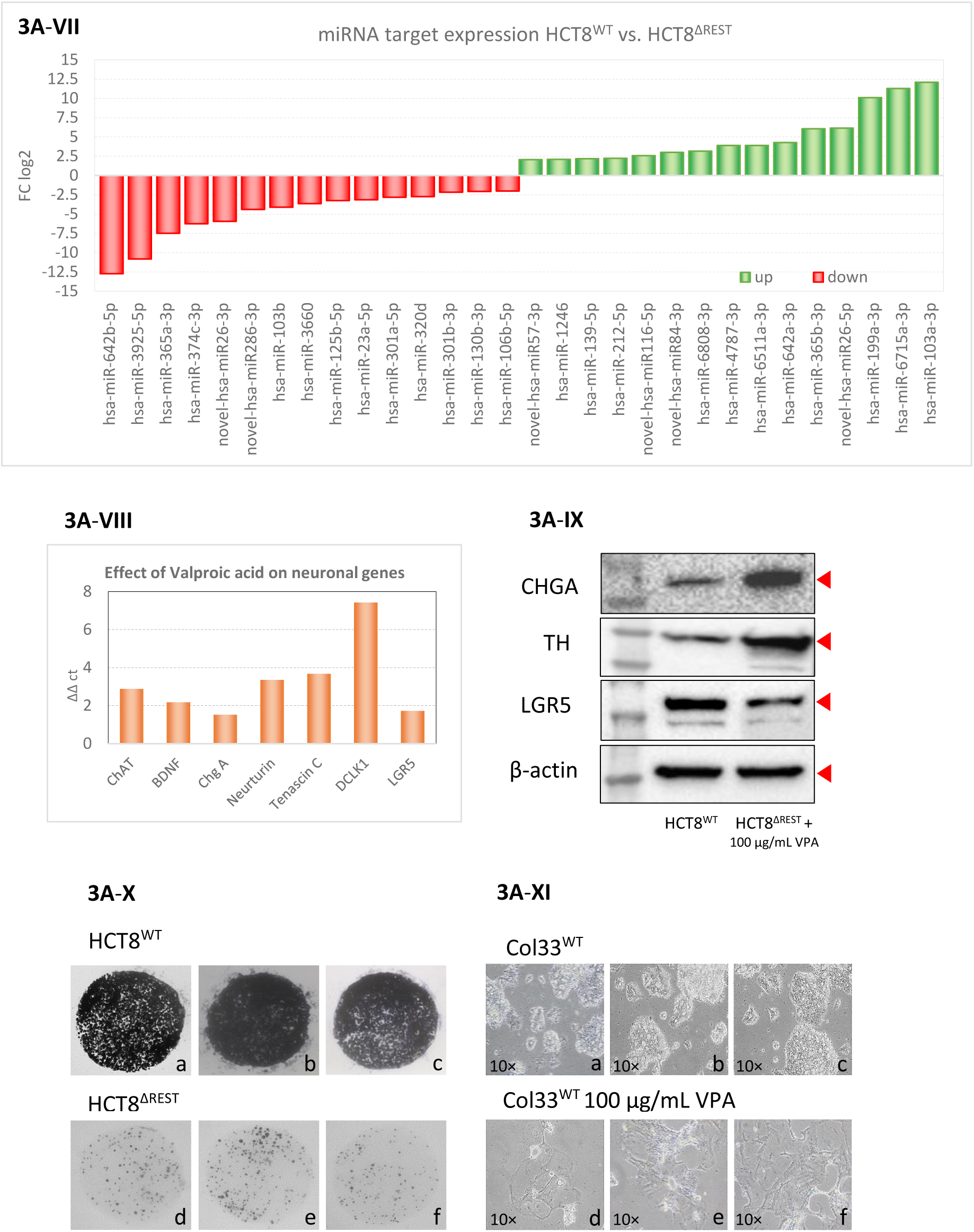
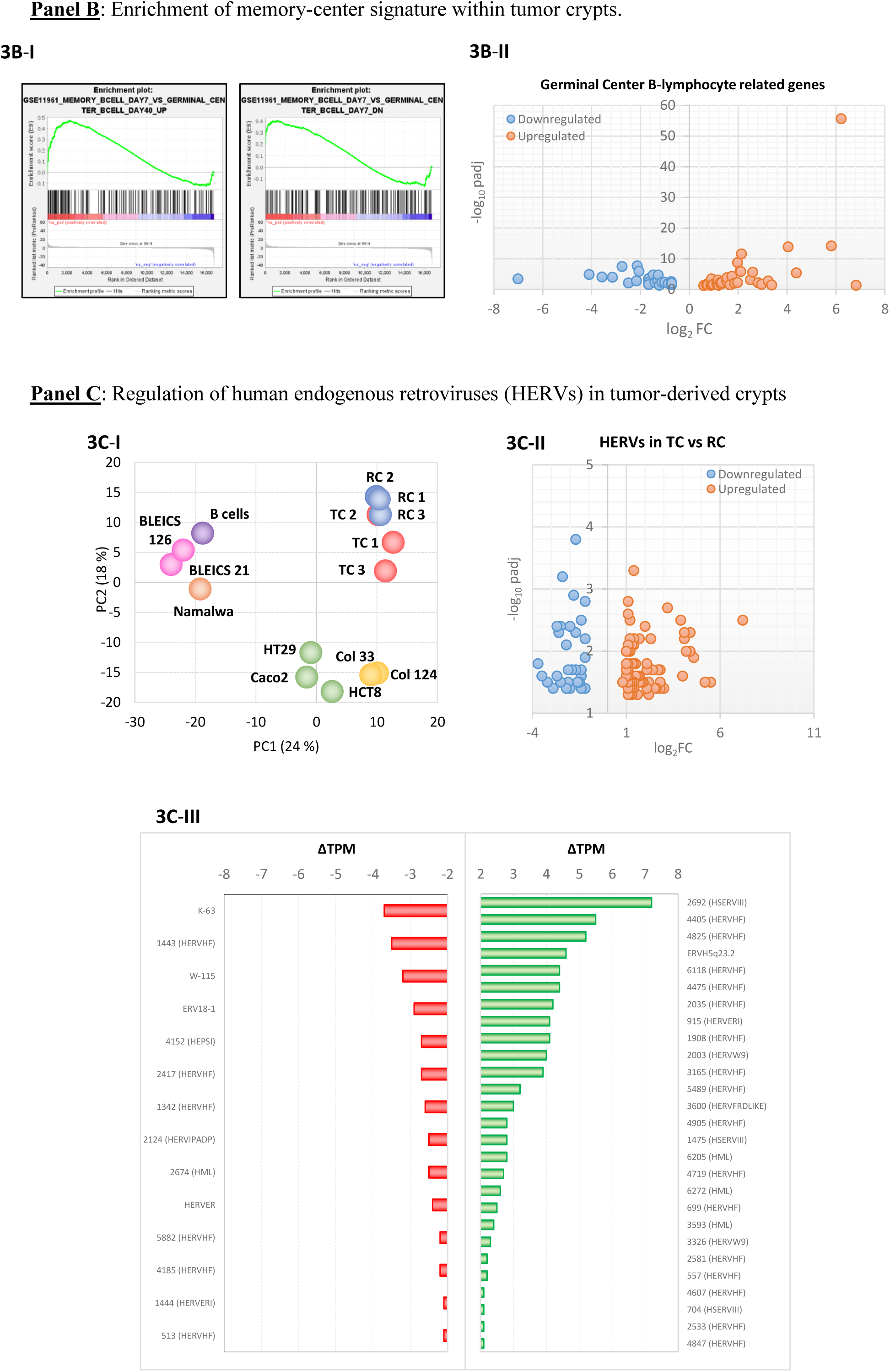
Panel. **A**: GSEA indicates strong negative enrichment of neuronal pathways in tumor crypts (TC) vs. regular crypts (RC) (3A-I), confirmed by DEG analysis (NES < 0, p < 0.05). Intersected DEGs highlight downregulated neuronal genes (3A-II: *CHGA*, *PHOX2B*, *SLC6A2*, *GABRG2*, *SNAP25*, *NRXN1*, *RELN*, *GAP43*, *NEFM*, *NLGN1*) (|log₂ FC| > 2 & padj < 0.05). Same patter were detected in CRC cells (Geneset: R-HSA-112316, |log₂ FC| > 2; 3A-III). *REST* is upregulated in tumor crypts (3A-IV). Tumor crypts revealed loss of neuroendocrine identity (3A-V) by downregulation of PHOX2B (log₂ FC ≈ -6.90), *PHOX2A* (-7.91), *SLC6A2*, and *CHGA* (-2.27) with an induction of *REG3A* (log₂ FC ≈ +9.10). REST loss in HCT8^ΔREST^ cells drives derepression of neuroendocrine/synaptic genes (3A-VI: *STX1A*, *SYT7*, *UNC13A*, *VGF*, *ATP1A3*, *BSN*, *L1CAM*, *SCAMP5*, *SEZ6L2*, *STMN3*, *HCN2*, *GPRIN1*, *GDAP1*, *MAPK8IP1*, *NPTXR*, and *PSD*); neurodevelopmental/degenerative genes (3A-VI: *ASPHD1*, *SRRM3*, *MAST1*, *RTBDN*); and collateral stress genes (3A-VI: *HBA1*, *HBA2*, *RGPD8*, *CYP4F8*, *CA9*, *CRIP2*, *TMEM198*, *LARGE2*, *PHGR1*, *SPNS2*, *HIC1*, *GOLGA7B*, *CYP4F3*). HCT8^ΔREST^ cells exhibit deregulated miRNAs that drive a shift toward neuroendocrine identity (3A-VII), converging on glutamatergic (miR-124, miR-181, miR-375), cholinergic (miR-212, miR-215), Oxytocin (miR-1468, miR-365b), thyroid (miR-195, miR-642, miR-651), and Alzheimer-associated modules (miR-103, miR-139, miR-447, novel-hsa-miR131-3p). Valproic acid (VPA, 30 µg/ml) induces neuronal genes (*CHAT*, *CHGA*, *BDNF*, *NRTN*, *TNC* and stemness markers *DCLK1*, *LGR5* measured by qPCR (3A-VIII). Western blots revealed an overexpression of CHGA and TH in HCT8^ΔREST^ cells incubated with VPA (100 µg/ml, 72 h), accompanied by a reduction in LGR5 (3A-IX). Consistent with this shift, HCT8^ΔREST^ cells show a marked impairment in organoid-forming capacity (3A-Xa-f). Continuous VPA (100 µg/ml, 14d) arrests growth and induces hypertrophic/senescent morphology (3A-XI). **Panel B**: GSEA reveals germinal center B-cell pathway enrichment in tumor crypts (3B-I); DEGs intersected with B-cell gene set (|log₂ FC| > 2 & p < 0.05; 3B-II). **Panel C**: PCA of HERV expression separates TC, RC, and CRC cells (3C-I). DEG analysis between TC and RC identified 111 DE-HERVs (78 up & 33 down; |log₂ FC| > 2 & p < 0.05; 3C-II). Differentially expressed HERVs are shown as bar graphs (3C-III, SI: Table 1 - Differentially expressed HERVs within colorectal crypts presenting ISD). Bars represent ΔTPM values. Statistical significance was defined as p < 0.05 (Benjamini-Hochberg correction).

### Atavistic Priming and Lineage Destabilization in Colorectal Tumors

Colorectal tumor crypts display a profound transcriptional reorganization characterized by activation of an early neuronal-like program without progression toward terminal neuronal or neuroendocrine differentiation. This state is defined by induction of early neurogenic and stress responsive regulators, including *DLX6*, *OLIG2*, *EGR3*, and immediate-early genes such as *FOS* and *EGR1*, together with a marked surge of primordial neuronal determinants *SOX1* and *NKX2*-*1*. Despite this initiation of a neurogenic trajectory, the differentiation program collapses at its inception: essential maturation factors (*POU3F3*, *PAX4*, *PAX6*, *NEUROD*1/2, and *KLF9*) are transcriptionally extinguished across all examined systems, including primary cultures and immortalized colorectal cancer lines. This early-activation/terminal-arrest configuration is reinforced by persistent expression of REST, some ZNFs [including *ZNF*598 (log_2_FC =+0.55, ≈1.46 fold), and *ZNF*146 (log_2_FC =+0.78, 1.72 fold)], as well as the *RCOR*1/3 complex, which impose a stable repressive architecture that blocks neuronal differentiation and destabilizes epithelial maturation programs. The resulting phenotype constitutes a conflicted neuro-epithelial state in which cells initiate an ancestral neuronal program yet remain structurally incapable of completing lineage specification. Transcriptomic analysis across seven independent models (TC1, TC3, Col33, Col124, HCT8, HT29, and Caco2) reveals a conserved two tiered regulatory architecture: a Fixed Core defined by extinction of maturation determinants and stable maintenance of the REST-ZNFs-SOX4 axis, and a Dynamic Surge in which SOX1 and NKX2-1 are strongly induced in the tumor niche but suppressed *in vitro* (SI: Neuronal transcription factors). This architecture demonstrates that colorectal cancer cells exist in a state of atavistic priming, in which the machinery for an ancient neuronal program is activated but permanently prevented from reaching completion, establishing a stable, lineage-disrupted identity that persists across biological contexts^66,67^.

### REST Inactivation Reveals Neuronal Cell Identity in Colorectal Carcinoma Models

The transition from a neuronal to a tumoral phenotype is governed by genetic and epigenetic mechanisms, with the transcription factor REST, probably acting as a proposed lineage gatekeeper. Transcriptomic analysis showed REST upregulation in tumor crypts compared to normal crypts (Mean TC ≈+8.7 TPM vs. Mean RC ≈+6.0 TPM, FC ≈+1.5). (**Figure 3**, **3A**-**IV**). These results were subsequently confirmed by quantitative qPCR (Fold Change 2^-ΔΔCt^ ∼1.52). Moreover, the transcriptome analysis revealed that ZNF800 -a transcription factor (TF) known to enforce the REST pathway-was upregulated in tumor crypts, a finding subsequently corroborated by qPCR (Fold Change 2^-ΔΔCt^ ∼2.83). Furthermore, approximately 76% of REST-dependent neuronal genes identified in tumor crypts were similarly downregulated in a publicly available metastatic small-cell lung cancer dataset (GSE40695 CREEDSID GENE) following REST inhibition, indicating that tumor crypts suppress neuronal gene expression via a REST-dependent mechanism during malignant progression. Robust differential expression analysis confirmed that tumor crypts exhibit an active repression of the neuroendocrine lineage and induction of regenerative transcripts (REG3A) (**Figure 3A**-**V**). A strong de-differentiation signature was evidenced by the near-complete extinction (over 90% reduction) of critical neuroendocrine genes. Master regulatory transcription factors-like *PHOX2A* (log_2_ FC ≈-7.91, p < 0.001), *PHOX2B* (log_2_ FC ≈-9.10, p < 0.001), *NEUROD1* (log_2_ FC ≈-1.38, p=0.036) and *PAX4* (log2 FC ≈-1.63, p=0.003) were among the most severely suppressed, which was corroborated by concomitant downregulation of neurotransmitter machinery (SLC6A2, log_2_ FC ≈-7.22, p < 0.001) and the pan-neuroendocrine biomarker *CHGA* (log_2_ FC ≈-2.27, p < 0.01).

Conversely, the transcriptional profile of HCT8^ΔREST^ (REST-null) cells showed widespread derepression, most pronounced in the neuroendocrine/synaptic program (**Figure 3A**-**VI**). This core group of 16 genes exhibited consistent upregulation (log_2_ FC range: +1.3 to +7.3), activating key components of neurotransmission (e.g., *UNC13A*, *STX1A*, *BSN*) and neuroendocrine signaling (*VGF*, *ATP1A3*). qPCR analysis confirmed a marked induction of *CHGA* in HCT8^ΔREST^ cells. REST deficient cells showed a ΔCt of 29.69 ± 0.39 compared with 34.19 ± 1.16 in parental HCT8, yielding a ΔΔCt of 4.5, corresponding to ≈+23 fold higher *CHGA* expression (p=0.001). These data validate *CHGA* as a robust REST repressed target in this system.

REST ablation also induced several non-canonical targets, underscoring its broad role in maintaining epithelial identity (**Figure 3A**-**VI**). The strongest inductions were *HBA1* (log_2_ FC ≈+15.8) and *HBA2* (log_2_ FC ≈+9.7). Ectopic activation of these hemoglobin subunits may reflect metabolic stress adaptation and a shift toward the energy-intensive secretory activity characteristic of neuronal/neuroendocrine cells. Furthermore, *RGPD8* (log_2_ FC ≈+12.5), robustly derepressed and possessing Ran-binding domains, suggests a role in nuclear transport specialization that reinforces dynamic gene regulation essential for the neuroendocrine phenotype. *CYP4F8* (log_2_ FC ≈+11.9), which modulates lipid mediator turnover, suggests that REST normally restrains oxygenase-driven metabolic programs, and its loss unleashes atypical lipid signaling modules. Finally, *ASPHD1* (log_2_ FC ≈+7.3), normally enriched in neuronal tissues, was derepressed, marking activation of a high-energy secretory program consistent with a neuroendocrine switch. The strong induction and relevant function of these atypical targets position them as candidate markers of neuroendocrine lineage identity in this model, confirming REST’s broad repressive scope and its role in destabilizing lineage fidelity.

### Loss of REST Induces a Neuroendocrine miRNA Program

Small RNA sequencing of HCT8^ΔREST^ versus HCT8^WT^ cells revealed a significant reconfiguration of the regulatory landscape (FDR < 0.05), driving a shift toward neuroendocrine identity. Using a threshold of Log_2_ FC > 1.5, the NE program was robustly defined by the upregulation of canonical NE regulators including miR-375-3p (≈+1.94), miR-365b-3p (≈+6.06), miR-212-5p (≈+2.24), and miR-215-5p (≈+1.62). Pathway enrichment confirmed convergence on neuronal modules (glutamatergic, cholinergic) and REST-linked Alzheimer’s disease pathways (**Figure 3A-VII**).

### REST Dependent Tumor Suppression Axis

Among deregulated miRNAs, miR 139 5p (+2.18) emerged as a dominant tumor suppressor. Its induction in HCT8^ΔREST^ cells supports a model in which REST represses miR 139 5p transcription in the wildtype context, contributing to malignancy. Derepression enables inhibition of oncogenic targets such as NOTCH1 and AMFR, providing a mechanistic basis for impaired colorectal cancer organoid growth (**Figure 3A-VII**).

### The Fate of the Malignancy: Regulatory Failure

The HCT8^ΔREST^ transcriptome confirms a profound failure of the HCT8 malignant program, resulting in impaired growth. The dominant cause is the simultaneous activation of the miR-139-5p tumor brake and a toxic neuroendocrine (NE) lineage switch (miR-375-3p, miR-365b-3p)^68,69^. This dual anti-malignancy signal is so potent that the tumor cannot maintain its proliferative epithelial identity. Molecularly, while the loss of tumor-suppressive miRNAs leads to the weak upregulation (Log_2_ FC ≈+0.1 to +0.4) of key oncogenes (*ADAM10*, *HELLS*, *STAT3*, and *MYC*), this effect is merely background noise (SI: miRNA in HCT8 REST-null). The failure to significantly upregulate *CDK4* and *CTNNB1* confirms the marked impairment of proliferation and core WNT signaling. The cells’ biological sensor detects the catastrophic loss of its epithelial identity and its WNT/NOTCH-driven proliferation, but the molecular machinery available to compensate -such as the release of *ADAM10*- is insufficient to overcome the dominant, cytotoxic lineage commitment (**Figure 3A-VII**).

### REST as Gatekeeper of Transcription-Translation Congruence

The HCT8^ΔREST^ phenotype restores protein levels congruent with RNA abundance because the loss of REST destroys its function as an indirect gatekeeper of translation. This regulatory collapse is marked by the catastrophic downregulation of numerous tumor-suppressive miRNAs (e.g.: hsa-miR-642b-5p, hsa-miR-3925-5p) that previously acted as a post-transcriptional gate, binding to and actively inhibiting the translation of NE effector mRNAs^70^. By removing this miRNA brake, REST-null cells are unable to restrict protein synthesis, resulting in protein levels that are now directly proportional to RNA abundance and contributing to the irreversible and ultimately cytotoxic commitment to the neuroendocrine lineage (**Figure 3A-VII**).

### Synergistic Reprogramming and Phenotypic Switch

In HCT8^WT^ cells, pharmacological inhibition of REST using VPA (30 µg/mL, 72 h) induces robust upregulation of the neuronal genes *CHAT*, *BDNF*, *CHGA*, Neurturin (*NRTN*), and Tenascin C (*TNC*), as well as the stemness associated markers *DCLK1* and *LGR5*, as measured by qPCR (**Figure 3A**-**VIII**). Protein validation confirms commitment to a neuroendocrine fate: the secretory markers CHGA and TH are strongly induced, whereas the stemness factor LGR5 is markedly reduced (**Figure 3A**-**IX**). This upregulation is further potentiated by VPA treatment (100 µg/mL, 72h), highlighting a synergistic epigenetic mechanism that reinforces NE differentiation (**Figure 3A**-**IX**). HCT8^WT^ cells efficiently form organoids under 3D neuronal medium conditions (**Figure 3A**-**IXa**-**c**), in contrast to HCT8^ΔREST^ cells, which display a substantial loss of organoid forming capacity (**Figure 3A**-**Xd**-**f**). This switch, accompanied by LGR5 downregulation, suggests that REST activity is required to sustain stemness under these conditions. In line with this, extended VPA treatment (100 µg/mL for two weeks) markedly reduces proliferation in primary CRC cultures, a response consistent with senescence induction (**Figure 3A**-**XI**).

### Immune Remodeling and Germinal Center Signature

Transcriptome analysis of tumor crypts revealed a loss of neuroendocrine identity but acquired immune-related gene expression (SI: Immune significant genes TC). Gene set enrichment analysis of tumor crypts vs. regular crypts, revealed a significant enrichment of transcripts associated with late-stage germinal center B cells, indicating that tumor crypts undergo immune remodeling characterized by ectopic GC-like activation (GSE11991 signature, **Figures 3B-I**). Volcano plot analysis (**Figure 3B-II**) revealed significant upregulation of germinal center -associated genes in tumor crypts, consistent with the emergence of a GC-like transcriptional program. These genes, typically involved in B-cell activation, cytokine signaling, and immune niche formation, suggest that tumor crypts undergo immune remodeling toward a germinal center-like phenotype (**Figure 3B-II**). This immune-like reprogramming is compatible with a fusion linked origin, in which epithelial tumor cells inherit B cell-derived transcriptional elements from EBV conditioned intermediates.

Importantly, CIBERSORTx deconvolution showed that tumor crypts lose plasma cell signatures (p=0.05) while gaining M1 macrophage-like programs (p=0.02), despite extensive washing and minimal immune cell presence. Together, these findings support a model in which tumor crypts acquire stable, cell intrinsic immune transcriptional programs through immune-epithelial interactions and fusion events, contributing to immune evasion and microenvironmental remodeling. (SI: CIBERSORTx Results).

### HERV Gene Segments with Immunosuppressive and Fusogenic Potential

ERVmap analysis revealed cell type specific transcriptional regulation of human endogenous retroviruses (HERVs). Principal component analysis (PCA) showed that tumor crypts display greater transcriptional dispersion than regular crypts, consistent with ongoing diversification during tumor evolution. Isolated colorectal cancer (CRC) primary cells and commercial cell lines clustered distantly from regular crypts (**Figure 3-C I**). Although BLEICS originate from B cells, they segregate from primary B cells due to their immortalized, culture adapted transcriptional state.

Volcano plot analysis (**Figure 3C-II**) identified 111 differentially expressed HERV loci in tumor crypts (78 upregulated, 33 downregulated; **Figure 3C-III**). Many DE HERVs retain partially conserved gag (CA, NC, and MA) and pol (RT, IN) segments, indicating preservation of subgenomic structure (SI: HERV transcriptome data TC). Upregulated loci were enriched for intact immunosuppressive domains (ISDs), whereas downregulated loci largely lacked ISD conservation^71^ (SI: Table 1: Differentially expressed HERVs within colorectal crypts presenting ISD). This pattern suggests that tumor crypts preferentially activate evolutionarily recent, ISD competent HERVs with preserved immunomodulatory potential.

Family specific analysis showed selective derepression of HERVH and HERVW, retroviral families associated with stemness, immune mimicry, and fusogenic signaling. These findings support a model in which retroelement activation contributes to epithelial-immune remodeling within the tumor microenvironment.

Because HERVs can be transactivated by exogenous viral proteins, their re expression in cancer has translational relevance. Introducing mRNA encoding viral structural proteins such as Ad5 E1B 55K or MCV LTAg into p53 mutated CRC cells was sufficient to induce HERVs with intact ISD or fusogenic properties (SI: HERV Transactivation by Viral Structural Proteins). This positions HERV activation as a potential biomarker of viral cofactor activity and a candidate target for immunotherapeutic intervention.

A focused analysis of HERV envelope fragments revealed that upregulated DE HERVs frequently contained fusogenic surface (SU) and transmembrane (TM) domains (41 TM, 25 SU, 19 both), whereas these domains were less common among downregulated loci (14 TM, 6 SU, 3 both). ISD containing loci also showed preferential upregulation (16 up vs. 5 down). Together, these features indicate that transcriptionally active HERVs in tumor crypts preferentially preserve immunosuppressive and fusogenic capacities, potentially shaping local immune dynamics and epithelial plasticity.

### NICA Cell Proliferation during the Transition from Carcinoid to Adenocarcinoma

Histopathological analysis of early-stage CRCs revealed a localized proliferation of CHGA^+^ NICA cells near the FAE (**Figures 4A**-**I**-**VIII**). The CHGA^+^ cells are mainly confined to the epithelial layer. The declining abundance of these cells over time suggests a transition from carcinoid-like tumors to the predominant adenocarcinoma classification. In the colonic crypts, LBP (Lipopolysaccharide Binding Protein) is detected as a sparse signal localized within the cytoplasm of epithelial cells, showing a distinct lack of co-expression with CHGA^+^ neuroendocrine cells. This spatial separation indicates that the innate immune sensing of LBP and the neuroendocrine program of CHGA are managed by independent cellular lineages within the crypt architecture.

**Figure 4:**
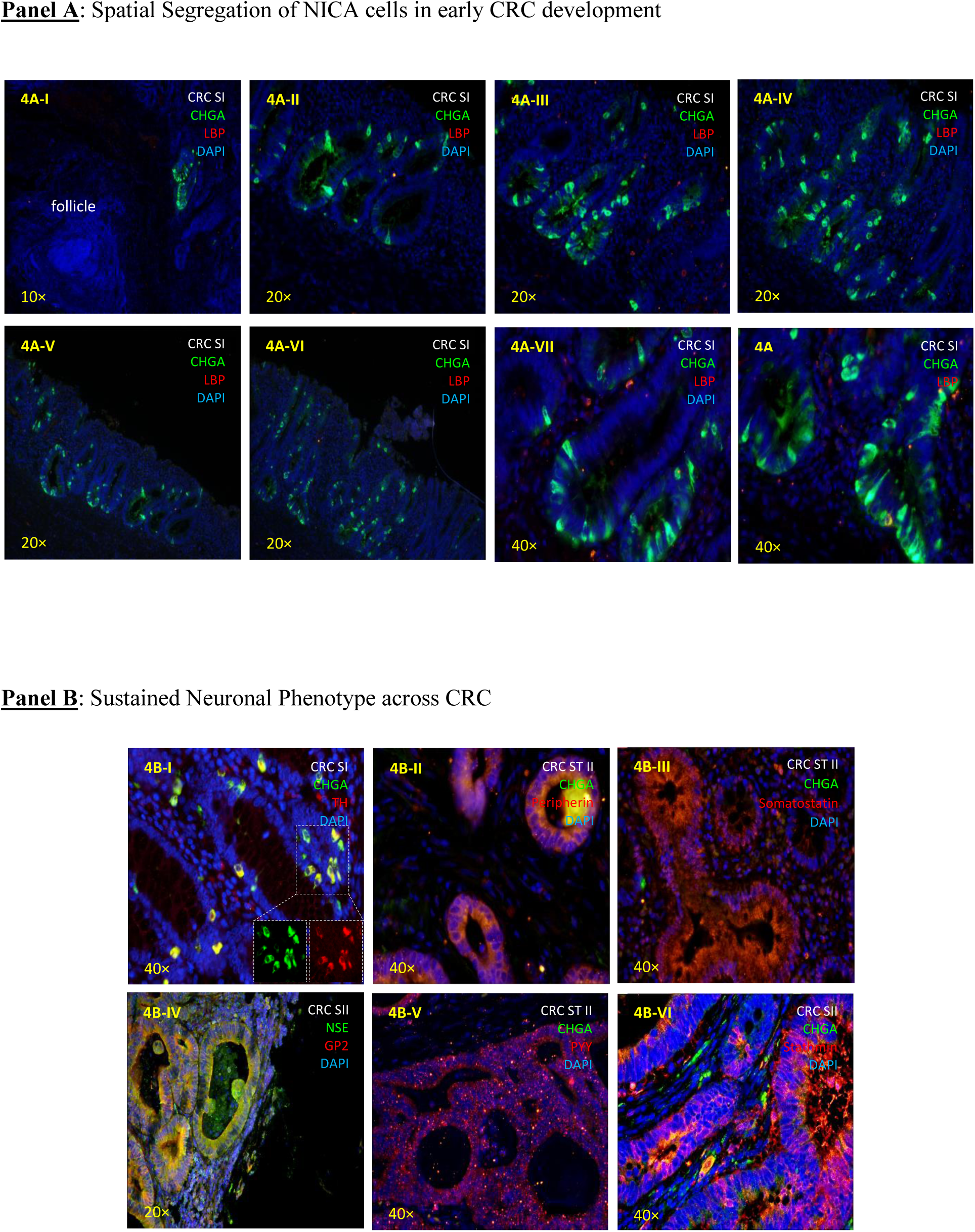
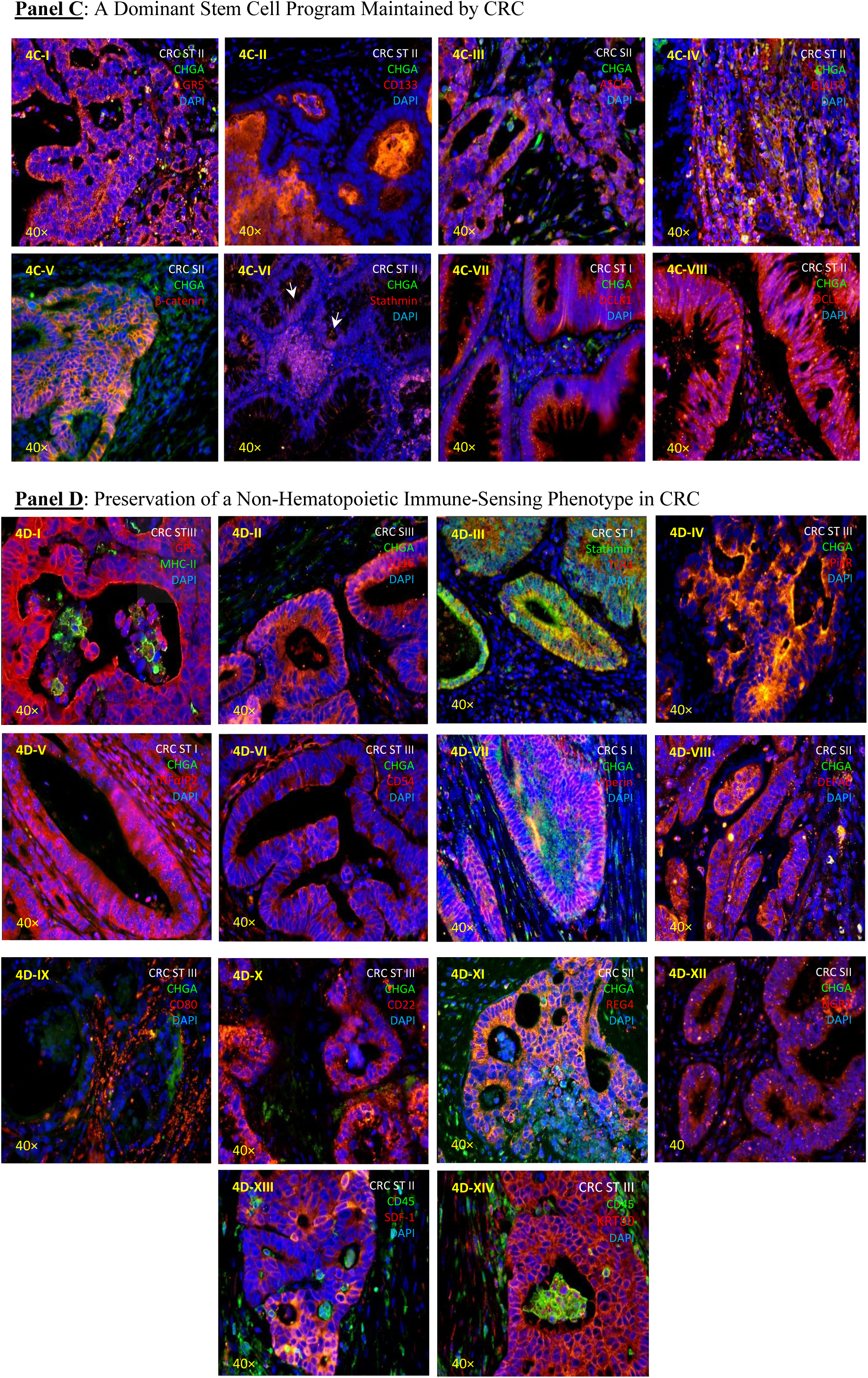
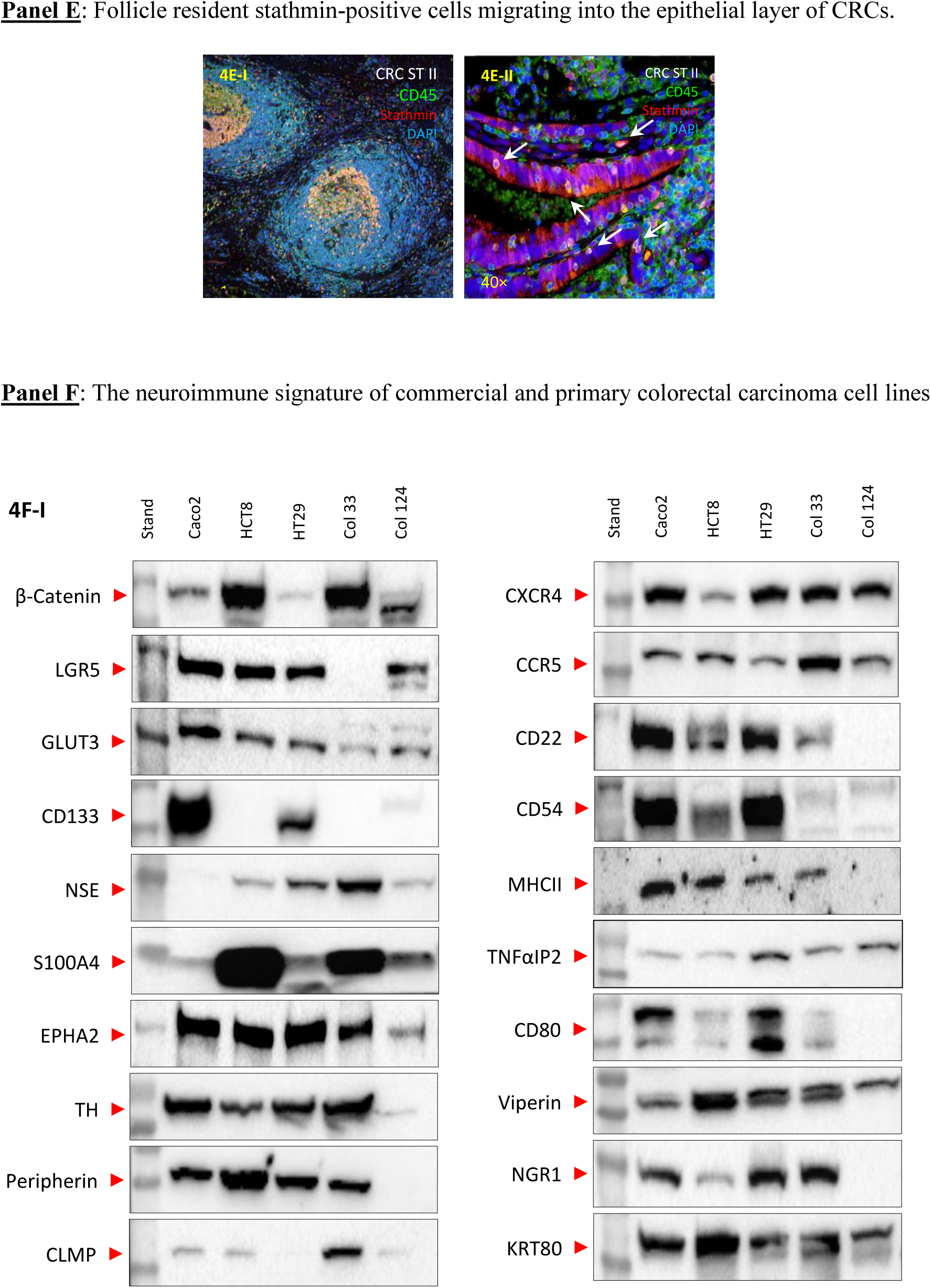
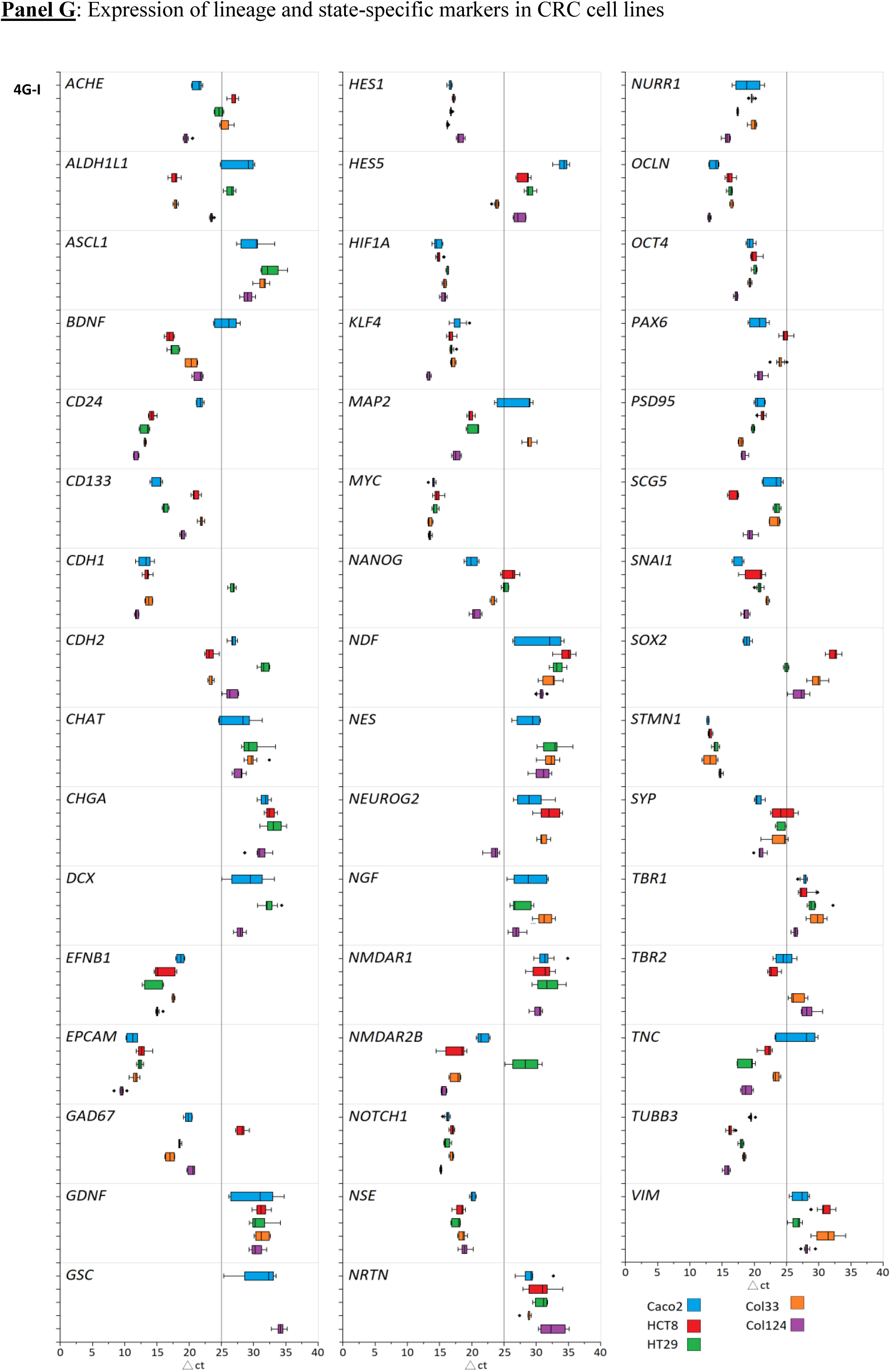
Panel. **A**: IHC. Early CRC shows increased CHGA-positive cells in crypts, especially near lymphatic tissue, with focal or extensive expression from basal portions upward (4A-I–VIII). **Panel B**: IHC. Residual neuronal markers include CHGA, TH, peripherin, somatostatin, NSE, PYY, stathmin, and DCLK1 (4B-I to 4B-VIII). **Panel C**: IHC. CRC retain stemness markers such as LGR5, CD133, ASCL2 (4C-I to 4C-III), GLUT3 (4C-IV) which marks metabolically active progenitor-like cells, β-catenin (4C-V), and stathmin expressed by infiltrating cells from the lamina propria (4C-VI, arrows). **Panel D**: IHC. Immune-related markers include MHCII, Gp2, ZG16, TLR4, PigR, TNFαIP2, CD54, viperin, DEFA6, CD80, CD22, REG4, NGR1, SDF-1 (4D-I to 4D-XIII), as well as the structural protein KRT80 (4D-XIV). **Panel E**: IHC. Stathmin-positive follicular/parafollicular (4E-I) cells migrate into CRC epithelium and fuse with crypt cells (4E-II, arrows). **Panel F**: Western blot. CRC cells show dual neuronal/immune signatures with stemness markers. Caco2, HCT8, and HT29 consistently express LGR5 and GLUT3, (4F-I). Neuronal markers NSE, S100A4, EPHA2, TH, and peripherin are broadly expressed across CRC cells. Immune markers CXCR5, CCR5, TNFαIP2, and viperin are expressed by all CRC cells. CD 22, MHCII, CD80, and NGRI are expressed in the majority of CRC cells, excluding Col 124 primary cells. The structural protein KRT80 is detected across all CRC cells (4F-I). **Panel G**: qPCR across five CRC lines (Caco2, HCT8, HT29, Col33, Col124) shows heterogeneous expression of neuronal (*MAP2*, *NSE*, *SYN*, *CHAT*, *GDNF*), neurodevelopmental (*ASCL1*, *SOX2*, *NES*, *NEUROG2*), stemness (*OCT4*, *NANOG*, *CD133*), epithelial (*EPCAM*, *CDH1*), and mesenchymal (*VIM*, *SNAI1*, *TNC*) markers (4G-I). IHC representative of n=10 patients; WB/qPCR ≥4 replicates.

### Sustained Neuronal Phenotype across CRC

The colorectal tumor architecture is characterized by a robust stemness phenotype defined by the high expression of key progenitor markers. The CRCs retained multiple neuronal features observed in NICA cells, including expression of TH (**Figure 4B**-**I**), Peripherin (**Figure 4B**-**II**), Somatostatin (**Figure 4B**-**III**), NSE (**Figure 4B**-**IV**), and PYY (**Figure 4B**-**V**).

### A Dominant Stem Cell Program Maintained by CRC

The colorectal tumor architecture is characterized by a robust stemness phenotype defined by the high expression of key progenitor markers. CRCs exhibited a strong stemness phenotype, expressing LGR5^72^ (**Figure 4C**-**I**), CD133^73^ (**Figure 4C**-**II**), GLUT3^74,75^ (**Figure 4C**-**III**), ASCL2^76^ (**Figure 4C**-**IV**, β-catenin^77^ (**Figure 4C**-**V**). In contrast to NICA cells, CRCs were highly positive for Stathmin (**Figure 4C**-**VI**), a protein crucial for neural progenitor maintenance^78^ and hematopoietic stem cell function^79^. Additionally, the neuronal and stem cell marker DCLK1 was highly expressed. Its expression increases during progression, shifting from moderate expressed in Stage I to very highly expressed in Stage II. (**Figures 4B**-**VII** &**VIII**).

### Preservation of a Non-Hematopoietic Immune-Sensing Phenotype in CRC

The colorectal tumors maintain a sophisticated “sentinel” profile attributable to their NICA lineage, despite the absence of traditional leukocyte markers. Immunohistochemical analysis confirms that while CRCs are CD45-negative and lack MHC-II expression in patient paraffin sections, they express a broad array of innate immune-sensing and mucosal defense antigens. IHC profiling confirmed a robust innate immune-sensing phenotype through the expression of GP2, ZG16, and TLR4, which mediate bacterial uptake and pattern recognition, alongside PigR, TNFαIP2, and CD54 for immunoglobulin transport and inflammatory signaling. The tumors further produce antimicrobial and regulatory factors, including Viperin, DEFA6, CD80, CD22, REG4, NRG1, and the chemoattractant SDF-1(**Figures 4D**-**I** **to 4D-XIV**). This complex sentinel identity is anchored by the structural protein KRT80, a keratin signature uniquely shared with NICA cells, confirming that these CRC cells function as specialized, non-hematopoietic immune sensors.

Strikingly, for the majority of antigens analyzed, protein detection by IHC was mirrored by transcriptional upregulation of the corresponding genes in freshly isolated tumor crypts, indicating that these markers do not merely represent residual or stabilized proteins but are actively maintained as part of a reinforced transcriptional program. This concordance is particularly evident for innate immune, secretory, stem cell, and metabolic regulators (e.g., *CD54*, *CD80*, *TLR4*, *DEFA6*, *REG4*, *ZG16B*, *LGR5*, *PROM1*, *SLC2A3*, *CTNNB1*), underscoring that the malignant crypt compartment is transcriptionally locked into a niche remodeling state (SI Transcriptome analysis).

### Follicular Stathmin^+^ B-lymphocyte Migration and Fusion with Nascent Tumor Crypts

Immunohistochemical analysis of the subepithelial compartment reveals organized lymphatic follicles characterized by a distinct internal architecture. Within these follicles, Stathmin-positive cells are localized primarily to the central germinal centers. These cells represent a specialized fraction of the broader CD45^+^ leukocyte population (**Figure 4E**-**I**). In terms of distribution and form, the Stathmin-positive cells appear as a dense cluster within the follicle core, contrasting with the more peripheral distribution of other CD45^+^ subsets. Observations indicate an active migration of these Stathmin-expressing cells, which exit the central follicle and move toward the surrounding tissue.

Observations of the tissue architecture indicate that Stathmin^+^ immune cells abandon their central position within the lymphatic follicles and migrate toward the nascent tumors (**Figure 4E**-**II)**. This directed movement results in the infiltration of these follicular-derived immune cells into the surrounding stroma as they move specifically toward the tumoral crypts. Since Stathmin is absent in regular crypt cells but abundant in leukocyte infiltrates, its presence in tumor cells suggests acquisition via cellular fusion. This hypothesis is supported by immunohistochemical evidence showing Stathmin^+^-B-lymphocytes infiltrating colonic crypts and undergoing cell-cell fusion, particularly in early tumor stages (**Figure 4E**-**II**, arrows). Furthermore, transcriptome analysis of the tumor crypts reveals a significant gain of genes specifically related to germinal centers (**Figures B3A**-**XII** and **3A**-**XIII**), reinforcing the link between these migrating follicular cells and the developing tumor identity.

### Stable Co-expression of Neuronal, Stemness, and Immune, Markers across CRC Cell Models

To validate the multi-lineage identity observed in patient tissues, we characterized a panel of commercial CRC cell lines and primary cultures, which confirmed that CRC cells maintain a dual neuronal/immune signature intrinsically linked with a robust stemness program. Analysis of Caco2, HCT8, HT29 and the primary CRC Col 33 and Col124 cells via Western blot demonstrated the consistent expression of the stem cell marker LGR5, the metabolic marker GLUT3, and the core regulators β-catenin and CD133 (**Figure 4F-I**, SI: Western blots & densitometrics). The neuronal component of this signature is broadly represented across all CRC models through the protein expression of NSE, S100A4, EPHA2, TH, and Peripherin. Parallel to these markers, the immune-sensing repertoire is maintained by the universal expression of CXCR4, CCR5, TNFαIP2, and Viperin. Furthermore, a majority of the cell lines -with the exception of Col124 primary cells-exhibit robust expression of CD22, CD80, and NRG1, alongside CD54. With respect to MHCII, although this immune marker is undetectable in CRC paraffin sections and in cultured CRC cells under basal conditions, its expression is robustly induced in vitro following exposure to IFN γ, as confirmed by Western blot analysis (SI: Induction of MHCII molecules in CRCs by IFN γ). Finally, the structural protein KRT80 was detected across all examined CRC cells, reinforcing the stability of this specialized cytoskeleton in vitro.

### Neuronal and Stemness Persistence in CRC In-Vitro Models

The persistence of the neuronal and stem cell programs in CRC was validated at transcriptional level using a panel of commercial cell lines and primary cultures, revealing a remarkably stable co-expression of these markers in vitro. qPCR analysis across Caco2, HCT8, HT29, Col33, and Col124 confirmed a robust neurogenic program characterized by the high-level expression of core neuronal markers such as *CHGA*, *SYP* (Synaptophysin), *ENO2* (NSE), and *DCX* (Doublecortin), alongside specialized structural and axonal proteins including *PRPH* (Peripherin) and *TUBB3* (β3-Tubulin). This neuronal identity is intrinsically linked to a primitive stemness state, as evidenced by the consistent detection of LGR5 and the high-affinity glucose transporter *SLC2A3*, alongside a suite of pluripotency factors including *SOX2*, *NANOG*, *POU5F1* (OCT4), and *KLF4*. The transcriptome further highlights a commitment to a neuronal lineage through the expression of key transcription factor genes such as *ASCL1*, *PAX6*, *NEUROG2*, and *GSC* (Goosecoid), as well as machinery for neurotransmitter signaling including *CHAT*, *GAD67*, and the glutamate receptors *NMDAR1* and *NMDAR2B*. Furthermore, these cells maintain trophic support pathways via *BDNF*, *NGF*, and *GDNF* (**Figure 4G**-**I**).

### Immune Landscape Alterations in CRC

To explore systemic immune alterations in colorectal cancer (CRC) patients, particularly in the context of BLEICS cell involvement, we established a structured, age stratified baseline that enables mechanistic interpretation across the human aging trajectory. We performed comprehensive flow cytometric profiling of peripheral blood leukocytes from 102 CRC patients (mean age 70) and compared them to young healthy individuals (<30 years) and elderly controls (mean age 82). This three tiered design allowed us to disentangle CRC associated immune distortion from the underlying processes of age related immunosenescence, thereby generating a population level immune landscape spanning youth → aging → cancer. All cytometric parameters, reagents, and protocols were rigorously standardized across groups to ensure comparability and to support mechanistic inference.

### Granulocyte to Lymphocyte Ratio as a Sentinel Marker of Systemic Immune Impairment in CRC

The systemic immune profile of CRC patients revealed a profound deviation from normal immune balance, characterized by pronounced myeloid skewing rather than any clinically defined granulocyte pathology. This shift reflects a relative expansion of the myeloid compartment -driven primarily by increased granulocyte abundance -together with a substantial reduction in circulating lymphocytes. Whereas healthy aging produced a moderate rise in the Granulocyte to Lymphocyte Ratio (GLR) from 1.96 to 3.38, CRC patients exhibited a striking escalation to 6.02 (p < 0.001) (**Figure 5A**-**I**). Clinically, such an elevated GLR indicates a tumor-associated systemic inflammatory state marked by emergency myelopoiesis, dominance of granulocytic myeloid populations, and reduced adaptive immune capacity. This myeloid dominant immune architecture is associated with poorer prognosis, diminished therapeutic responsiveness, and increased systemic vulnerability, underscoring the GLR as a clinically meaningful integrative biomarker of CRC related immune dysregulation^80^.

**Figure 5:**
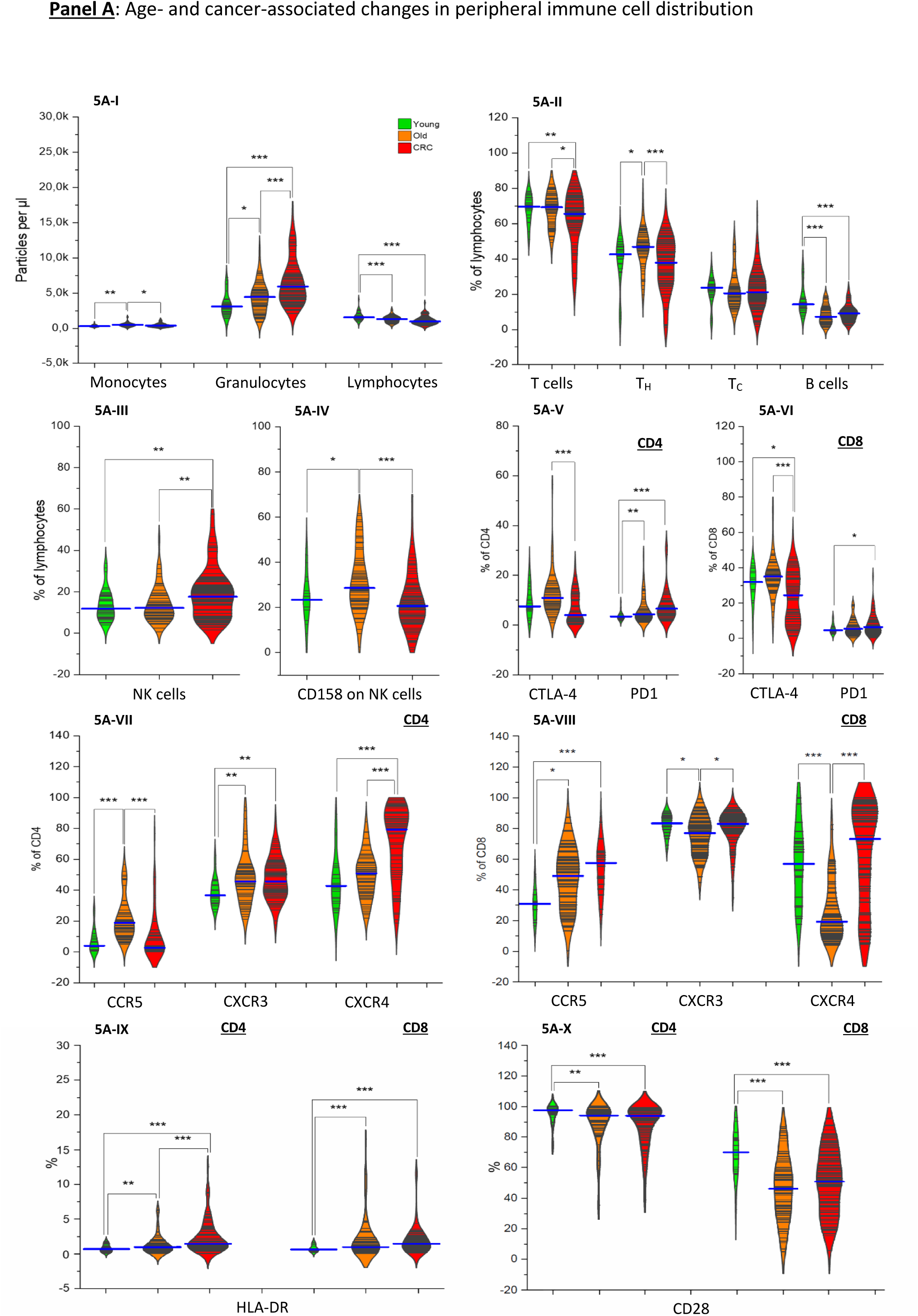
Immune landscape alterations in colorectal cancer patients. FACS analysis of peripheral immune cells across three groups (Young, 25y; Older, 83y; and CRC, 72y) reveals a profound systemic shift characterized by myeloid expansion and adaptive immune dysfunction. The Granulocyte-to-Lymphocyte Ratio (GLR) proved to be the most sensitive marker of this imbalance; while physiological aging increased the ratio from 1.96 to 3.38, the CRC group exhibited a dramatic escalation to 6.02 (***p < 0.001) (5A-I). Lymphocyte subset distribution (5A-II) shows a significant reduction in CD4^+^ frequency in CRC patients compared to both healthy cohorts, while CD8^+^ levels remain stable and B-cells (CD19^+^) show a partial recovery compared to the older cohort. Innate lymphoid alterations (5A-III, IV) include a relative expansion of NK cells (**p < 0.01) paired with a significant loss of CD158 expression, suggesting an immature or anergic NK phenotype. In contrast to classic exhaustion, CRC T-cells exhibit a significant downregulation of the checkpoint CTLA-4 (***p < 0.001) with a significant change in PD-1 levels (5A-V, VI). Finally, analysis of trafficking markers (5A-VII, VIII) reveals a significant upregulation of CCR5 and CXCR4 in CD4^+^ cells and CXCR4 in CD8^+^ cells (***p < 0.001), suggesting a high potential for T-cell sequestration into the tumor microenvironment. Paradoxically, remaining CD4^+^ and CD8^+^ cells show increased HLA-DR expression compared to controls (*p < 0.05), indicating chronic activation (5A-IX), yet they suffer from a deficit in costimulation due to significant CD28 downregulation (*p < 0.05) (5A-X). Flow cytometry was performed on freshly collected EDTA blood. For all median and p values see SI: Immune status median and p values.

### CD4 T-cell Loss and NK Cell Activation Mark Systemic Immune Remodeling in CRC

The most consistent and statistically robust alteration within the lymphoid compartment was a reduction in CD4 T helper cells, which emerged as a defining immunological distinction between CRC patients and healthy controls. This decline reflects a loss of adaptive coordination capacity and contributes to the broader immune impairment observed in CRC. B-cell counts were also reduced in both elderly individuals and CRC patients, while CD8 cytotoxic T cells remained relatively stable across all groups, indicating selective attrition within the helper and humoral arms of adaptive immunity (**Figure 5A**-**II)**. In apparent compensation, natural killer (NK) cells were significantly elevated in CRC patients (**Figure 5A**-**III**), and their reduced expression of the inhibitory receptor CD158 (**Figure 5A**-**IV**) suggests a shift toward a more activated cytotoxic phenotype. However, this innate response appears insufficient to control tumor progression, implying the presence of tumor-intrinsic resistance mechanisms or microenvironmental constraints that suppress NK cell efficacy despite their numerical and phenotypic activation.

### Reduced CTLA-4 and Elevated PD 1 Mark T-cell Exhaustion in CRC

Checkpoint molecule profiling revealed significantly reduced CTLA-4 expression in both CD4 and CD8 T cells from CRC patients (**Figure 5A**-**V** **and 5A**-**VI**), which may limit the effectiveness of CTLA-4 -directed immunotherapies in this population. In contrast, PD-1 expression was markedly elevated, particularly on CD4 cells (**Figure 5A**-**V**), indicating a state of chronic activation or functional exhaustion.

### Age and Tumor Driven Rewiring of T-cell Chemokine Receptors

Chemokine receptor profiling revealed coordinated but lineage-specific remodeling of trafficking potential across CD4 and CD8 T cells. On CD4 T cells, CXCR5 was markedly elevated in elderly individuals compared with the young cohort (p < 0.001), but significantly downregulated in CRC patients relative to age matched controls (p < 0.001), indicating a loss of follicular homing capacity in the tumor context. CXCR3 showed significant differences between young and older individuals (p < 0.01) and between young and CRC patients (p < 0.01), while remaining unchanged between CRC and elderly controls, suggesting age driven rather than tumor specific modulation. CXCR4 expression was strongly overexpressed in CRC patients compared with elderly controls (p < 0.001), consistent with tumor-driven alterations in lymphoid trafficking dynamics (**Figure 5A**-**VII**). On CD8 T cells, CXCR5 was significantly upregulated in both elderly individuals (p < 0.05) and CRC patients (p < 0.001) relative to young controls. CXCR3 was reduced in the elderly group compared with young individuals (p < 0.05), yet CRC patients expressed higher CXCR3 levels than their aged controls (p < 0.05), indicating a partial tumor-associated activation signal. CXCR4 showed a highly significant reduction in elderly individuals compared with the young cohort (p < 0.001), but was strongly overexpressed in CRC patients relative to their aged controls (p < 0.001), suggesting tumor-driven redirection of CD8 T-cell trafficking toward CXCR4 dependent niches (**Figure 5A**-**VIII**).

### T-Cell Activation and Co-Stimulatory Remodeling

Expression profiling of activation and co-stimulatory markers revealed coordinated remodeling of T-cell functionality across aging and colorectal cancer (CRC). HLA-DR expression was significantly elevated on CD4 T cells in elderly individuals compared with the young cohort (p < 0.01), and further increased in CRC patients relative to age-matched controls (p < 0.001), indicating progressive immune activation. On CD8 T cells, HLA-DR was markedly upregulated in both elderly individuals and CRC patients compared with young controls (p < 0.001), reflecting sustained cytotoxic activation in response to age-related and tumor-associated stimuli (**Figure 5A**-**IX**). In contrast, CD28 expression declined with age and disease. CD4 T cells showed reduced CD28 levels in elderly individuals (p < 0.01) and even lower expression in CRC patients (p < 0.001), while CD8 T cells exhibited significant CD28 loss in both elderly and CRC cohorts (p < 0.001) (**Figure 5A**-**X**). Because CD28 expression does not differ significantly between elderly individuals and CRC patients, the observed CD28 loss reflects age associated immunosenescence, not a CRC specific alteration. CRC patients simply retain this senescent, co stimulation deficient phenotype rather than acquiring additional CD28 loss due to the tumor.

### Emergence of a B Cell-Derived, M-Cell-Like Lymphoid Population in CRC Primary Cultures

Establishment of CRC cultures revealed a recurring ≈5 µm lymphocyte-like population in 17% of samples. These cells emerged alongside CAFs and displayed a distinctive squid-like morphology and motility (SI: Videographies 1-8). Approximately half of these populations failed to expand beyond early passages, whereas others persisted and spontaneously immortalized. All source tumors were adenocarcinomas, and no comparable cells arose from adjacent healthy tissue, excluding contamination or primary lymphoma^81^. Light microscopy identified pleomorphic ≈5 µm lymphoid cells with a characteristic polarized, petal-like morphology (**Figure 6A-I**), which was confirmed by scanning electron microscopy (**Figure 6A-II**, white arrows). Transmission electron microscopy revealed electron dense apical lipid droplets suggestive of pathogen trapping structures (**Figure 6A-III**, white arrow), and a schematic representation is provided in **Figure 6A-IV**.

**Figure 6:**
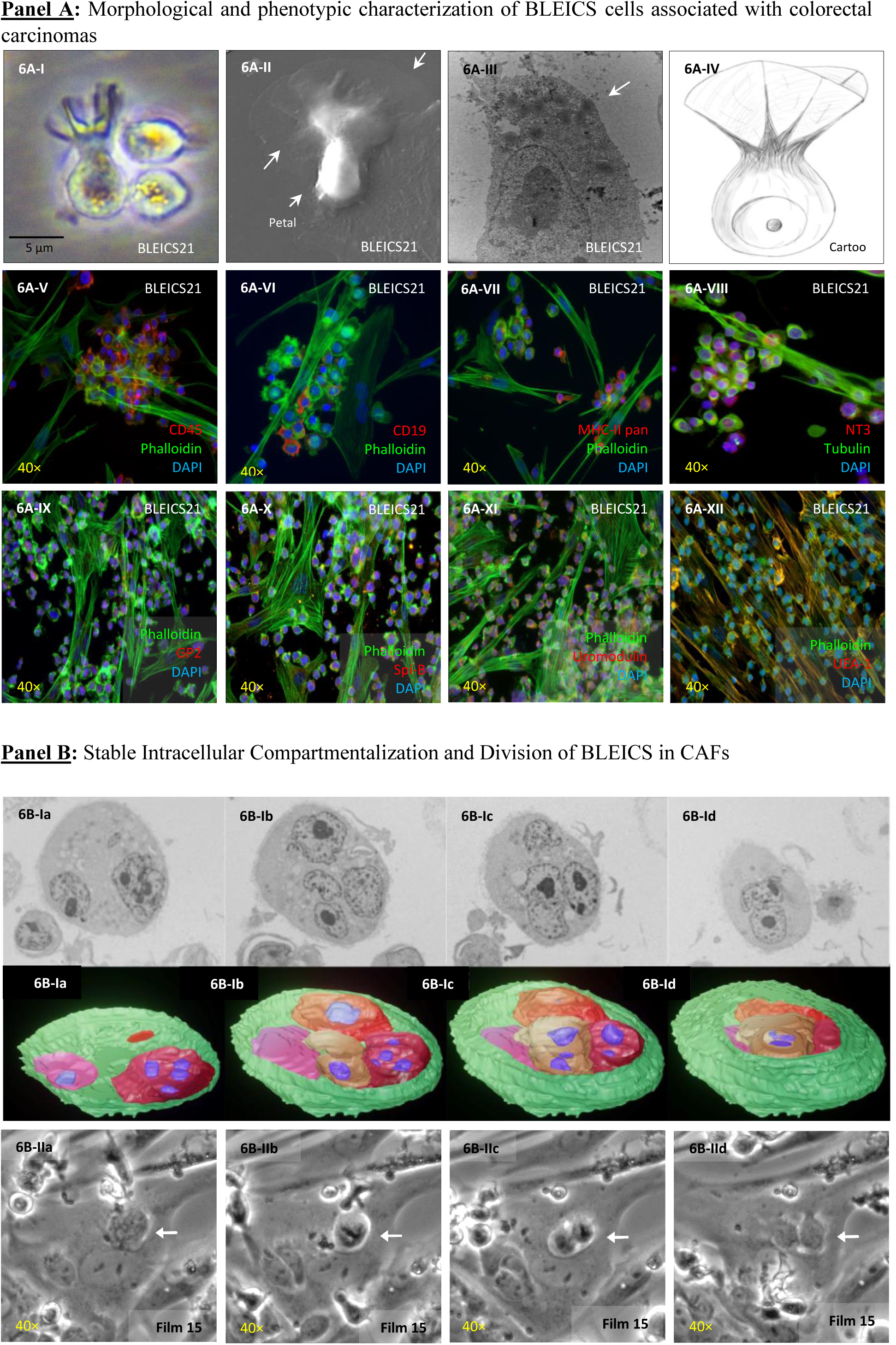
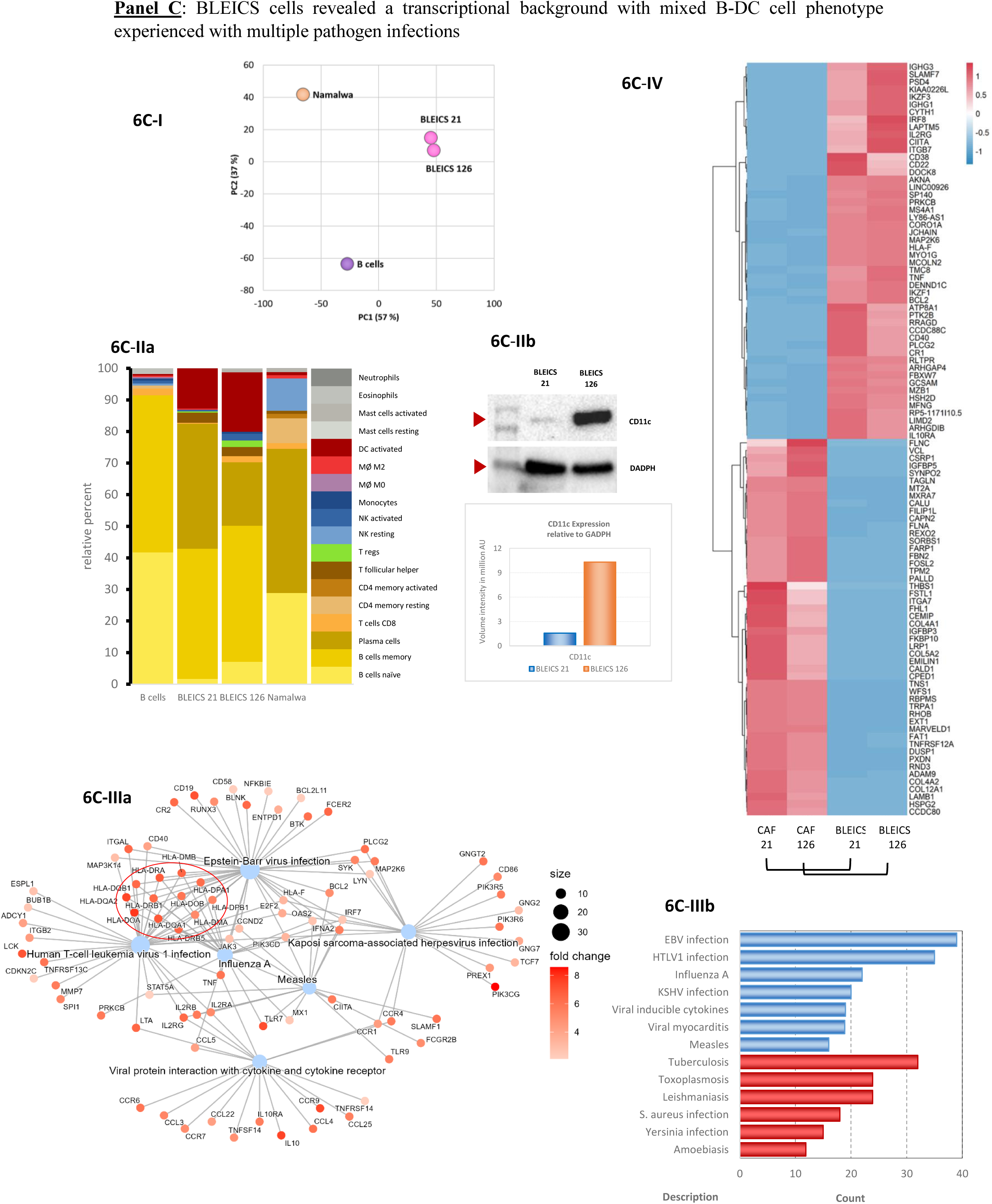
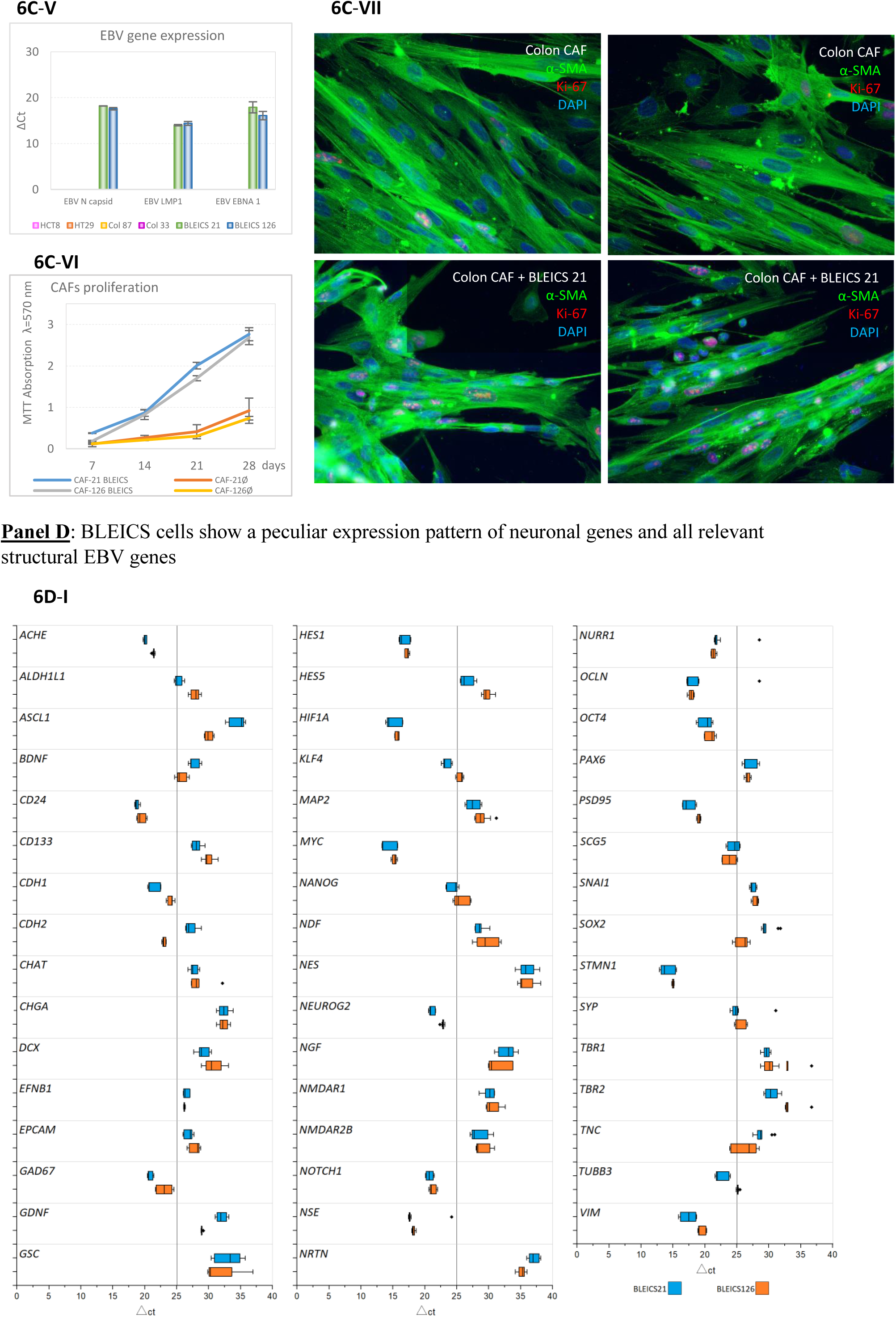
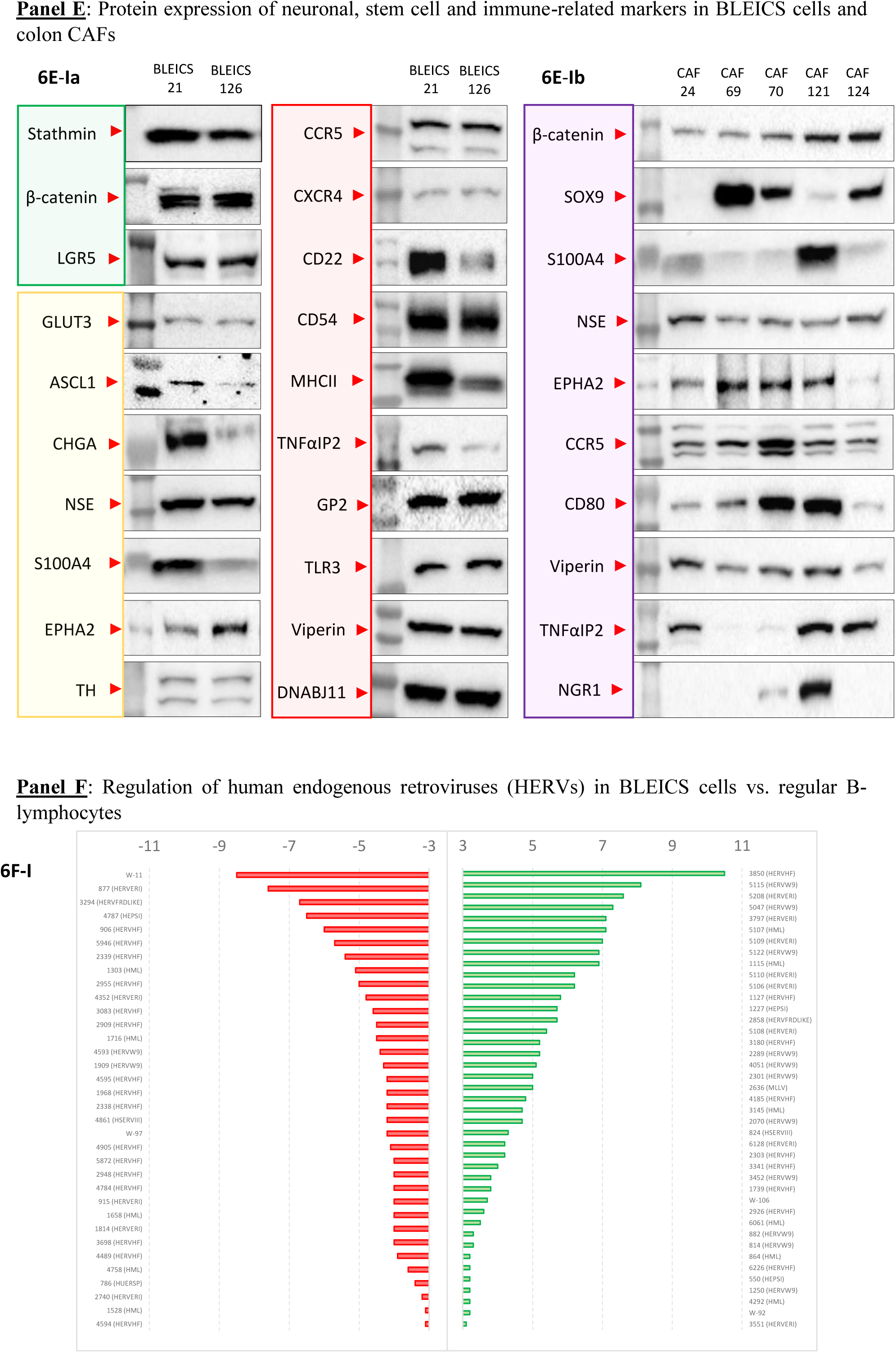
B-lymphocyte, EBV-infected, Calamari-shaped (BLEICS) cells in colorectal carcinomas. **Panel A**: Light microscopy shows pleomorphic lymphoid cells (≈5 µm) with polarized petals (6A-I), confirmed by SEM (6A-II, arrows). TEM reveals electron-dense apical lipid droplets (6A-III, arrow). 6A-IV provides a schematic representation. ICC confirms B-cell phenotype (6A-V to 6AVII: CD45, CD19, and MHCII). BLEICS express NT3 (6A-VIII), M-cell markers (6A-IX to 6A-XI: Gp2, Spi-B, uromodulin) and bind UEA-1 lectin (6A-XII). **Panel B**: Transmission electron microscopy (TEM) reveals BLEIC-CAF fusions forming polykaryocytes, as evidenced by reconstructing sequential TEM sections and generating a 3D model (6B-Ia-d). Intracellular life cycles were observed by time-lapse microscopy (6B-IIa-d; SI Videography 15). **Panel C**: NGS of the top 100 differentially expressed genes distinguishes BLEICs from normal B cells and the Namalwa line (6C-I). CIBERSORTx deconvolution shows memory/plasma cell genotype plus activated DC signature (6C-IIa, SI: CIBERSORTx Results); CD11c validated by WB (6C-IIb). BLEICS carry infection signatures (6C-IIIa & 6C-IIIb: EBV, HTLV-I, influenza A, KSHV, measles virus, M. tuberculosis, S. aureus, T. gondii, Leishmania sp.). CAFs express muscle-related genes (6C-IV: *EMILIN1*, *RHOB*, *FHL1*, *FSTL1*). qPCR confirms EBV transcripts (6C-V: N capsid, LMP1, EBNA1). BLEICS stimulate CAF growth (6C-VI) and increase Ki67 in α-SMA⁺ CAFs under co-culture (6C-VII). **Panel D**: qPCR. BLEICS express neuronal markers (*ACHE*, *BDNF*, *GAD67*, *CDH2* (N-cadherin), *NEUROG2*, *ENO2*, *PSD95*, *SCG5*, SYP (Synaptophysin)) and stemness/EMT markers (*HES1*, *HIF1A*, *KLF4*, *NANOG*, *NOTCH1*, *NURR1*, *VIM)*; *STMN1* is highly expressed (6D-I). **Panel E**: WB shows BLEICS expression of stemness (green), neuronal (yellow), and immune (red) markers (6E-Ia). CAFs from CRC cultures express variable levels of β-catenin, NSE, CCR5, and Viperin (6E-Ib) with variable levels of SOX9, S100A4, among others (6E-Ib, and SI: Western blots & densitometrics). **Panel F**: Differential expression analysis identifies modulated HERV loci in BLEICS cells; bar plots show TPM differences (6F-I, and SI: Table 2 - Differentially expressed HERVs with ISD in BLEICS cells). Results are representative of n=4 experiments. Bars represent ΔTPM values. Statistical significance was defined as adjusted p < 0.05 (Benjamini-Hochberg correction).

Immunocytochemistry and FACS (SI: BLEICS Phenotype by FACS) verified their B-cell identity. We therefore designated these cells BLEICS (B lymphocyte, EBV Infected, Calamari Shaped) to reflect their morphology, viral status, and B-cell phenotype. Phenotypically, all BLEICS were CD45 (**Figure 6A-V**), with a B-cell-like profile detectable in a subset of cells, as indicated by the variable expression of CD19 (**Figure 6A-VI**) and MHC II (**Figure 6A-VII**). BLEICS additionally expressed the neuronal marker NT-3 found in proliferating cells (**Figure 6A-VIII**), displayed M cell-associated markers such as GP2 (**Figure 6A-IX**), Spi-B^82^ (**Figure 6A-X**), Uromodulin^83^ (**Figure 6A-XI**), and bound the M cell-specific lectin UEA-1 (**Figure 6A-XII**), indicating partial functional convergence with mucosal M-cells.

### Stable Intracellular Compartmentalization and Division of BLEICS in CAFs

In vitro, BLEICS showed strong physical association with fibroblasts, dividing on their surfaces or forming dense aggregates reminiscent of hematologic malignancies. TEM analyses revealed distinct nuclear volumes and continuous membrane interfaces between host fibroblasts and the internalized cells, while 3D reconstructions confirmed full spatial encapsulation and a dual-nuclear architecture (**Figure 6B-Ia-d**; SI: 3D reconstruction from TEM). These features support stable intracellular residency of BLEICS-like cells within fibroblast syncytia. Serial section SEM combined with 3D reconstruction further demonstrated their localization within fibroblast polykaryons, reinforcing their presence inside multinucleated stromal compartments.

Time lapse microscopy (**Figure 6B-IIa-d**; Videographies 12-15; SI: Commentaries on Videography) captured the intracellular division of a BLEICS cell within a CAF. White arrows consistently highlighted the BLEICS cell positioned adjacent to the CAF nucleus, while the CAF itself remained morphologically stable throughout the sequence. The BLEICS cell underwent rounding, cytoplasmic reorganization, and segmentation into two compartments, consistent with mitotic division. The absence of membrane rupture or extrusion -corroborated by TEM and 3D reconstructions-confirmed that BLEICS cells remained fully enclosed within the CAF cytoplasm and exhibited dual nuclear volumes. Together, these observations document a previously unreported event: division of a BLEICS cell inside a living CAF. “It is worth noting that long-term cultures of BLEICS experienced CAFs undergo a distinct cellularization process, the nature and drivers of which remain unclear and warrant deeper investigation (SI: Cellularization Processes in BLEICS Experienced CAFs).

### BLEICS Adopt a Unique Transcriptional Program Distinct from B-cell Lineages and Marked by Infection Linked Signatures

Next generation sequencing (NGS) of the top 100 differentially expressed genes revealed a distinct transcriptional profile that clearly separated BLEICS from both normal B cells and the Namalwa B-cell line (**Figure 6C-I**). Principal component analysis (PCA) showed that BLEICS samples clustered tightly together, occupying a transcriptional space far removed from Namalwa and primary B cells, with PC1 and PC2 accounting for 57% and 37% of the variance, respectively. This separation underscores the emergence of a unique gene expression program in BLEICS, distinct from canonical B-cell lineages and transformed cell lines, suggesting lineage divergence or functional reprogramming.

CIBERSORTx deconvolution (**Figure 6C-IIa**) revealed that BLEICS exhibit a mixed transcriptional signature consistent with memory/plasma B cells, alongside an unexpected activated dendritic cell (DC) component. This hybrid identity or transdifferentiation was further supported by CD11c expression, validated at the protein level by Western blot (**Figure 6C-IIb**, red arrowheads) and quantified relative to GAPDH, with BLEICS126 showing markedly higher CD11c levels than BLEICS21. Together, these data suggest that BLEICS adopt a composite immune phenotype, combining B-cell lineage markers with DC-like activation features.

BLEICS exhibited transcriptional signatures linked to diverse pathogens, including EBV, HTLV I, influenza A, KSHV, measles virus, M. tuberculosis, S. aureus, T. gondii, and Leishmania sp. (**Figure 6C-IIIa**/**b**). Network analysis revealed shared high fold change transcripts across multiple viral programs, while infection counts highlighted EBV and HTLV-I as dominant axes of enrichment.

### Stromal Remodeling Driven by BLEICS via Myofibroblast Like CAF Activation

CAFs expressed a distinct set of muscle related genes, including *EMILIN1*, *RHOB*, *FHL1*, and *FSTL1* (**Figure 6C-IV**). Hierarchical clustering of gene expression profiles revealed consistent upregulation of these transcripts in both CAF 21 and CAF 126, suggesting a myofibroblast-like activation state. This stromal signature contrasts sharply with BLEICS profiles and supports a contractile, remodeling competent phenotype in the tumor-associated fibroblast compartment. This profile aligns with the myofibroblast-like activation state that is a hallmark of many CAF populations.

Transcriptome analysis revealed that BLEICS cells harbor both known Epstein-Barr virus subtypes, EBV-1 and EBV-2. qPCR confirmed EBV latency gene expression (*LMP1*, *EBNA1* from EBV-1) in BLEICS cells (**Figure 6C-V**). Cytogenetic analysis showed chromosomal instability with near diploid but aberrant karyotypes in ≈40% of metaphases (SI: Cytogenetic I & II). BLEICS cells carried a pathogenic MUTYH mutation, a lesion strongly associated with colorectal cancer, particularly in the context of MUTYH associated polyposis (MAP), (SI: Genetic Mutations Analysis). Ig gene rearrangement analysis revealed monoclonality in BLEICS21 and biclonality in BLEICS126, with two productive VH rearrangements (IGHV3 20/D6 16/J4; IGHV5 51/D6 19/J5). All lines exhibited somatic hypermutation (3.0% BLEICS21; 17.0% BLEICS105; 5.9-7.8% BLEICS126), consistent with germinal center/memory B-cell origin. Ig production was confirmed (SI: Clonality & Ig Expression), supporting EBV driven malignant transformation. Together, these findings provide orthogonal validation of the germinal center imprint inferred from deconvolution.

BLEICS cells sustain and stimulate CAF growth in vitro, as demonstrated by proliferation kinetics measured via MTT assay over 28 days in BLEICS associated conditions (**Figure 6C-VI**). Immunofluorescence microscopy revealed increased α SMA and Ki-67 staining in CAFs co cultured with BLEICS, indicating activation and proliferative expansion, consistent with the observed growth dynamics (**Figure 6C-VII**). Moreover, transcriptomic analysis revealed that these CAFs express multiple extracellular matrix-remodeling molecules, including MMP3, MMP14, TIMP1, ADAM9, and TIMP1 4, consistent with a myofibroblast like, matrix reorganizing phenotype (SI: Metalloproteinases).

### BLEICs Adopt a Multilineage Hybrid Identity that Integrates Neuronal, Stem Like, Epithelial, and Immune Programs to Shape Stromal Behavior

BLEICS cells displayed a pronounced neuronal-like transcriptional background, evidenced by the gene expression of *ACHE*, *NEUROG2*, *GAD67*, *ENO2*, *NR4A2* (Nurr1), *PSD95*, and *STMN1* (Stathmin). In parallel, they expressed high levels of stemness/progenitor associated genes, including *MYC*, *POU5F1*, *HES1*, and *NOTCH1*, together with EMT related markers such as Vimentin. Remarkably, despite their hematopoietic origin, BLEICS cells also expressed genes typically restricted to epithelial tissues, including *CDH1* (E cadherin), *OCLN* (Occludin), and *CD24*, indicating ectopic activation of epithelial adhesion programs. The presence of HIF1α further revealed that these cells are hypoxia adapted, consistent with survival in tumor-associated niches characterized by low oxygen availability (**Figure** **6D**-**I**). Collectively, these features highlight profound lineage infidelity, mixed transcriptional programs, and functional heterogeneity among EBV infected B-cell derivatives, providing a molecular basis for their distinct stromal interactions.

To validate the neuro-stem-immune hybrid phenotype of BLEICS cells, we performed immunoblotting across selected lineage and signaling markers (**Figure 6E**-**Ia**, SI: Western blots & densitometrics). Both BLEICS lines exhibited consistently elevated levels of Stathmin, β-catenin, and LGR5 -markers commonly associated with colorectal cancer (CRC) biology -indicating partial adoption of CRC-like transcriptional programs. Protein analysis further confirmed the expression of several neuronal associated markers previously elevated at the transcriptional level, including GLUT3, ASCL1, CHGA (reduced in BLEICS126), NSE, and TH, supporting the neuronal-like identity of BLEICS cells. In addition, proteins such as S100A4 and EPHA2, which are linked to EMT-like plasticity and epithelial/CSC associated signaling, were also detected, consistent with the hybrid and highly plastic phenotype of these EBV derived B-cell derivatives.

Immune related markers were uniformly expressed across both BLEICS populations, indicating a shared immune interactive baseline despite their divergent differentiation states. Both BLEICS21 and BLEICS126 expressed moderate to high levels of CCR5 and CXCR4, suggesting chemokine responsive migratory potential. Expression of CD22 and CD54 supports retention of B-cell lineage features and adhesion competence, while robust MHCII expression indicates preserved antigen presentation capacity. Elevated TNFαIP2 and GP2 point to inflammatory and mucosal associated signaling programs, respectively. Innate immune sensors such as TLR3 and the antiviral effector Viperin were also consistently expressed, together with the stress responsive chaperone DNABJ11. Collectively, this uniform immune marker profile highlights a conserved immune reactive and antiviral signature across BLEICS lines, independent of their transcriptional divergence (**Figure 6E-Ia**). However, CAFs associated with different BLEICS populations showed markedly less stable expression patterns, with variable levels of β-catenin, NSE, CCR5, and Viperin. This heterogeneity suggests that CAFs dynamically adjust their signaling and stress response programs depending on the specific BLEICS line they interact with, reflecting context dependent stromal plasticity rather than a fixed activation state (**Figure 6E-Ib**).

### Differentially Expressed HERVs with Intact ISDs in BLEICS cells

ERVmap analysis identified 164 DE HERVs between BLEICS and normal B cells (**Figure 6F-I**). Upregulated elements included 74 Class I, 20 Class II, and 4 Class III ENV proteins; 24 retained SU domains, 56 TM domains, and 21 encoded both. Downregulated elements comprised 49 Class I, 14 Class II, and 3 Class III, with 18 possessing both SU and TM domains.

Focus on ISD containing elements (per Vargiu et al)^71^ revealed 36 DE HERVs with intact ISDs - 17 upregulated, 19 downregulated. These domains were highly conserved, with only minor substitutions (SI: Table 2: Differentially expressed HERVs with ISD in BLEICS cells). The preservation of fusogenic and immunosuppressive potential underscores functional relevance to BLEICS biology, suggesting ISDs may actively modulate host immunity.

Altogether, transcriptomic, deconvolution, and mechanistic analyses support BLEICS as a pathologically reprogrammed B-cell population with myeloid-like functions. CIBERSORTx revealed a stable B myeloid hybrid defined by co-expression of plasma/memory B-cell markers with APC signatures (activated DCs, M1 macrophages). This convergence suggests a B-DC or B-macrophage fusion event, notably supported by high expression of fusogenic HERV envelope proteins.

### EBV-Positive B-Cell Systems Exhibit Distinct Levels of Antiviral Effector Activation

Transcriptomic profiling revealed that EBV-positive B-cell populations differ markedly in the magnitude of their antiviral effector programs. Primary EBV-positive B lymphocytes isolated from healthy individuals displayed a modest antiviral signature, with *OAS1* (52-88 TPM), *OAS2* (74-131 TPM), *MX1* (41-96 TPM), *RSAD2* (3.1-7.4 TPM), and *BST2* (112-284 TPM). In contrast, the constantly replicating EBV-positive B-cell populations BLEICS21 and BLEICS126, which express a broad repertoire of EBV genes, showed substantially higher expression of all major antiviral effectors. BLEICS21 exhibited *OAS1* at 214 TPM, *OAS2* at 1,842 *TPM*, *MX1* at 612 TPM, *RSAD2* at 248 TPM, and *BST2* at 3.126 TPM, while BLEICS126 showed similarly elevated levels (*OAS1* 198 TPM, *OAS2* 1.674 TPM, *MX1* 544 TPM, *RSAD2* 221 TPM, *and BST2* 2.987 TPM). Namalwa cells, representing a fully transformed EBV-positive B-cell line, displayed the strongest antiviral effector activation, with OAS1 at 1.124 TPM, *OAS2* at 3.982 TPM, *MX1* at 1.876 TPM, *RSAD2* at 1.142 TPM, and *BST2* at 4,218 TPM. Together, these data demonstrate that antiviral effector expression varies substantially across EBV-positive B-cell systems, with primary B cells showing a modest baseline, BLEICS21/126 exhibiting a strongly amplified antiviral state, and Namalwa defining the upper limit of EBV-associated interferon-stimulated gene activation.

### BLEICS-Mediated Transactivation of the HERV-H/F Superfamily

Transcriptomic profiling identifies HSERVIII (2692) as the primary molecular marker of the tumor microenvironment, representing the most significantly upregulated retroviral element in tumor crypts with a ΔTPM of approximately 7 (**Figure 3C**, **III**). This locus maintains 70-80% SU homology with HERV-Fc1, driving a high local density of retroviral envelope proteins within the crypt landscape.

Experimental validation in HCT8 and primary Col33 cell lines confirms that 72-hour exposure to EBV-harboring BLEICS triggers a focused, high-amplitude transactivation of the HERV-H/F superfamily (**Figure 7-AI**-**II**). This interaction specifically induces a massive spike in HERV-Fc1 env and HERV-H env transcripts, suggesting that EBV-positive proteins act as potent transactivators of these shared LTR motifs. While both models demonstrate robust and specific induction, HCT8 exhibits a higher and more sustained pool of HERV RNA -peaking at nearly 8-fold for HERV-Fc1 env-whereas Col33 displays a similar but more transient response.

**Figure 7:**
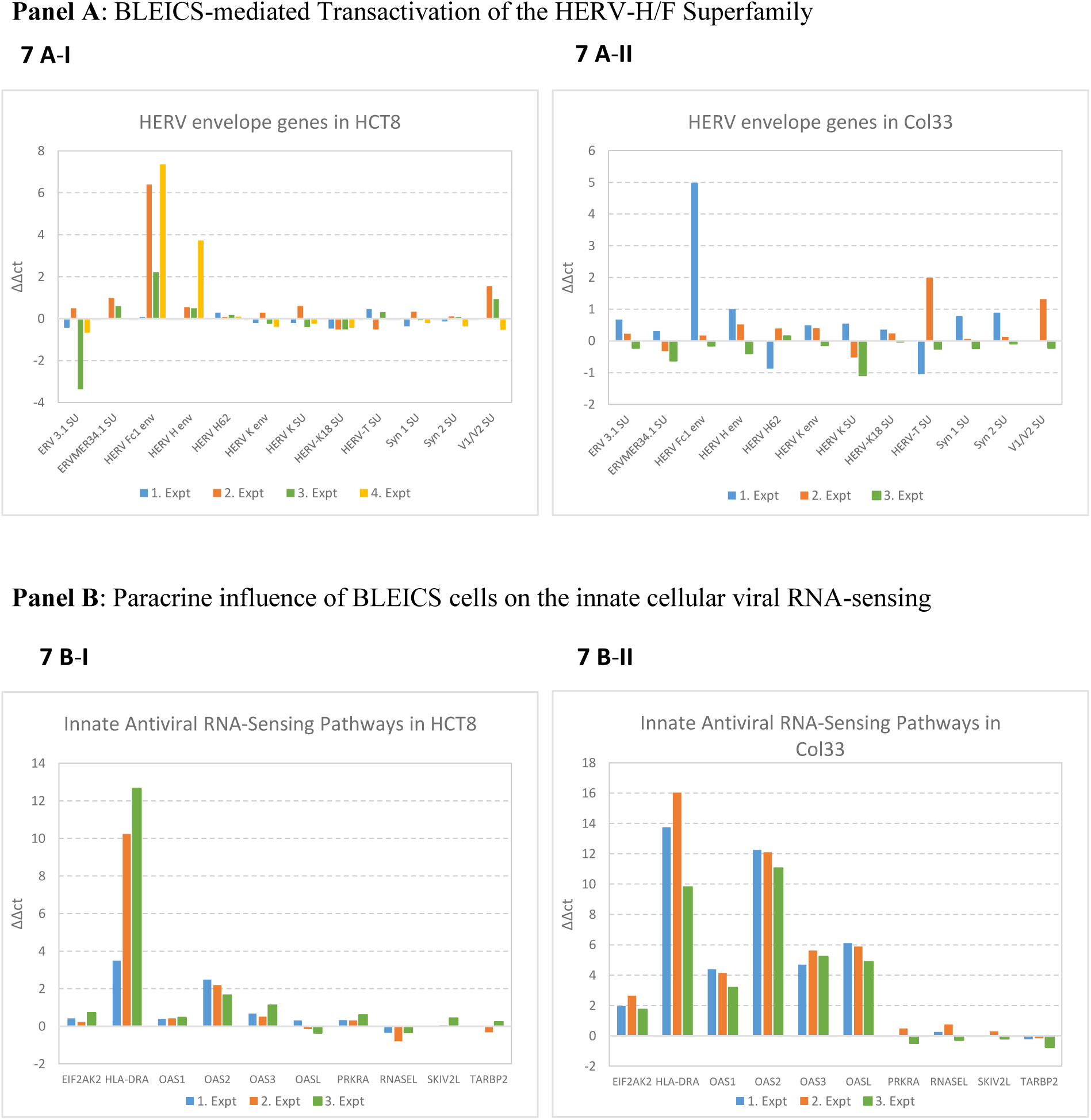
<u>Panel A</u>: BLEICS-mediated transactivation of the HERV-H/F superfamily. Relative expression of HERV elements in HCT8 (left) and primary Col33 (right) cells after 72 h exposure to EBV-harboring BLEICS (four independent replicates). BLEICS interaction induces a focused, high-amplitude activation of the HERV-H/F superfamily, with strong upregulation of HERV-Fc1 env and HERV-H env. HCT8 shows the highest and most sustained induction (approaching ∼8-fold for HERV-Fc1 env), whereas Col33 displays a similar but more transient response. This selective transactivation defines the elevated retroviral transcript pool characteristic of the tumor crypt. <u>Panel B</u>: Paracrine activation of innate viral RNA-sensing pathways. Bar charts show relative expression of ten antiviral genes across three independent experiments following interaction with EBV-harboring BLEICS. Both cell lines upregulate HLA-DRA, but Col33 exhibits a markedly stronger and more consistent antiviral signature, particularly across the OAS family (OAS1, OAS2, OAS3, OASL). In contrast, RNASEL and TARBP2 remain near baseline or slightly reduced in both models, indicating that BLEICS exposure preferentially activates the OAS/RNase L axis rather than broadly inducing all antiviral effectors.

It is important to note that our qPCR analysis targeted only a limited panel of HERV loci, specifically SU encoding regions. Therefore, the full retroviral landscape induced by BLEICS pressure may extend beyond the HERV H/F elements quantified here. A broader transcriptomic survey of HERV families would likely reveal additional loci responsive to immune neighborhood conditioning and could further refine the retroelement signature characteristic of the tumor crypt niche

Collectively, these data indicate that the BLEICS-associated viral stimulus facilitates a localized retroviral transactivation that defines the transcriptomic signature of the tumor crypt.

### Selective Downstream Restricted Antiviral Effector Activation in Tumor Crypts and Its Progressive Loss in CRC *In Vitro* Models

Transcriptomic profiling shows that tumor crypts (TC) mount a selective, downstream restricted antiviral effector program, rather than a broad interferon-like or inflammasome driven response. Within the OAS–RNase L and PKR effector axis, the most significantly upregulated genes are OAS1 (log FC ≈ +1.66, p = 0.0003), OAS2 (log FC ≈ +1.14, p = 0.015), and EIF2AK2/PKR (log FC ≈ +0.52, p = 0.0001). Notably, RNASEL remains downregulated in tumor crypts (SI Transcriptome Analysis), indicating that effector activation is incomplete. In contrast, no upstream viral sensors (RIG I/DDX58, IFI16, ZBP1), no inflammasome components (AIM2, NLRP3, IL1B, IL18), and no interferon ligands (IFNA, IFNB, IFNL) reach statistical significance. Several canonical sensors, including IFIH1/MDA5 and ISG15, are unregulated or entirely absent. This transcriptional pattern rules out acute viral infection, active nucleic acid sensing, or inflammasome activation as drivers of the observed differences and instead reflects a ligand independent antiviral imprint.

To determine whether this downstream restricted program is preserved outside the in vivo niche, we examined the same antiviral effectors in a primary CRC culture (Col124) and in commercial CRC cell lines. Col124 retains a substantial portion of the antiviral signature, with high expression of OAS1 (116-TPM), OAS2 (143-TPM), MX1 (482-TPM), and BST2 (732-TPM). In contrast, commercial CRC lines show a marked collapse of this program: OAS1 drops to 20–42 TPM, OAS2 to 0.11– 2.31 TPM, MX1 to 2.62–4.88 TPM (with partial retention in HCT8), and BST2 to 7.76–43.6 TPM. These data establish a clear quantitative gradient-tumor crypts > Col124 > CRC cell lines-indicating that the antiviral imprint is progressively lost as epithelial cells are removed from their native microenvironment.

Together, these findings demonstrate that the antiviral signature in tumor crypts represents a downstream, ligand independent interferon imprint -a “viral scar”-that is strongest in vivo, partially retained in primary CRC cultures, and largely absent in long-term cell lines. The selective activation of antiviral effectors only in models with preserved sensing competence supports a model in which endogenous retroelement derived dsRNA contributes to viral mimicry, but maintenance of the antiviral imprint requires intact interferon responsive capacity rather than HERV abundance alone.

### The Pseudo-Immune State and the Execution Gap

Co culture of CRC cells with BLEICS reveals a bifurcated antiviral architecture in which HCT8 cells adopt a pseudo immune state, whereas BLEICS retain full antiviral competence. In BLEICS, the transcriptome is dominated by strong induction of OASL (+3.54) and broad activation of the OAS cluster -OAS1 (+1.19), OAS2 (+0.95), OAS3 (+1.55)-together with increased expression of the downstream effector RNASEL (+1.13). This represents a complete and functional antiviral program, consistent with the immune origin of these cells. In contrast, HCT8 cells exhibit a deliberate uncoupling of this pathway. Although they sense the immune neighborhood through selective OAS2 induction (+2.48) and paracrine driven HLA DRA upregulation, they fail to bridge the Execution Gap, as RNASEL remains suppressed (-0.34). This asymmetric pattern closely mirrors the tumor microenvironment (**Figure 7B**, **I**-**II**), where detection signals persist but execution is blocked. Col33 primary cells provide an intermediate phenotype. Like HCT8, they upregulate HLA DRA in response to BLEICS, but unlike HCT8 they mount a stronger and more coherent antiviral signature, with robust induction across OAS1, OAS2, OAS3, and OASL. Despite this enhanced sensing capacity, RNASEL and TARBP2 remain near baseline in both models, indicating that BLEICS exposure preferentially activates the OAS-dominated sensing arm without engaging the full OAS/RNase L effector cascade. This positions Col33 as a partially competent antiviral state, retaining more of the native epithelial sensing machinery than HCT8 but still exhibiting an execution deficit. Importantly, HLA-DRA induction is functionally uncoupled from HERV RNA levels. While HERV transcripts likely provide ligand input for intracellular sensing via OAS2, the upregulation of MHC-II molecules is best explained by paracrine cues from EBV harboring BLEICS, rather than by classical IFNγ signaling. This interpretation is supported by the absence of OAS1 activation, which argues against a canonical interferon-γ response.

### Tumoral crypt expansion is driven by CAF-BLEICS interactions and shaped by early NICA cell accumulation

Aberrant crypt architecture is a common histological feature in colorectal cancer (CRC). Upon enzymatic dissociation of fresh tissues, tumor crypts were found to be up to three times larger than normal crypts (**Figure 8A**-**I** & **II**). This enlargement was strongly associated with stromal proliferation, particularly the expansion of CAFs. As demonstrated in **Figure 3D**-**II**, this fibroblast overgrowth is driven by direct interaction with EBV-infected BLEICS cells, likely mediated by stathmin induction^84^. These interactions contribute to progressive crypt deformation and expansion (**Figure 8A**-**III**).

**Figure 8:**
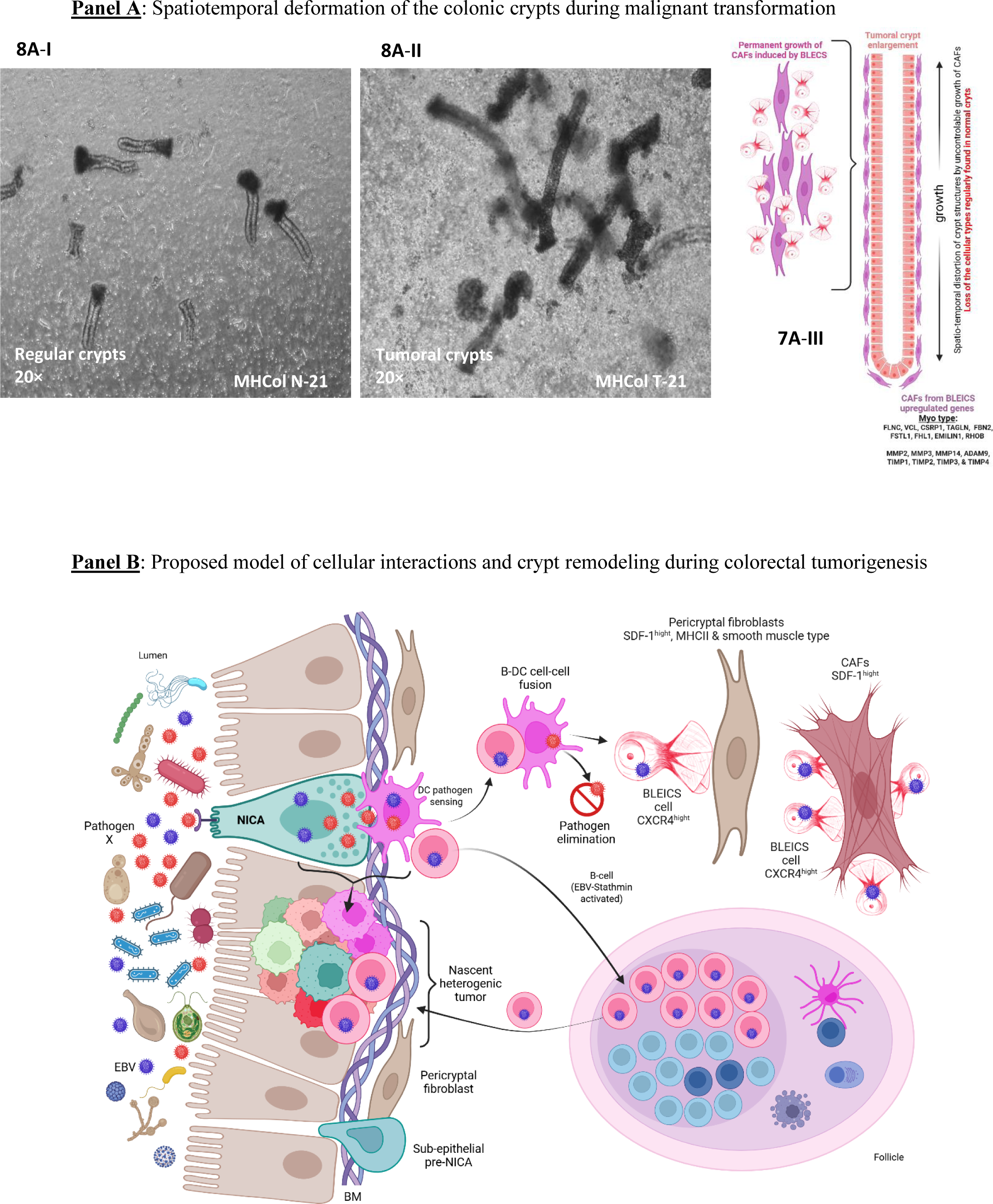
Main events during malignant transformation in CRC. **Panel A**: Comparison of regular vs. tumor crypts shows enlargement up to threefold (7A-I,II). BLEICS cells sustain CAF growth (3D-III); removal slows proliferation and induces senescence. CAFs express smooth-muscle genes, consistent with myofibroblast/pericryptal fibroblast identity, and overexpress MMP3, MMP14, TIMP1 (SI: Metalloproteinases). BLEICS-driven CAF expansion explains stromal remodeling and crypt deformation (7A-III). **Panel B**: Schematic of carcinogenesis pathway. NICA cells may arise from subepithelial compartments (pre-NICA), integrate into crypts, and act-like M cells via pathogen-recognition molecules. EBV recruits dendritic cells, which activate B-lymphocytes; EBV-infected B cells express stathmin and, through DC interaction, form BLEICS (CXCR4^high^). Fibroblast-secreted SDF-1 drives BLEICS-CAF engagement, promoting CAF differentiation and crypt elongation. BLEICS infiltrate tumor tissues. Interaction and potential fusion between NICA cells and BLEICS (B-DC hybrids) may trigger the genetic instability and lineage-blending that characterizes the nascent tumor crypt. Tumor cells display NICA markers and MHC-II (likely DC-derived), but lack immunoglobulin expression. Table 3: Summary of molecular markers and viral receptors. NICA cells (Balalaika-shaped argentaffin) show neuronal, stemness, and immune traits, lectin binding (PNA, UEA-1), and viral receptors (EPHA2, CAR, GP2, CD80, PigR, TLR4, CXCR4). BLEICS cells (Calamari-shaped, EBV-infected B-lymphocytes) combine B-cell/immune identity, neuronal/neuroendocrine programs, stemness/EMT plasticity, and viral/fusogenic activity (EBV transcripts, HERV envelope genes). Together, these features highlight the NICA and BLEICS cells as key oncogenic intermediate enabling immune mimicry, viral susceptibility, and stromal remodeling. Cartoons generated with BioRender.com.

**Table 3:** Summary of molecular markers and viral entry receptors in NICA and BLEICS cells.

| Category | NICA | BLEICS |
| --- | --- | --- |
| Neuronal /<br>Neuroendocrine | CHGA , TH, Peripherin,<br>Somatostatin, NSE, PYY,<br>Stathmin, EPHA2 | CHGA, TH, NSE, Peripherin, Stathmin<sup>high</sup>,<br>Synaptophysin, PSD95, Neurog2, GAD67,<br>BDNF, NT3, MAP2, Doublecortin, Nestin,<br>CHAT, Ephrin B1/EPHA2 |
| Stemness /<br>Regenerative | LGR5, CD133,<br>GLUT3/SLC2A3, ASCL2,<br>β-catenin, DCLK1, KRT80 | SOX2, Nanog, Oct4, Klf4, c-Myc, ASCL1,<br>HES1, HES5, Nurr1, β-catenin, S100A4,<br>Vimentin, N-cadherin, Snail, Notch1,<br>ALDH1L1, SCG5 |
| Innate Immune | Lectins: PNA, UEA-1; GP2,<br>ZG16, TLR4, REG4, PigR,<br>TNFαIP2, CD54, Viperin,<br>DEFA6, SDF-1 | GP2, REG4, PigR, TNFαIP2, CD54,<br>Viperin, TLR3, SDF-1, NgR1 |
| Adaptive Immune | CD22, CD80, MHC-II (low),<br>CD45 absent | CD45<sup>+</sup>, CD19<sup>+</sup>, MHC-II (HLA-DR/DP/DQ),<br>CD22, CD80 |
| Viral Entry<br>Receptors | EPHA2 (EBV), GP2<br>(Rhinovirus/M-cell), CD80<br>(EBV), PigR, TLR4, CAR<br>(HAdVs ) CXCR4 | CXCR4<sup>high</sup> (HIV, EBV), CCR5 (HIV),<br>EPHA2 (EBV), NgR1 (EBV), Gp2, CD22<br>(EBV),<br>CD80 (EBV), MHC-II (EBV),<br>Uromodulin/UEA-1 binding |
| Viral / Fusogenic<br>Features | Susceptible to EBV and<br>Adenoviral infections;<br>pathogen gateway | EBV transcripts (N capsid, LMP1, EBNA1);<br>HERV envelope genes with ISD; pathogen<br>signatures (HTLV-1, Influenza A, KSHV,<br>Measles, Mycobacterium, S. aureus,<br>Toxoplasma, Leishmania) |
| Functional<br>Interactions | Pathogen sensing, immune<br>mimicry, crypt integration | Fusion with CAFs (CXCR4-SDF-1 axis),<br>stromal remodeling, polykaryocyte<br>formation, CAF proliferation, crypt<br>deformation |

### Proposed Model for BLEICS Cell Mediated Colorectal Carcinoma Pathogenesis: An Evolutionary Hybrid Lineage as an Active Actor in Colonic Carcinogenesis

We propose a multi-stage model where Colorectal Carcinoma (CRC) pathogenesis is initiated by the formation of BLEICS cells through a unique heterotypic fusion event driven by chronic viral infection in the tumor-initiating niche (**Figure 8B**).

1. NICA cells reside at crypt bases adjacent to isolated lymphoid follicles and integrate neuronal, neuroendocrine, stemness, and innate immune programs, positioning them as pathogen sensing epithelial sentinels.
2. Chronic microbial and viral exposure recruits CXCR4 B cells via NICA derived SDF 1 and establishes a follicle conditioned microenvironment in which EBV infection drives the emergence of BLEICS intermediates, a multilineage B cell-derived population with activation of fusogenic HERV envelope loci.
3. BLEICS fuse with fibroblasts and epithelial/NICA cells, generating hybrid stromal and epithelial lineages that expand cancer associated fibroblasts, deform crypt architecture, and create a stromal rich, tumor permissive niche.
4. Within this remodeled environment, WNT driven REST upregulation in NICA/BLEICS derived epithelial hybrids selectively extinguishes the deep neuroendocrine transcriptional module (CHGA, *PHOX2A*/*B*, *NEUROD1*, *PAX4*) while preserving a broader neuronal/neuroepithelial scaffold and stemness determinants.
5. This REST mediated pruning converts a neuroimmune sentinel lineage into a proliferative neuro immune epithelial cancer cell, explaining the coexistence of neuronal background markers with loss of classical neuroendocrine identity in tumor crypts.
6. REST inhibition genetically or pharmacologically reactivates the suppressed neuroendocrine module, induces neuroendocrine miRNAs, and collapses malignant growth, identifying REST as a central lineage gatekeeper linking pathogen exposure, fusion biology, stromal remodeling, and epithelial transformation in colorectal cancer.

#### Concluding Synthesis

##### CRC as an Evolutionary Hybrid Lineage

Colorectal cancer arises from a pathogen-conditioned epithelial niche in which Neuro Immune Crypt Associated (NICA) cells -neuronal, neuroendocrine, stem-like sentinels positioned at crypt bases near lymphoid follicles-sense microbial and viral cues through GP2, TLR4, REG4, lectins, and viral receptors, and recruit CXCR4 B cells via SDF 1. Epstein-Barr virus infects these recruited B cells, generating BLEICS intermediates that combine B-cell identity, APC-like activation, neuronal/neuroendocrine and stemness traits, epithelial adhesion, and strong antiviral signaling while activating fusogenic HERV envelope loci. BLEICS fuse with fibroblasts to expand CAFs and deform crypt architecture, and fuse with NICA or epithelial cells to generate hybrid epithelial lineages that retain neuronal background, stemness, immune sensing capacity, and viral/HERV linked plasticity. Within this remodeled microenvironment, WNT driven REST upregulation selectively extinguishes the deep neuroendocrine transcriptional module (*CHGA*, *PHOX2A*/*B*, *NEUROD1*, *PAX4*) while preserving the broader neuronal scaffold and stemness programs, converting NICA/BLEICS derived hybrids into proliferative neuro immune epithelial tumor cells. Conversely, genetic or pharmacologic REST inhibition reactivates the suppressed neuroendocrine module, induces neuroendocrine miRNAs, and collapses malignant growth, identifying REST as the central lineage gatekeeper linking pathogen exposure, fusion biology, stromal remodeling, and epithelial transformation in colorectal cancer.

## Discussion

For decades, pathologists have noted a shift in colon tumors from neuroendocrine toward epithelial phenotypes^85^. Gene expression studies across multiple histologies reveal a persistent neuronal background, suggesting that many cancers may arise from cells with neuroepithelial features^67,86^. Moreover, tumors also display immune related traits, historically attributed to fusion events between effector immune cells and emerging tumor tissues^87,88^. In CRC, the highest incidence -from the left colic flexure to the distal rectum^23,89^-aligns with regional gradients of endocrine cells and isolated lymphoid follicles^21,22^, suggesting these regions are preferential sites for malignant transformation.

Within this context, we identified a population of argentaffin cells with a distinctive balalaika-like morphology in colonic crypts and subepithelial layers. These NICA cells co-express neuronal, immune, and stemness-associated markers consistent with a neuronal progenitor-like identity and localize near lymphoid follicles, positioning them as potential mediators between epithelial, innate, and adaptive compartments. Although they display features of undifferentiation, their stable morphology indicates lineage consistency, supporting a distinct epithelial cell type. The NICA immune-sensing repertoire renders these cells uniquely vulnerable to pathogens and suitable as a CRC cell of origin. Acting as specialized sentinels, they integrate bacterial transcytosis (GP2), lectin-mediated recognition (ZG16) molecules, TLR4 sensing and antimicrobial effectors (CHGA, Viperin, DEFA6). Crucially, the expression of viral receptors CAR and EPHA2 identifies NICA cells as privileged portals for adenoviruses and gamma-herpesviruses. Supporting this vulnerability, a subset of NICA cells displays a mosaic expression of the adenoviral protein E1B-55K. This suggests that historical or subclinical adenoviral footprints persist within this specific epithelial lineage.

The neuroimmune and stemness phenotypes seen in NICA cells in histological sections are likewise prominent in colorectal cancer tissue and are recapitulated at the gene expression level in isolated tumor crypts. Importantly, in early CRC we observe an expansion of CHGA positive cells, whereas CHGA expression progressively fades in higher stage tumors, coinciding with the upregulation of REST.

The relevance of this shift becomes clearer when placed in the broader context of neuroendocrine-epithelial plasticity. Studies in small-cell lung cancer show a neuroendocrine to epithelial transition driven by the MYC-Notch axis^90^. Some authors have proposed that CRC progression involves lineage plasticity, where tumors partially lose epithelial traits and acquire a neuroendocrine-like phenotype^91,92^. In fact, CRC (and other carcinomas) exhibit the expression of several neuronal markers independently of the clinical stage, suggesting that this neuroepithelial imprint is a constitutive feature of these tumors, as reported in the present work. Molecular profiling demonstrates that MiNENs and NECs remain genetically aligned with conventional adenocarcinomas^93^, consistently carrying “adenocarcinoma-like” mutations in *APC*, *KRAS*, *TP53*, and *BRAF*, supporting a shared cellular origin. REST overexpression contributes to this transition by silencing neuronal genes in non neuronal cells^36^.

REST is strongly upregulated in isolated tumor crypts, coinciding with loss of neuroendocrine transcription factors, functioning as an adaptive mechanism that suppresses neuroendocrine identity and removes neuronal growth constraints. Functional studies reinforce this interpretation. Genetic ablation of REST in CRC models reveals the activation of downstream programs linked to neurobiology and neurodegeneration, as evidenced by both transcriptional profiling and miRNA regulation; furthermore, epigenetic modulation of REST acetylation is shown to reactivate neuronal genes at subtoxic levels. In REST null cells, VPA restores CHGA and TH, indicating reawakening of a silenced lineage. REST loss also alters miRNA profiles, enabling neuroendocrine gene expression and inducing tumor-suppressive miRNAs, positioning REST as a therapeutic target. This is supported by the antitumoral activity of REST modulating agents such as VPA and the EZH2 inhibitor GSK126 in CRC xenograft models^94,95^. High REST expression is a mechanistic driver of poor CRC prognosis, as supported by Human Protein Atlas data. Its upstream activation remains unclear, though oncoviruses such as EBV and KSHV can hijack REST co repressors and silence neuronal genes^96^.

Beyond lineage repression, CRC shows systemic immune remodeling with granulocytosis and reduced B cells, consistent with latent viral drivers such as EBV^97^. CD4 T cells are diminished and overexpress CXCR4, suggesting tumor directed chemotaxis^98,99^, while CD4 and CD8 subsets show CTLA 4 downregulation, indicating impaired checkpoint control and compromised surveillance^100^. This reduced CTLA 4 expression implies that therapies targeting this checkpoint would require prior assessment of it levels, as low expression may limit therapeutic efficacy.

Against this immunological backdrop, we identified a distinct EBV infected B-cell population in tumors with squid-like morphology, termed BLEICS. These immunoglobulin producing cells show dendritic-like signatures suggestive of immaturity or fusion derived hybridization^101^, and co-express immune, neuronal, and stemness markers together with stathmin, an EBV inducible oncoprotein^84^. Their transcriptome displays features of trained immunity^102^, likely reflecting mimicry rather than active infection. BLEICS overexpress CXCR4 and fuse with SDF 1 secreting fibroblasts, promoting fibroblast proliferation and stromal expansion^103^. This fusion is plausibly facilitated by HERV derived envelope proteins with intact fusogenic and immunosuppressive domains^104^, contributing to heterogeneity and immune evasion.

These properties position BLEICS as more than a bystander population. Their presence in colorectal tumors may explain the frequent emergence of lymphoma-like malignancies in xenografted tumors in immunodeficient mice^105^ transplanted with epithelial cancers. Stathmin positive lymphocytes infiltrate early lesions, with subepithelial follicles acting as reservoirs, reinforcing a tumor-follicle axis. In this regard, we observed the fusion of stathmin^+^ BLEICS-like cells with epithelial tumor cells. We identified the fusion of stathmin+ BLEICS-like cells with epithelial tumor cells, a process that significantly accelerates intratumoral heterogeneity. By generating hybrid progeny, this cell-cell fusion acts as a critical driver for enhanced metastatic potential and the emergence of multi-drug resistance^106^. Cancer-immune cell fusion is well documented, including direct evidence of myeloid-epithelial fusion in renal carcinoma after bone marrow transplant^107^, and leukocyte-cancer hybrids have been repeatedly implicated in metastasis^87,88,108^. Hybrids adopting macrophage-like programs gain full migratory and invasive capacity, whereas microglia-like hybrids remain locally confined. This contrast explains the rarity of extracranial glioblastoma metastasis and supports the view that metastatic behavior arises from immune derived phagocytic programs.

Viral structural proteins, like adenoviral E1B-55K or MCV LTAg, directly transactivate HERV loci in p53-mutant CRC cells, unlocking elements with immunosuppressive and fusogenic potential. The detection of E1B-55K in NICA cells indicates that historical adenoviral exposure primes this epithelial lineage, creating a latent retroviral substrate. This substrate is later amplified and reshaped by EBV-infected BLEICS within the tumor niche.

The resulting HERV transcriptome in tumor crypts is non-random, selectively favoring evolutionarily recent elements with intact immunosuppressive (ISD) and fusogenic domains^104^, providing a complementary mechanism for epithelial-stromal fusion and immune evasion within the niche. This functional bias suggests that HERV activation is a co-opted toolkit rather than a neutral byproduct of epigenetic relaxation. By utilizing these envelope fragments, tumor cells and BLEICS actively drive local immunosuppression, epithelial plasticity, and niche remodeling.

A key mechanistic insight from our study reveals that BLEICS directly modulate the retroviral and antiviral landscape of CRC cells, recapitulating the patterns observed within tumor crypts. Their EBV positive status and paracrine activity selectively transactivate HERV-H/F elements, establishing the retroviral signature that dominates the crypt transcriptome. Although EBV can transactivate HERV-K, our data indicate a distinct mechanism in CRC whereby B-cell derived paracrine signals selectively induce HERV-H, a retroelement associated with stem cell-like transcriptional circuits^109^. This provides a mechanistic explanation for the pronounced HERV H upregulation in tumor crypts.

The epithelial response to this retroviral pressure is equally revealing. Tumor crypts display a downstream restricted antiviral imprint characterized by activation of OAS-PKR effectors in the absence of upstream sensors, interferons, or inflammasome components. This pattern rules out acute viral infection and instead reflects a ligand independent antiviral memory -a “viral scar”-imprinted by chronic exposure to immune derived stimuli. The imprint is strongest *in vivo*, partially retained in primary CRC cultures, and largely lost in long-term cell lines, underscoring its dependence on the native microenvironment rather than on retroelement abundance alone.

Co culture experiments recapitulate this architecture. BLEICS mount a complete antiviral program, whereas CRC epithelial cells activate only the sensing arm, failing to engage RNase L and thereby reproducing the ‘execution gap’ observed in tumor crypts. Primary CRC cells retain more of this antiviral competence than commercial lines, highlighting a gradient of immune responsiveness that mirrors the *in vivo* hierarchy. Importantly, HLA DRA induction in epithelial cells is uncoupled from HERV RNA levels and is best explained by paracrine signaling from EBV harboring BLEICS rather than by canonical interferon pathways. Together, these findings position BLEICS as central orchestrators of both retroviral activation and antiviral imprinting in the tumor niche.

Integrating these observations, our study redefines the cellular architecture of colorectal carcinogenesis by identifying two converging drivers of malignant transformation: NICA cells and EBV infected BLEICS. NICA cells emerge as a plausible epithelial cell of origin, given their anatomical position, phenotype, and enrichment in early lesions, prompting a reassessment of the earliest oncogenic events in the colon. In parallel, BLEICS introduce a stromal axis of transformation through their disruptive effects on fibroblasts, echoing fibroproliferative conditions such as keloids and cirrhosis, where aberrant fibroblast expansion and B-cell infiltration correlate with malignant potential^110,111^. Together, these observations support a pathogen first, fusion driven framework that complements and challenges mutation centric models of CRC initiation. In other words, chronic pathogen-induced inflammation is a highly plausible driver of tumorigenesis, as it simultaneously triggers genotoxic mutations and epigenetic silencing to functionally inactivate critical tumor-suppressor genes.

The antiviral behavior of these compartments further reinforces this view. EBV positive BLEICS display escalating antiviral activation with increasing viral activity, whereas epithelial cells progressively lose their antiviral imprint once removed from the tumor niche. This opposing behavior indicates that chronic viral pressure in vivo imprints a transient epithelial ‘viral scar’ that dissipates in vitro when the EBV conditioned microenvironment is lost.

Taken together, our findings support a pathogen-first, fusion-driven model of colorectal carcinogenesis^112,113^ in which NICA cells, EBV infected BLEICS, stromal remodeling, and immune evasion converge as early drivers of malignant transformation. The identification of a niche restricted viral scar strongly supports a hit and run, pathogen conditioned mechanism that cannot be explained by tumor wide HERV upregulation alone. Instead, it reflects a microenvironment sustained at least in part by paracrine pathogen derived cues.

This framework challenges the mutation centric paradigm by positioning viral and retroviral programs as early modulators of canonical oncogenic pathways (Wnt/β-catenin, PI3K/AKT, MAPK/ERK, p53, NF κB), later stabilized by somatic mutations^112,114^. We propose re-evaluating the role of EBV in colonic malignant transformation, particularly its capacity to directly modulate these signaling nodes, where early viral rewiring may cooperate with -and ultimately be “locked in” by-mutational events^112^. By reframing CRC as pathogen mediated lineage disruption and immune-stromal fusion, this model highlights new therapeutic opportunities: targeting viral cofactors, interrupting fusogenic programs, and restoring checkpoint regulation. Even if EBV requires helper viruses or additional oncomodulators to complete transformation, disrupting a single critical link -such as through vaccination^115^ or antiviral strategies-could reduce CRC incidence. This conceptual shift underscores the need to integrate pathogen biology into cancer prevention and therapy, opening translational avenues beyond mutation focused approaches.

## Supporting information

Supplemental information revised

SI BLEICS Phenotype by FACS

SI CIBERSORTx results

SI Cytogenetic II

SI Genetic Mutations Analysis

SI HERVs transcriptome data TC

SI Immune significant genes TC

SI Immune status median and p values

SI Metalloproteinases

SI miRNA in HCT8 REST Null

SI Neuronal transcription factors

SI Neuronal transcription factors

SI Western blots & densitometrics

## Data availability

All data supporting this study as well as primary cells used are reasonably available upon request. Transcriptome data from this study was deposited in NCBI GEO under the accession number GSE303666.

## Acknowledgements

This work was supported by a FoRUM grant from Ruhr University Bochum (AZ F860R 2016) and institutional funds from Marien Hospital Herne. We thank Mrs. Kerstin Heise (Institute of Cell Biology, University of Essen) for assistance with immunoglobulin gene analysis, Mrs. Iris Over (Institute of Human Genetics, Ruhr University Bochum) for karyotyping support and Dr. Ulrik Stervbo (Center for Translational Medicine, Marien Hospital Herne) for help with BLEICS cell phenotyping. We regret any omission of relevant studies. AI based tools were used only for minor editorial assistance; all data and interpretations are author generated.

## Contributions

<u>David Díaz-Carballo</u> (concept, study design and manuscript preparation). <u>Adrien Noa-Bolaño</u>, transcriptome analysis and analysis of HERVs sequences, establishment of primary cell cultures, bioinformatics, ICC, IHC, FACS, WBs, qPCR, and support in article preparation. <u>Udo Rahner</u>, ICC, IHC qPCR, WBs. <u>Ali H. Acikelli</u>, qPCR, WBs, Fontana-Masson staining, bioinformatics, FACS, and support in article preparation. <u>Sahitya Saka</u>, ICC, WBs. <u>Jacqueline Klein</u>, CRISPR/Cas9 experiments design and conduction, IHC, karyotyping, ICC. <u>Sascha Malak</u>, Critical reading. <u>Anne Höppner</u>, qPCR. <u>Annabelle Kamitz</u>, 3D reconstruction of images and graphic representation. <u>Carla Casula</u>, primary cell culture and organoids. <u>Lalitha Kamada</u>, transcriptome analysis, bioinformatics. <u>Andrea Tannapfel</u>, Histopathological analysis. <u>Jens Christmann</u>, NGS, mutation analysis. <u>Sarah Albus</u>, IHC. <u>Enrico Fruth</u>, SEM and TEM. <u>Daniela Gerovska</u>, transcriptome analysis of HERVs, bioinformatics. <u>Marcos J.</u> <u>Araúzo-Bravo</u>, transcriptome analysis of HERVs, bioinformatics. <u>Metin Senkal</u>, sample collection. <u>Crista Ochsenfarth</u>, molecular biology experiments. <u>Sven P. Berres</u> and <u>Julian Uszkoreit</u> have validated the bioinformatic methods of transcriptome analysis, except for HERV analysis. <u>Andrea</u> <u>Tannapfel</u>, histopathological diagnosis. <u>Dirk Strumberg</u>, analysis of the data and approval of the final version.

## Methods

### Ethical Considerations, Patient Material, and Cell Lines

Ethical approval was obtained from the Ethics Committee of the Ruhr University Bochum, Medical Faculty (approval numbers: 15 5416, 17 6114, and 5235 15). All procedures adhered to institutional guidelines and relevant regulations, and written informed consent was secured from every participant. In total, 120 CRC patient samples (peripheral blood and tissue sections) were analyzed. Specimens were anonymized and coded to preserve confidentiality, with access restricted exclusively to authorized research personnel. No participants withdrew during the study. Sex and gender data were not collected, as they were not pertinent to the study objectives. Blinding was not implemented, given that analyses were conducted on anonymized samples with predefined gating strategies.

### Colorectal Carcinoma Cell Lines

The colorectal carcinoma cell lines HCT8 (HRT-18, CVCL 2478), HT29 (HTB-38, CVCL 0320), and Caco2 (HTB-37, CVCL 0025) were obtained from the tumor bank of the Medical Faculty of the University of Duisburg-Essen. Cell lines were authenticated by the cell bank using standard short tandem repeat (STR) profiling. Cells were maintained in Dulbecco’s Modified Eagle Medium (DMEM; PAN Biotech, Aidenbach, Germany) supplemented with 10% heat-inactivated fetal calf serum (FCS) and 3 µg/mL doxycycline. Cultures were incubated at 37 °C in a humidified atmosphere containing 5% CO. Cells were routinely tested for mycoplasma contamination using the MycoAlert Mycoplasma Detection Kit (Biozym, Hessisch Oldendorf, Germany), according to the manufacturer’s instructions.

### Isolation of Primary Colorectal Crypts, Carcinoma Cells, and BLEICS Cells

Colorectal tissue samples from tumor and adjacent normal regions of the same patients were freshly collected immediately following surgical resection. Specimens were washed three times with pre-chilled Hank’s Balanced Salt Solution (HBSS) supplemented with 1% penicillin/streptomycin/amphotericin B (Pan Biotech, Aidenbach, Germany), 25 µg/mL gentamycin, 25 µg/mL fluconazole, and 50 µg/mL metronidazole. The lamina propria mucosae was mechanically separated from the underlying submucosa using sterile forceps and fine scissors.

The epithelial mucosa was rinsed twice in fresh, pre-chilled HBSS and transferred into 50 mL conical tubes containing 25 mL of an enzymatic digestion cocktail composed of 1 mg/mL each of collagenase I, collagenase II, and dispase I in 1× HBSS. Tissue was finely minced within the enzymatic solution and incubated at 37 °C for 30 minutes on an end-over-end rotator. After digestion, samples were centrifuged at 500 × g for 10 minutes. Pellets were resuspended in fresh enzymatic cocktail and subjected to a second 30-minute incubation under identical conditions. The resulting cell suspensions were filtered through 500-µm mesh strainers and centrifuged again at 500 × g for 4 minutes at room temperature.

Crude isolates of regular crypts (RCs) and tumor-derived crypts (TCs) were recovered and washed three times in DMEM/F12 medium supplemented with 10% FCS and the same antibiotic/antimycotic cocktail (excluding fluconazole). Crypts were either cultured in DMEM/F12 medium containing 20% FCS and antibiotics to establish primary colorectal carcinoma cultures, or washed with PBS and immediately processed for RNA extraction and transcriptomic analysis.

Following isolation, tumor-derived crypts were cultured to promote expansion of epithelial cells. During early culture stages, non-epithelial cells (e.g., fibroblasts and immune cells) were gradually depleted through selective medium changes and wash steps. Upon passaging, epithelial tumor cells were preferentially retained due to their strong adherence and colony-forming properties, while loosely attached contaminating cells were removed. Continued passaging enriched the epithelial fraction, leading to the establishment of stable primary colorectal carcinoma cell populations. Independently adhering fibroblasts that proliferated during this process were expanded separately for use in co-culture and mechanistic assays.

### Characterization of Primary Colorectal Carcinoma Cell Lines and BLEICS Cells

In addition to commercial colorectal carcinoma cell lines, two primary tumor-derived cell lines -Col33 and Col124-were selected for further experimental analysis based on their distinct biological characteristics. Col33 is a rapidly proliferating culture capable of forming tubular, crypt-like structures in two-dimensional culture systems. Notably, these cells exhibit a radial organization centered on a distinctive polarity axis (see SI: CRC primary cells morphology and cellular organization). In contrast, Col124 is a slow-growing culture characterized by robust expression of E-Cadherin 17 and resistance to detachment by standard enzymatic dissociation methods.

BLEICS cells (B-lymphocyte, EBV-Infected, Calamari-Shaped) emerged spontaneously in fibroblast cultures within 5-10 days post-tumor isolation. These cells were maintained in DMEM/F12 medium supplemented with 10% fetal calf serum and 3 µg/mL penicillin/streptomycin/amphotericin B (PSA). BLEICS cells remained in suspension and were readily collected by harvesting the culture supernatant.

### Immunotechniques

All immunological techniques -including Western blotting, immunocytochemistry (ICC), Immunohistochemistry (IHC), and flow cytometry-were performed according to standard protocols as previously described^86,116^. To ensure the specificity of antibody binding, appropriate negative controls were included in every immunological assay. Specifically, isotype-matched controls were used for all primary antibodies, and corresponding secondary antibody-only controls were performed for immunocytochemistry (ICC), immunohistochemistry (IHC), and fluorescence-activated cell sorting (FACS) experiments.

To strengthen the validity of our findings, we confronted our results with external public repositories, including Human Protein Atlas, thereby providing an independent reference framework for the observed expression patterns.

### Western Blotting

Cells in the exponential growth phase were harvested for protein extraction. Monolayers were washed with PBS and detached using 0.05% trypsin (PAN Biotech, Aidenbach, Germany). The resulting cell suspensions were neutralized with culture medium and centrifuged at 300 × g for 5 minutes. Cell pellets were washed in cold PBS, centrifuged, and lysed in RIPA buffer (150 mM NaCl, 1 mM EDTA, 1% Triton X-100, 1% sodium deoxycholate, 0.1% SDS, and 50 mM Tris-HCl, pH 7.4) supplemented with a protease inhibitor cocktail (Roche Diagnostics GmbH, Basel, Switzerland). Lysis was carried out on ice for 30 minutes, followed by centrifugation at 14,000 × g at 4 °C for 20 minutes. Supernatants were collected, and protein concentrations determined using the Bradford assay.

Thirty micrograms of total protein per sample were resolved by SDS-PAGE using a 4-20% gradient gel (Biotrend, Cologne, Germany) in Tris-glycine buffer (0.025 M Tris-HCl, 0.192 M glycine, pH 8.5), then transferred onto 0.2 µm nitrocellulose membranes (Bio-Rad Laboratories, CA, USA). Membranes were blocked with 10% non-fat dry milk in PBST (0.05% Tween-20 in 1× PBS), washed, and incubated overnight at 4 °C with primary antibodies diluted in 1.25% non-fat milk in PBST, as recommended by the manufacturers (SI: Table 3: Antibodies). After three PBST washes, membranes were incubated with horseradish peroxidase-conjugated secondary antibodies diluted in 2.5% non-fat milk in PBST for 2 hours at room temperature. Immunoreactivity was visualized using Western Lightning Plus-ECL (PerkinElmer) and imaged using the ChemiDoc XRS+ system equipped with Image Lab Version 2.0.1 software (Bio-Rad Laboratories).

### Immunocytochemistry (ICC)

Cells were cultured in chamber slides to the appropriate density, washed with 1× PBS, fixed with 4% formaldehyde in PBS for 20 minutes, and stored at -20 °C until use. Prior to staining, slides were thawed, rinsed twice with PBS (5 minutes each), permeabilized with 0.1% Triton X-100 in PBS, and blocked for 2 hours at room temperature with 10% bovine serum albumin (BSA; AbD Serotec, London, UK) and 1:500 human FcR Blocking Reagent (Miltenyi Biotec, Gladbach, Germany) in PBST. Primary antibodies (SI: Table 3 Antibodies), diluted in 1% BSA in PBST, were applied overnight at 4 °C.

Following three PBST washes, cells were incubated with secondary antibodies diluted in 1% BSA in PBST for 2 hours at room temperature in the dark. After an additional PBST wash (5 minutes), cell nuclei were counterstained with Hoechst 33258 (1 µg/mL in PBST) for 15 minutes. Slides were rinsed three times in PBST and mounted using Faramount Mounting Medium (Agilent Technologies, CA, USA). Samples were visualized using an Eclipse i-50 microscope (Nikon, Tokyo, Japan) and analyzed with NIS-Elements Advanced Research imaging software.

### Immunohistochemistry (IHC)

Paraffin-embedded tissue sections (5 µm thickness) were baked overnight at 60 °C to ensure firm adhesion to microscope slides and to eliminate residual paraffin. Sections were deparaffinized by two 20-minute incubations in xylene substitute (Thermo Scientific, London, UK), followed by rehydration in a graded ethanol series (3×100%, 95%, 70%, and 50%; 25%, dH_2_O, PBS, 5 minutes each). Antigen retrieval was performed by heating the sections in 10 mM sodium citrate buffer (pH 6.0; Sigma-Aldrich, Taufkirchen, Germany) at 95 °C for 30 minutes using a domestic vegetable steamer. After cooling, slides were washed twice with 1× PBS for 5 minutes and blocked for 2 hours at room temperature in 5% BSA (AbD Serotec, London, UK) and 1:500 human FcR Blocking Reagent (Miltenyi Biotec, Gladbach, Germany) prepared in PBST. Primary antibodies (SI: Table 3: Antibodies) were diluted in 1% BSA in PBST according to the manufacturers’ specifications and applied overnight at 4 °C. The following day, slides were washed three times with PBST (5 minutes each), and tissue sections were incubated for 2 hours at room temperature with conjugated secondary antibodies (Cell Signaling, Leiden, Netherlands) diluted in 1% BSA in PBST, following the manufacturers’ instructions. Nuclear staining with Hoechst 33258, mounting, and imaging were performed as described in the Immunocytochemistry section.

### Peripheral Blood Collection and Flow Cytometry Analysis

Peripheral blood samples were collected from 102 patients diagnosed with colorectal cancers (mean age: 70 years), as well as from two control groups: young healthy individuals (<30 years) and elderly healthy donors (mean age: 82 years). To investigate systemic immune alterations, particularly in relation to BLEICS cell involvement, comprehensive flow cytometric profiling of peripheral blood leukocytes was performed. All cytometric parameters, including antibody panels, fluorochrome combinations, and gating strategies, were standardized across all study groups. Reagents and protocols were applied uniformly to ensure comparability of results. For immune phenotyping, flow cytometric analyses were performed using established protocols. In brief, 50 µL of EDTA-anticoagulated whole blood was incubated for 20 minutes with fluorochrome-conjugated antibodies (SI: Table 3: Antibodies - Coupled FACS antibodies). Following staining, samples were diluted to a final volume of 500 µL with VersaLyse solution (Beckman Coulter, Krefeld, Germany) and incubated at room temperature for 5 minutes to lyse erythrocytes. Appropriate isotype-matched control antibodies (SI: Table 3: Antibodies) were employed for signal gating. Labeled cells were analyzed using a CytoFLEX Research Cytometer B5-R5-V5 (Beckman Coulter, Krefeld, Germany). For each sample, a minimum of 2 × 10 events were acquired to ensure sufficient statistical power. The main cell population was identified based on forward and side scatter (FSC/SSC) dot plot density, allowing exclusion of dead cells. Isotype control signals were used to define the threshold for negative staining.

### Histochemistry

Fontana-Masson staining was performed for the histological detection of argentaffin cells using a validated kit (ScyTek, Utah, USA), following the manufacturer’s protocol. After deparaffinization, tissue sections were immersed in silver nitrate solution pre-equilibrated with ammonium hydroxide and incubated at 58 °C until they developed a yellowish-brown coloration, typically after 30 minutes.

Slides were then rinsed three times with distilled water and treated with 0.2% gold chloride solution for 30 seconds. Following another three rinses in distilled water, sections were incubated in 5% sodium thiosulfate solution for 2 minutes to stabilize silver deposits. After additional rinsing, tissues were counterstained with Fast Red solution for 5 minutes and subsequently mounted using Eukitt® mounting resin.

### Transmission Electron Microscopy (TEM) and Three-Dimensional Reconstruction

BLEICS cells were fixed in 2.5% glutaraldehyde (GA) in PBS for 3 hours at room temperature, washed three times for 20 minutes each with PBS, and post-fixed in 1% osmium tetroxide (pH 7.4) for 45 minutes. Residual osmium tetroxide was removed by three additional PBS washes (3 × 20 min). Samples were then pre-dehydrated in 30% ethanol (3 × 20 min) and subjected to a graded ethanol dehydration series (30%-100%), 5 minutes per step. Dehydrated samples were incubated in a 1:1 (v/v) mixture of propylene oxide and ethanol for 15 minutes, embedded in pure EPON®, followed by a 1:1 (v/v) mixture of EPON® and Araldite for 15 minutes, and polymerized by desiccation at 80 °C for 48 hours. Resin-embedded blocks were sectioned into 60 nm ultrathin slices using a Leica Ultracut UCT ultramicrotome (Leica, Wetzlar, Germany). Sections were mounted on 200-mesh hexagonal copper grids (Leica) and contrasted using a Leica EM AC20 system with 0.5% aqueous uranyl acetate for 45 minutes at room temperature. Grids were further stained with 3% lead citrate and rinsed with Milli-Q water for 10 minutes. Samples were examined using a JEOL 2100 transmission electron microscope (JEOL, Tokyo, Japan) operating at 120 kV. Images were captured at magnifications ranging from 3 000× to 140 000× using ITEM® 5.0 software (Soft Imaging Systems, Cardiff, UK). For three-dimensional reconstruction, semi-thick (0.5 µm) sections were generated using the same ultramicrotome and stained with 1% toluidine blue, azure II, methylene blue, and sodium tetraborate in distilled water. Sections were mounted on glass slides, air-dried, and cover slipped with a xylene substitute. Serial images from 50 consecutive sections were acquired at 1 000× magnification on an Eclipse i-50 microscope (Nikon, Tokyo, Japan), aligned using Adobe Photoshop CS4, and processed into animated reconstructions using JASC Animation Shop 3.01.

### Scanning Electron Microscopy (SEM)

BLEICS cells co-cultured with cancer-associated fibroblasts (CAFs) were seeded on Thermanox coverslips and grown to appropriate densities. After medium removal, samples were washed three times with PBS and fixed with 2.5% GA in PBS for 2 hours at room temperature. Fixed specimens were rinsed three times with Milli-Q water (15 minutes each), dehydrated via a graded ethanol series (10%-100%, 5 minutes per step), and air-dried in a fume hood. After sputter-coating with gold, samples were imaged using a Zeiss DSM 982 scanning electron microscope equipped with Gemini optics and the Digital Image Scanning System (DISS5; Point Electronic, Halle, Germany).

### Time-Lapse Videography

To investigate the fusogenic capacity of the newly established BLEICS cells (B-Lymphocyte, EBV-Infected, Calamari-Shaped cells), time-lapse videography was employed. Crucially, these experiments were conducted exclusively using BLEICS cells cultured alongside primary colorectal fibroblasts (Cancer-Associated Fibroblasts, CAFs). This setup was designed to demonstrate the cell line’s ability to engage and fuse with supporting stromal elements, as evidenced by the transfer of cytoplasmic material and subsequent morphological changes.

Live-cell imaging was performed using a TE2000 inverted microscope (Nikon, Tokyo, Japan) equipped with a PeCom environmental incubation system (5% CO, 37 °C) to maintain physiological culture conditions. Time-lapse image acquisition and processing were carried out using NIS-Elements BR software (Nikon Vision Systems).

### Total RNA Isolation and RT-qPCR

Cells were cultured in DMEM/F12 medium supplemented with 10% FCS (or 20% FCS for primary cultures) and harvested at approximately 70% confluency. Total RNA was extracted using TRIzol® reagent (Life Technologies, California, USA), followed by DNase I treatment (7 Kunitz units; Qiagen, Hilden, Germany) to remove genomic DNA contamination. RNA samples were further purified using RNeasy Mini columns (Qiagen) according to the manufacturer’s protocol. RNA integrity was assessed using a 2100 Bioanalyzer (Agilent Technologies, Santa Clara, CA, USA).

Neuronal marker transcripts were quantified via RT-qPCR using custom-designed primers and probes (SI: Table 4: qPCR primers and probes), synthesized by IDT Inc. (Iowa, USA). Amplification of 100 ng of cDNA per reaction was performed in triplicate using 250 nmol primers and PrimeTime® Gene Expression Master Mix (IDT) on a CFX96™ Real-Time PCR Detection System (Bio-Rad Laboratories, CA, USA). Relative gene expression levels were calculated using the comparative ΔCt method. For downstream analysis, only transcripts with ΔCt values below 29 cycles were considered reliably expressed.

### Transcriptome Analysis

Total RNA (0.5 µg) was extracted from colorectal carcinoma cell lines (HCT8, HT29, CaCo2, Col33, and Col124), regular and tumor-derived crypts from three individual patients, BLEICS21 and BLEICS126 cells, the B-lymphoma cell line Namalwa, and peripheral B cells isolated from buffy coats of ten healthy donors. RNA samples were subjected to next-generation sequencing (NGS) on the Illumina platform (BGI Genomics, Hong Kong, China). Transcriptome data from this study was deposited in NCBI GEO under the accession number GSE303666. Initial transcriptomic profiling was performed using the Dr. Tom platform (BGI Genomics), followed by additional bioinformatic analyses. Differential gene expression analysis was conducted using the DESeq2 package (v1.42.1), and gene set enrichment analysis (GSEA) was performed using the GSEA 4.3.3 desktop application (Broad Institute). Protein-protein interaction (PPI) networks and enrichment clusters were explored via the STRING database, while immune cell-type deconvolution was carried out using CIBERSORTx with the LM22 signature matrix. Transcriptome expression levels were quantified as transcripts per million (TPM). Differential expression was defined using log2 fold change thresholds of 1.5-2.0 for both upregulated and downregulated genes, with a false discovery rate (FDR) < 0.05. Statistical significance was assessed at α = 0.05 or 0.01, as specified. For data visualization and enrichment mapping, Cytoscape was employed in conjunction with the EnrichmentMap plugin and associated analytic tools. To investigate the expression of human endogenous retroviruses (HERVs), the genomic coordinates of all transcriptional peaks were intersected with a curated reference set comprising ≈3 220 individual HERV loci annotated in the human genome. These loci were previously characterized using the ERVmap database, which provides high-resolution annotations of HERV family classifications, genomic positions, and transcriptional activity^117^.

### Metagenomic Analysis of Crypt-Associated Pathogens Using Whole-Transcriptome Sequencing

Whole transcriptome profiles were generated from crypts dissected from colorectal carcinoma tissue and from matched non tumoral mucosa located approximately 3 cm from the tumor margin. All samples were obtained from the same individuals, enabling controlled intra patient comparisons. Crypt isolation included multiple sequential washes, effectively removing loosely adherent microorganisms and enriching for pathogens firmly attached to the crypt surface or retained within the crypt lumen. Non human reads from each transcriptome were analyzed using the Kranke2 metagenomic pipeline to identify and quantify pathogen associated transcriptional signatures. Comparative analyses were performed within and across the three patients studied, first to identify pathogens uniquely present in tumor derived crypts and absent from normal crypts, and subsequently to determine taxa that, while present in both compartments, were enriched in tumor-associated crypts.

### CRISPR/Cas9 Ablation of Genes Encoding RE1-Silencing Transcription Factor (REST)

CRISPR/Cas9 (Clustered Regularly Interspaced Short Palindromic Repeats/CRISPR-associated protein) technology was utilized to knock out all variants of the *REST* gene (NM_005612.5). The Alt-R™ CRISPR/Cas9 System and HPRT gRNA controls (IDT Technologies, Leuven, Belgium) were employed according to the manufacturer’s recommendations. crRNAs targeting REST were specifically designed using the IDT CRISPR design tool (https://eu.idtdna.com) and synthesized by the same provider (SI: REST CRISPR/Cas9 ablation). Guide RNA (gRNA) sequences were aligned to the human genome using the Basic Local Alignment Search Tool (BLAST) to assess potential off-target effects. The crRNA:tracrRNA complexes were prepared at equimolar concentrations in Nuclease-Free Duplex buffer (IDT), incubated at 95 °C for 5 minutes, and subsequently cooled to room temperature.

Ribonucleoprotein (RNP) complexes were formed following IDT protocols by incubating 0.1 µM of the crRNA:tracrRNA complex with 1 U of Alt-R® S.p. HiFi Cas9 Nuclease V3 (IDT Technologies) for 20 minutes at room temperature. The resulting RNPs were immediately transfected into HCT8^WT^ cells using the 4D-Nucleofector™ System (Lonza), applying pulse code EH-100. Genome editing was confirmed using the EnGen™ Mutation Detection Kit (New England BioLabs, MA, USA). Genomic DNA was isolated with the NucleoSpin DNA RapidLyse Kit (Macherey-Nagel). Amplicons were generated via endpoint PCR using control primers. Wild-type and edited PCR products underwent heteroduplex formation (denaturation at 95 °C for 5 minutes, followed by annealing from 95 °C to 25 °C at a ramp rate of 0.2 °C/s). Heteroduplexes were then treated with EnGen T7 Endonuclease I at 37 °C for 15 minutes. DNA fragments were separated on a 2.0% agarose gel, stained with ethidium bromide, and visualized using the ChemiDoc™ XRS+ Imaging System (Bio-Rad, CA, USA). CRISPR/Cas9-edited cells were cloned using limiting dilution. Cells were diluted to a concentration of 100 cells/mL and seeded into 96-well plates at one cell per well, with single-cell occupancy confirmed by field microscopy. Wells containing a single cell that exhibited growth were selected, and the cells were recovered by trypsinization (100 µL). These were subsequently transferred to 24-well plates for expansion and then to 25 cm² flasks for further cultivation and expansion. Clonal populations were analyzed by qPCR to assess changes in REST mRNA expression, and REST protein levels were evaluated by Western blotting. Clones lacking REST expression were considered successfully mutated and selected for downstream analyses.

### Generation of Organoids from Colorectal Carcinoma Cells

Organoids are three-dimensional cellular assemblies capable of self-organization and self-renewal, designed to replicate the complex growth patterns and cellular diversity of tumors. To generate organoids from colorectal carcinoma (CRC) cells, we followed a previously described protocol with minor modifications^86^. Colorectal carcinoma (CRC) cells were embedded in the basal membrane matrix Geltrex™ (Thermo Fisher Scientific, MA, USA), forming domes containing 3 000 cells in 10 µL volumes on culture slides. The embedded cells were subsequently maintained in organoid medium (SI: Table 5: Colonic Organoid Medium), with media changes performed every three days. Organoids were used approximately one week after initiation to evaluate the impact of REST ablation and the effects of REST inhibitors on their growth.

### Epigenetic Modulation of the Transcription Factor REST in CRC Cells Using Specific Inhibitors

To induce a semi-neuronal phenotype in colorectal cancer (CRC) cells and thereby explore latent lineage characteristics, the HCT8 cell line and Col33 CRC cells were treated with subtoxic concentrations of key epigenetic modulators. The cytotoxicity of the REST inhibitor drugs X5050 and GSK 126 was initially assessed in organoids derived from Col33 cells (see SI: Toxicity of REST inhibitors X5050 and GSK126). Valproic acid was excluded from testing due to its well-documented low toxicity in culture, showing no impact on cell growth at concentrations up to 100 µg/mL. The experimental aim was to partially reverse oncogenic epigenetic reprogramming and assess transcriptional changes linked to neural gene regulation. Cells were incubated with the histone deacetylase (HDAC) inhibitor valproic acid (30 µg/mL) and the EZH2 methyltransferase inhibitor GSK126 (10 µg/mL) for 72 hours. Valproic acid was selected due to its well-established ability to alter chromatin accessibility by inhibiting HDAC activity, thereby facilitating the expression of silenced neuronal genes. GSK126 targets EZH2 -the catalytic component of the polycomb repressive complex 2 (PRC2)-which catalyzes trimethylation of histone H3 on lysine 27 (H3K27me3), a mark associated with gene silencing. Inhibiting EZH2 can derepress genes involved in neuronal differentiation, including targets of REST.

Following treatment, total RNA was extracted according to previously described protocols. Gene expression analysis was performed, focusing on the expression levels of REST-regulated and neurogenesis-related genes. This strategy allowed evaluation of how epigenetic remodeling affects REST functionality and downstream gene networks, potentially identifying regulatory mechanisms relevant to CRC differentiation and therapeutic susceptibility.

### Statistic and Reproducibility

Statistical analysis of immune profiling data was performed using paired, two-tailed Student’s *t*-tests, with significance defined as *p* < 0.05. Graphical representation and statistical computations were conducted using Microsoft Excel 2016 and OriginLab 2022 (v9.9.0.225; OriginLab Corp., MA, USA). Western blot quantification was carried out by densitometric analysis, with signal intensities normalized to housekeeping protein levels.

Immunocytochemistry (ICC) and immunohistochemistry (IHC) data were evaluated qualitatively for descriptive purposes and were not subjected to statistical testing. All experimental results reflect findings from at least three independent biological replicates.

