## Supplemental information revised for "Neuro-Immune Crypt-Associated Cells and REST-Mediated Reprogramming: Pathogen-Driven Stromal Activation, HERVs Induction, and Abortive Antiviral Signaling in Colorectal Carcinoma"

**SI: DCLK1 in subepithelial colon**

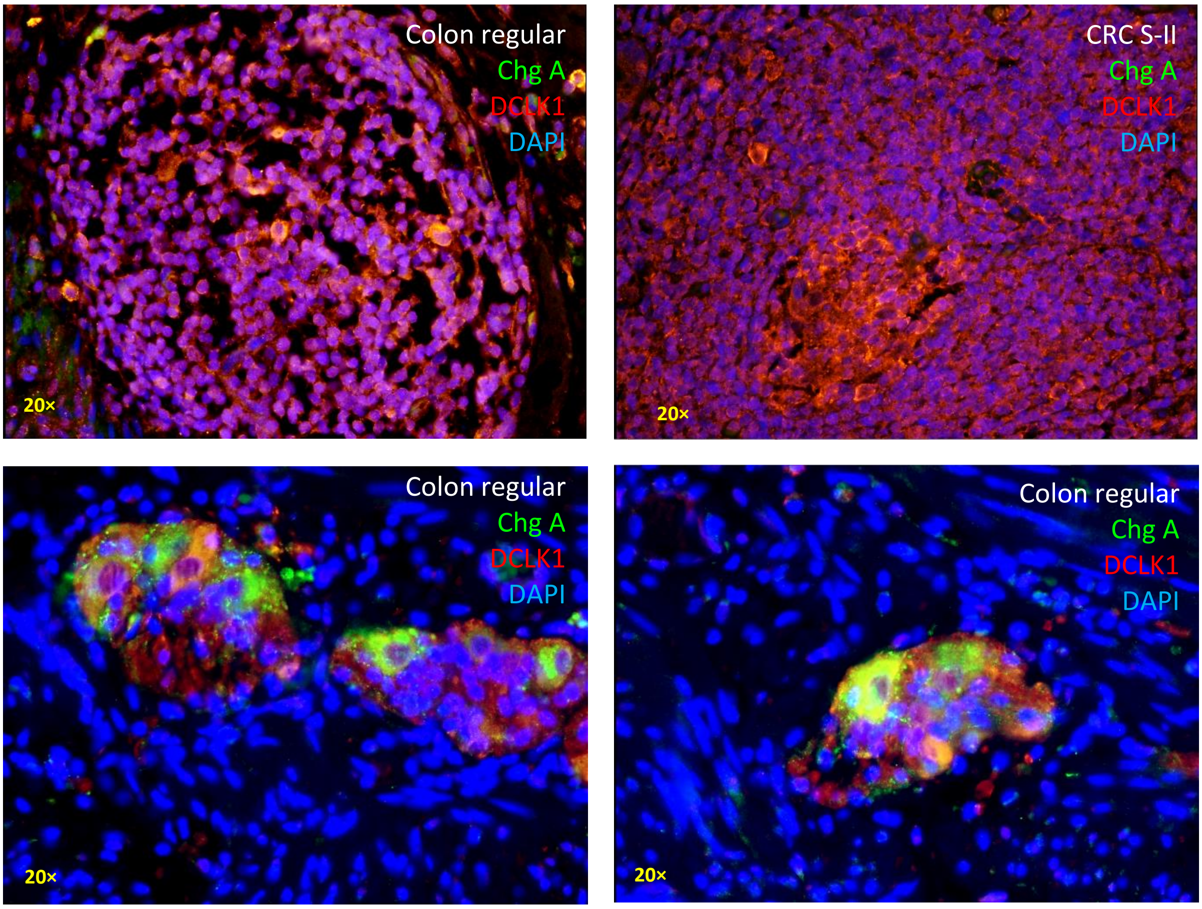

**SI: NICA basolateral Pocket**

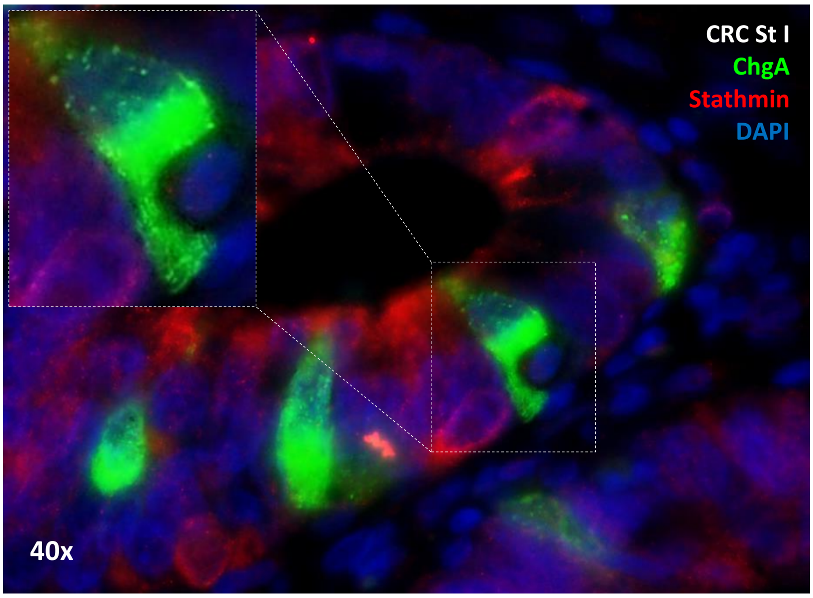


**Gp2 produced by NICAS is emptied into the crypt's lumen covering the epithelial surface**

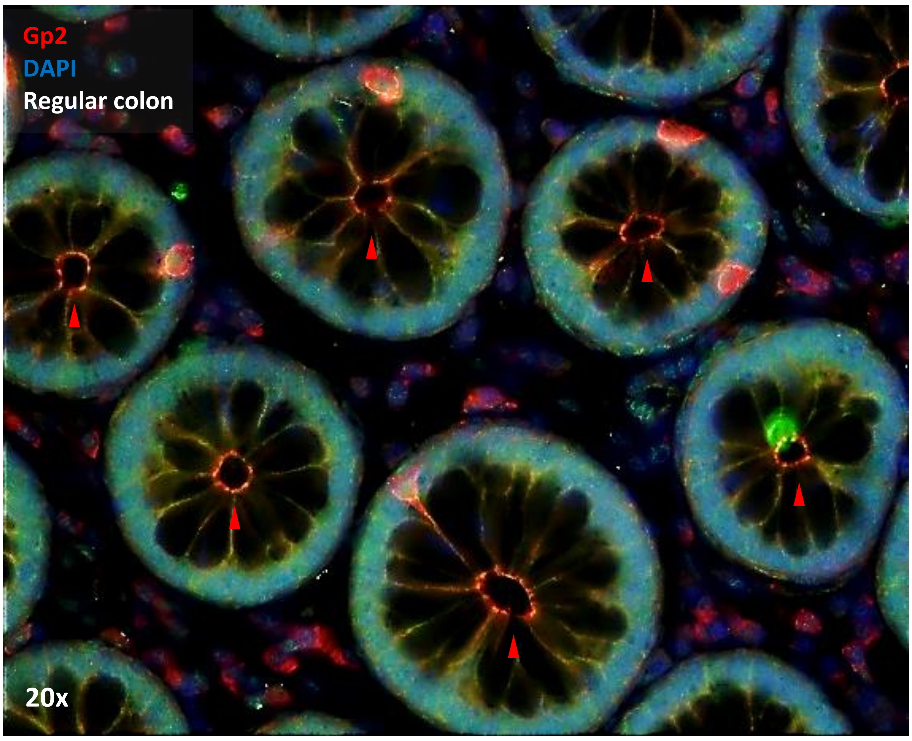

### SI: CAR and EB1-K55 distribution in NICA and colon tissues

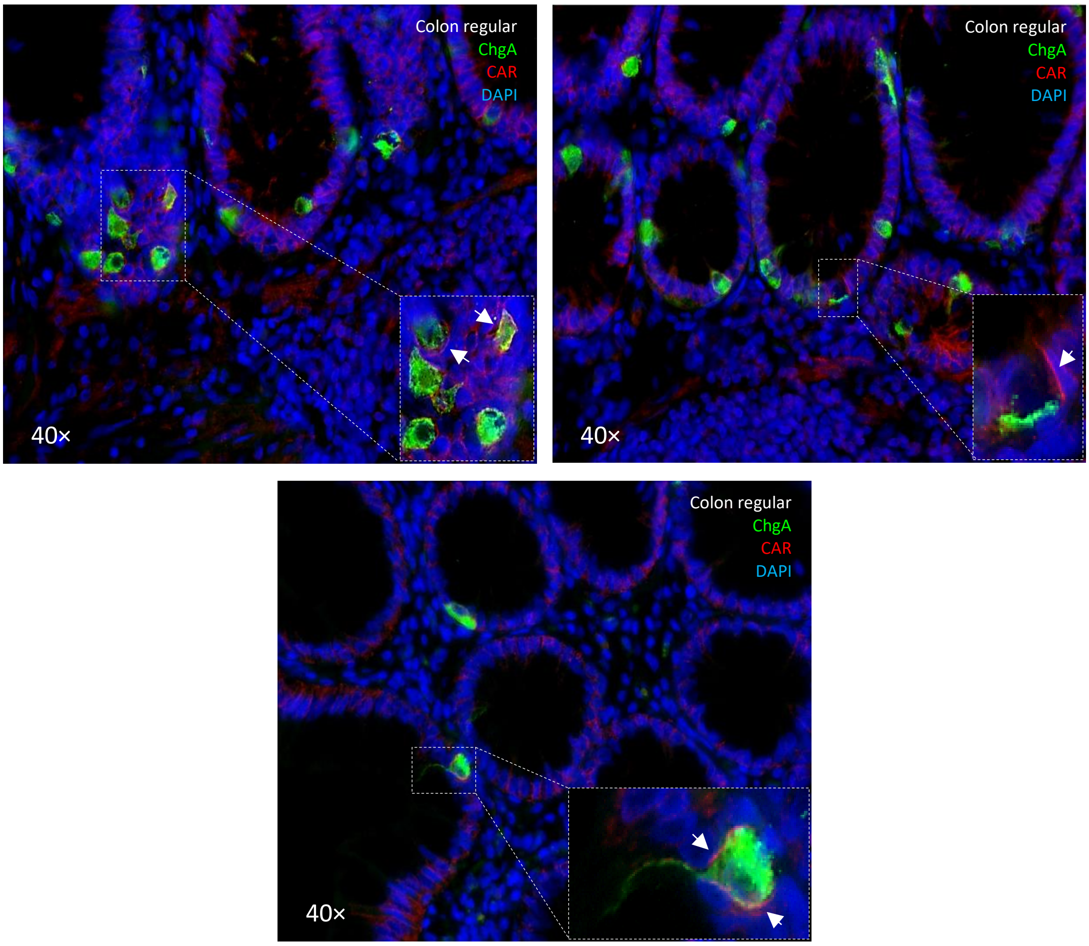


**<u>Panel A</u>:** Expression of the adenovirus type 5 receptor CAR in human colon epithelium

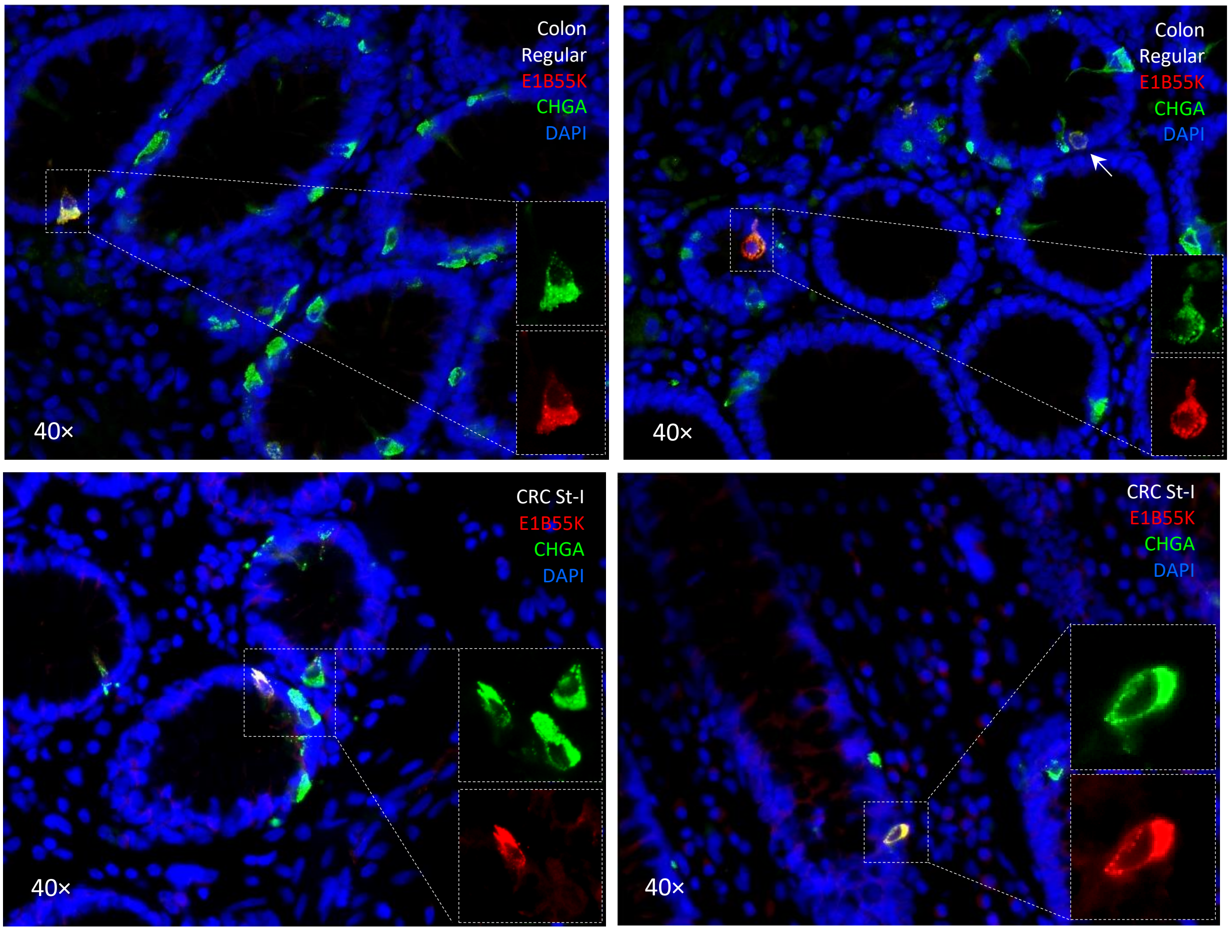


**Panel B**: E1B55K Distribution in Neuroimmune Crypt-Associated (NICA) Cells of the Colon

**Panel A: Expression of the adenovirus type 5 receptor CAR in human colon epithelium**. IHC analysis reveals that CAR is expressed in the majority of crypt epithelial cells, as well as in immune cells located within lymphatic follicles and those patrolling the lamina propria. The CAR receptor shows a polarized distribution, with expression predominantly enriched at the basolateral membrane, consistent with its known localization at epithelial junctions. Notably, $ChgA^+$ neuroimmune crypt-associated (NICA) cells exhibit basolateral CAR expression, with intensity slightly exceeding that of surrounding crypt epithelium. This selective enrichment positions NICA cells as potential viral entry points for adenovirus type 5 within the colonic mucosa. The co-localization of CAR with ChgA confirms receptor expression in the neuroimmune lineage, highlighting a novel intersection between epithelial polarity, immune surveillance, and viral tropism. **Panel B: E1B55K Distribution in Neuroimmune Crypt-Associated (NICA) Cells of the Colon.** At 40x magnification, the immunofluorescence imaging reveals a distinct mosaic distribution of the E1B55K protein (red) within the CHGA+ (green) enteroendocrine cell population. While the DAPI (blue) nuclear staining outlines the crypt architecture, the merged yellow signal confirms that E1B55K expression is restricted to a specific subset of secretory cells. Notably, the protein is not universally present; many CHGA-positive cells do not express E1B55K, while only a selected few show clear co-localization. This heterogeneous pattern remains consistent across both regular colon and Stage I Colorectal Cancer (CRC) tissues. Representative for at least 10 individuals.

| HERV ID | -LOG$_{10}$ (PVAL) | ΔTPM | SUBGENES | ISD | BESTREFRV | SUPER GROUP |
|---|---|---|---|---|---|---|
| 3542 | 1.8 | 1.1 | 5LTR PBS CA NC Prot RT IN SU TM PPT 3LTR | YQNRLvLDYLLv | Human endogenous retroviral DNA (4-1), complete re 92% -98 | HERVERI |
| 3593 | 1.7 | 2.4 | 5LTR CA NC RT IN SU TM PPT 3LTR | RQTVtWMGDkIMSLeHrLQMQC | Human endogenous retrovirus HERV-K10. 60% -2255 | HML |
| 6073 | 2 | 1 | 5LTR PBS IN SU TM PPT 3LTR | RQTVIWMrDRIiSLEHRLQmqC | Human endogenous retrovirus HERV-K(I) DNA, complete 64% -2205 | HML |
| 1805 | 1.5 | 1.9 | CA NC RT IN SU TM PPT 3LTR | LRQTVIWMGDRI | Human endogenous retrovirus HERV-K10. 61% -2110 | HML |
| 2048 | 1.6 | 1.1 | NC Prot RT SU TM PPT | LRQSVIWLGDwV | Homo sapiens tandemly repeated human endogenous re 62% -3890 | HML |
| 704 | 1.7 | 2.1 | 5LTR PBS NC Prot RT IN TM PPT 3LTR | LQNzmALnivTAAzGGTCAlLG | | HSERVIII |
| 510 | 1.8 | 1 | 5LTR PBS CA NC Prot RT IN SU TM 3LTR | LQNRwGLDLimAEKRdLCLsLG | Homo sapiens RGH2 gene, retrovirus-like element. 57% -1199 | HERVHF |
| 3165 | 2.5 | 3.9 | CA Prot RT IN TM 3LTR | LQNRRGLDLLtTEKGGsCLsLG | Homo sapiens human endogenous rv HML6 gag,pol,env 50% 307 | HERVHF |
| 906 | 1.3 | 1.2 | 5LTR PBS MA CA NC Prot RT IN SU TM PPT 3LTR | LQNRqsLDLLTAEKGGLCIFLN | Homo sapiens RGH2 gene, retrovirus-like element. 83% -22 | HERVHF |
| 5480 | 2.4 | 2 | 5LTR PBS MA CA NC Prot RT IN SU TM PPT 3LTR | LQNRqGLnLLTAEKGGLCIFLN | Homo sapiens RGH2 gene, retrovirus-like element. 80% -48 | HERVHF |
| 765 | 1.7 | 1.1 | 5LTR PBS NC Prot RT IN TM PPT 3LTR | LQNqmALDiiTtARGGTCslLG | | HSERVIII |
| 4475 | 2.3 | 4.4 | 5LTR PBS MA CA NC Prot RT IN TM PPT 3LTR | LQNLqGLDLLTAEERgLCIFLN | Homo sapiens RGH2 gene, retrovirus-like element. 73% -463 | HERVHF |
| 2518 | 1.3 | 1.1 | CA NC Prot RT IN TM PPT | LQNhRGLDLLTvEKGGLCtFLG | Human endogenous retrovirus, complete genome. 68% -1409 | HERVHF |
| 3698 | 2.8 | 1.1 | IN SU TM PPT 3LTR | LQNhRGLDLLTAEKGGLCIFLN | Homo sapiens RGH2 gene, retrovirus-like element. 77% -4627 | HERVHF |
| 4489 | 2.5 | 1.2 | 5LTR PBS MA CA NC Prot RT IN SU TM 3LTR | LQNgqGrDLLTAEKGGLCIFLN | Human endogenous retrovirus, complete genome. 84% -159 | HERVHF |
| 3016 | 1.5 | 1.9 | 5LTR PBS CA NC Prot RT IN SU TM PPT 3LTR | LQNcqGLDmLmAAQGGICLALd | | HERVFRDLIKE |
| 2643 | 1.6 | -1.4 | 5LTR PBS CA IN TM PPT 3LTR | YQNRLALDYLLA | Human endogenous retroviral DNA (4-1), complete re 68% -1020 | HERVERI |
| 868 | 2.4 | -2 | 5LTR PBS MA CA NC Prot RT IN TM PPT 3LTR | wgNRiALDmLLA | Homo sapiens human endogenous retrovirus HERV-P4.6 73% -177 | HERVIPADP |
| 2124 | 1.5 | -2.5 | 5LTR PBS MA CA NC IN TM | weNRiALniiLA | Homo sapiens human endogenous retrovirus HERV-P4.6 75% -784 | HERVIPADP |
| 2754 | 2.8 | -1.2 | CA NC Prot RT SU TM PPT | LQNRzGLDLLTAEKGGLCtFLG | Human endogenous retrovirus pHE.1 (ERV9). 85% -503 | HERVW9 |
| 2674 | 2.4 | -2.5 | Prot IN SU TM PPT | hQTVpzMGDzIiNLEHRiQIQC | Human endogenous retrovirus HERV-K(II) DNA, comple 64% -5004 | HML |

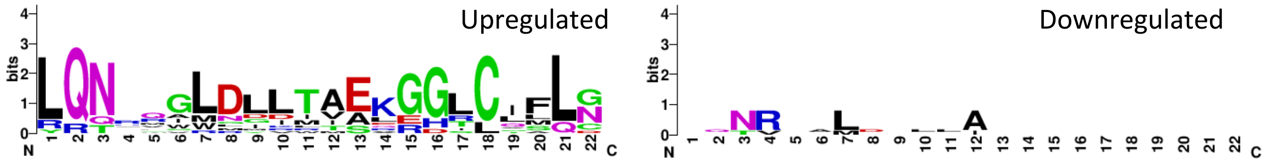


**Table 1:** Differentially expressed HERVs within colorectal crypts presenting ISD

### SI: HERV Transactivation by Viral Structural Proteins

The viral protein E1B 55K from Adenovirus type 5 (Ad5), along with the polyomavirus proteins JCV LTAg and Merkel cell polyomavirus (MCPyV) LTAg, were cloned into the pEGFP-N1 vector, whose expression is driven by a CMV promoter. All proteins were GFP-tagged to enable cellular visualization, and transcript production was confirmed via quantitative PCR (qPCR). For in vitro mRNA synthesis, the plasmids were linearized using specific restriction enzymes and transcribed using the HiScribe T7 ARCA mRNA Kit (NEB). Polyadenylated mRNA was subsequently purified using the Monarch RNA Clean-up Kit (NEB), following the manufacturer's protocol. Transfection of either plasmids or mRNA into colorectal carcinoma cell lines HCT8, HT29, and Col 33 was performed by electroporation using the 4D-Nucleofector™ system (Lonza).

Results

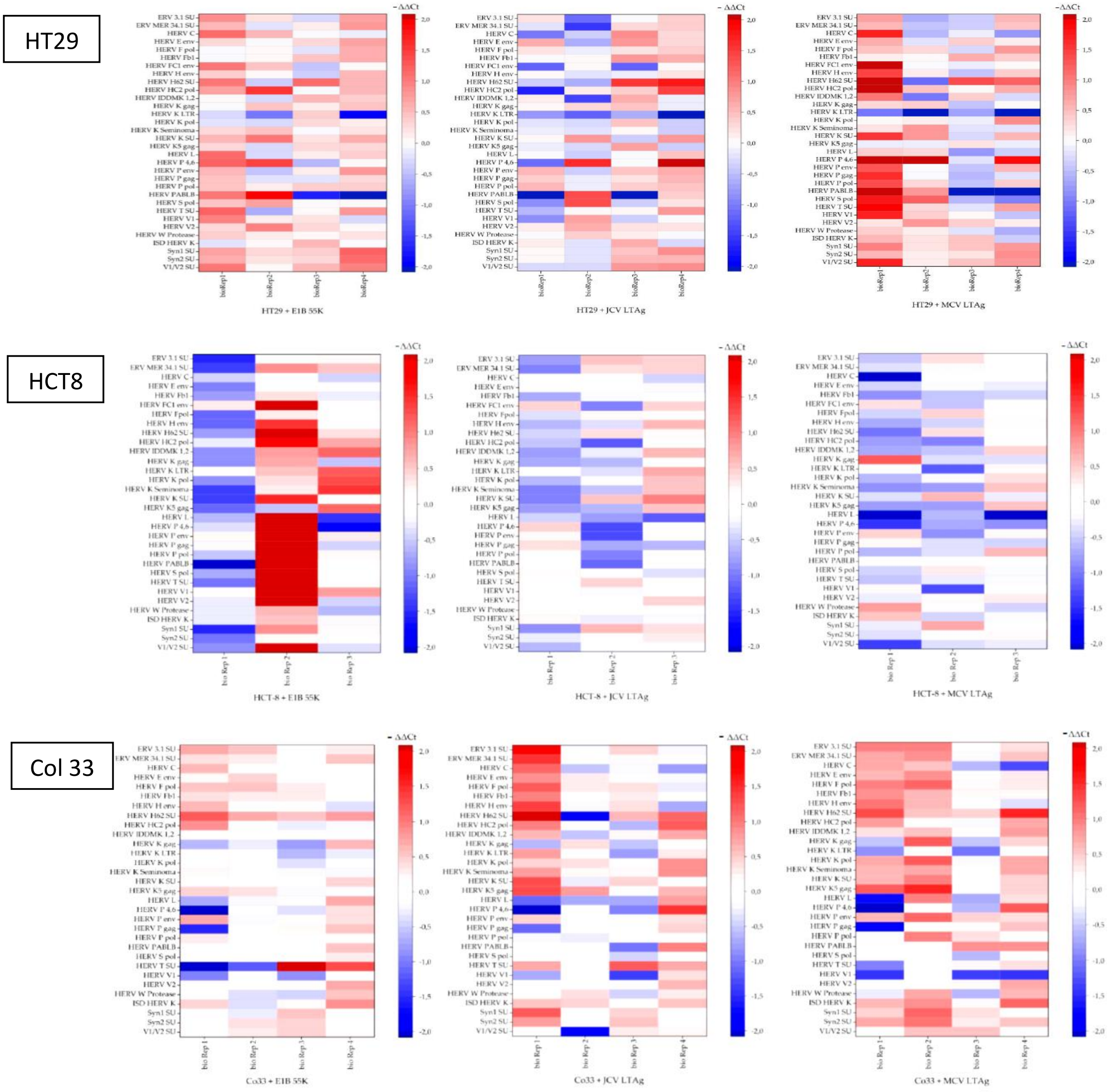

**Transcriptional Remodeling of HERVs: results across three CRC Cell Lines**

The experiments were designed to detect the added effect of viral proteins on HERV expression, measuring transactivation (upregulation) against the already high basal HERV transcript levels present in the CRC cell lines (HT29, HCT−8, and Co33). The results demonstrate distinct viral strategies for HERV manipulation.

**1. Adenovirus E1B 55K: The Most Potent General Transactivator**

E1B 55K consistently acted as the strongest general transactivator, amplifying HERV expression beyond basal levels:

- HCT-8 Maximal Effect: In HCT-8 cells, the upregulation was maximal, with many HERV loci showing ΔΔCt values exceeding +2.0, essentially saturating the detection scale across multiple families (e.g., HERV L and HERV P families).
- Broad Transactivation: In HT29, E1B 55K induced broad upregulation of loci like HERV L, Syn1 SU, and HERV V1/V2SU. In Col 33, it specifically transactivated $HERV_{3.1}$ SU, HERV C, and HERV E env.

**2. Polyomavirus LTAgs: Selective and Context-Dependent Transactivators**

The LTAgs primarily induced repression but exhibited highly selective and cell-specific transactivation:

- JCV LTAg Selectivity: In HT29 cells, JCV LTAg caused highly selective transactivation of the envelope subunits HERV H62 SU (ΔΔCt average ≈+0.53) and HERV HC2 pol. This specific activation pattern was lost in HCT-8 where these elements were generally repressed.
- MCV LTAg Plasticity: MCV LTAg demonstrated the greatest shift in function: it was an inconsistent regulator in HT29 but acted as a strong general transactivator in the primary Col 33 cells, upregulating most HERV families, including HERV H and HERV K.

**3. Consistently Transactivated Loci**

The HERV H62 SU element (an envelope surface subunit) was consistently transactivated by at least one LTAg across all three cell lines, suggesting a conserved viral mechanism targeting this specific envelope gene. Additionally, HERV K Seminoma was frequently upregulated by all three viral proteins across the cell lines.

**Biological consequences of HERV transactivation in cancer**

The viral-induced HERV transactivation (added upregulation over basal levels) has critical consequences, as it exacerbates processes already linked to CRC malignancy.

**1. Amplification of Oncogenic Signaling and Tumor Aggression**

The strong, added HERV transactivation, particularly by E1B 55K, further drives the malignant phenotype. Upregulation of elements like HERV K Seminoma amplifies the expression of endogenous retroviral proteins known to promote cell proliferation and inhibit apoptosis, thus increasing tumor aggressiveness.

**2. ISD Mediation of Immune Escape**

The transactivation of HERV envelope (env) genes is highly relevant for immune evasion. Specifically, the immunosuppressive domain (ISD) found in the envelope surface subunit (e.g., the consistently transactivated HERV H62 SU) can coat the CRC cell surface. This activity facilitates cancer immune escape by suppressing local immune responses, such as by promoting T-cell anergy or affecting regulatory T cells.

**3. Increased Cancer Heterogeneity via Fusogenic Capacity**

The transactivation of HERV env genes, such as HERV E env, can lead to the expression of envelope proteins with fusogenic capacity. This increased fusogenic potential can result in cell-cell fusion events within the tumor. Such fusion promotes cancer heterogeneity, genetic instability, and the formation of polyploid or aneuploid cells, which are strongly associated with increased invasiveness, metastasis, and therapy resistance.

**4. Viral Optimization and Host Context**

The selective transactivation seen with JCV LTAg (e.g., HERV H62 SU in HT29) suggests the virus precisely targets specific HERV products that aid transformation or immune modulation, while its concurrent repression conserves resources. The overall differential responsiveness confirms that the magnitude and specificity of HERV re-expression (transactivation) are shaped by the availability of host chromatin factors unique to each CRC cell line.

### SI: Induction of MHC-II molecules in CRCs by INF-γ

HCT8^WT

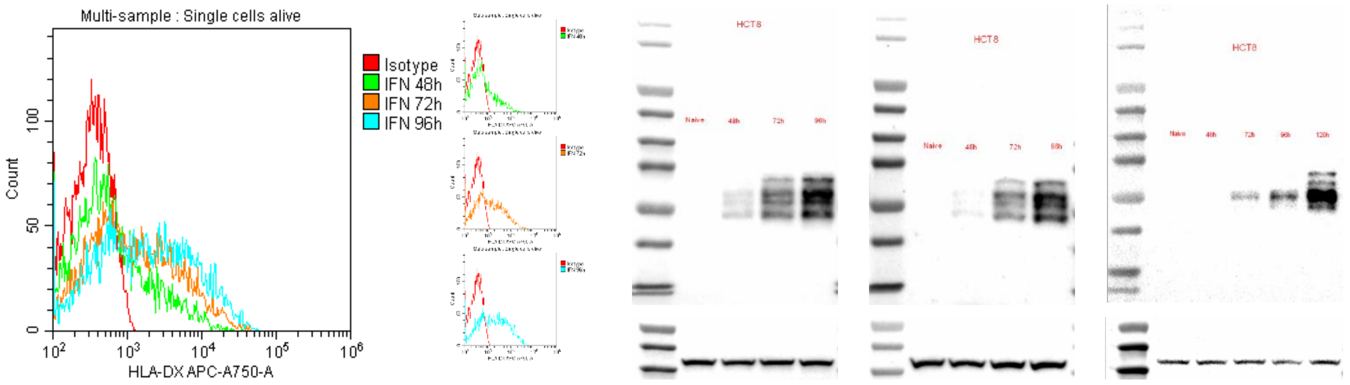


HT29^WT

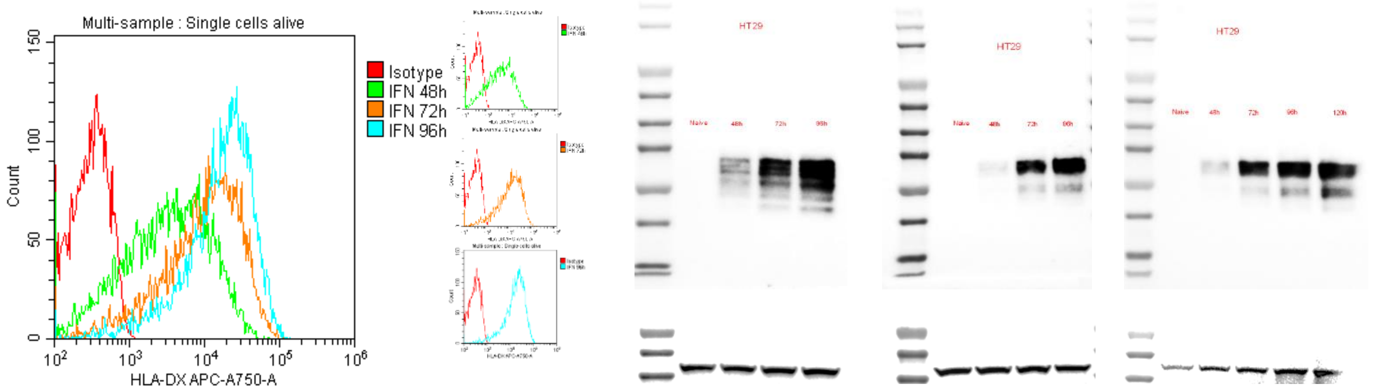


Col33^WT

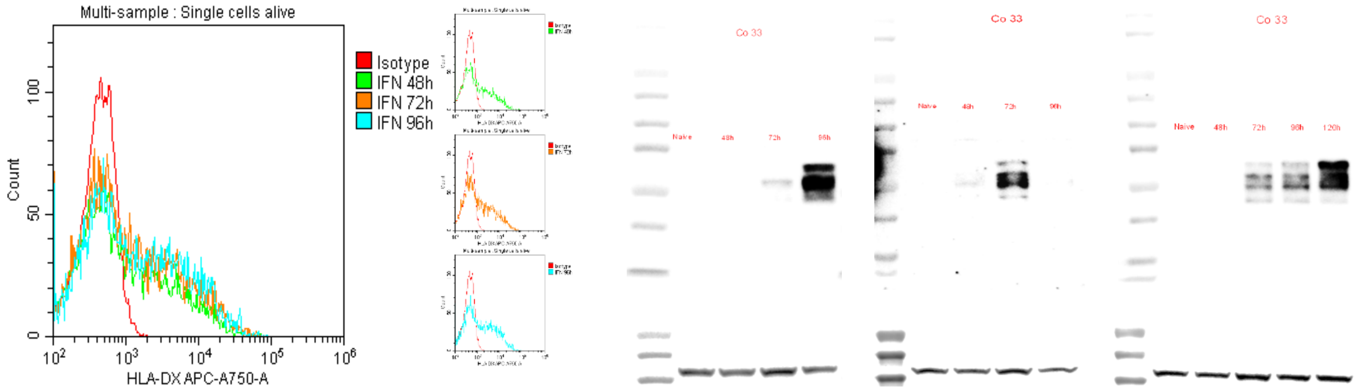

**SI: 3D Reconstruction from TEM**

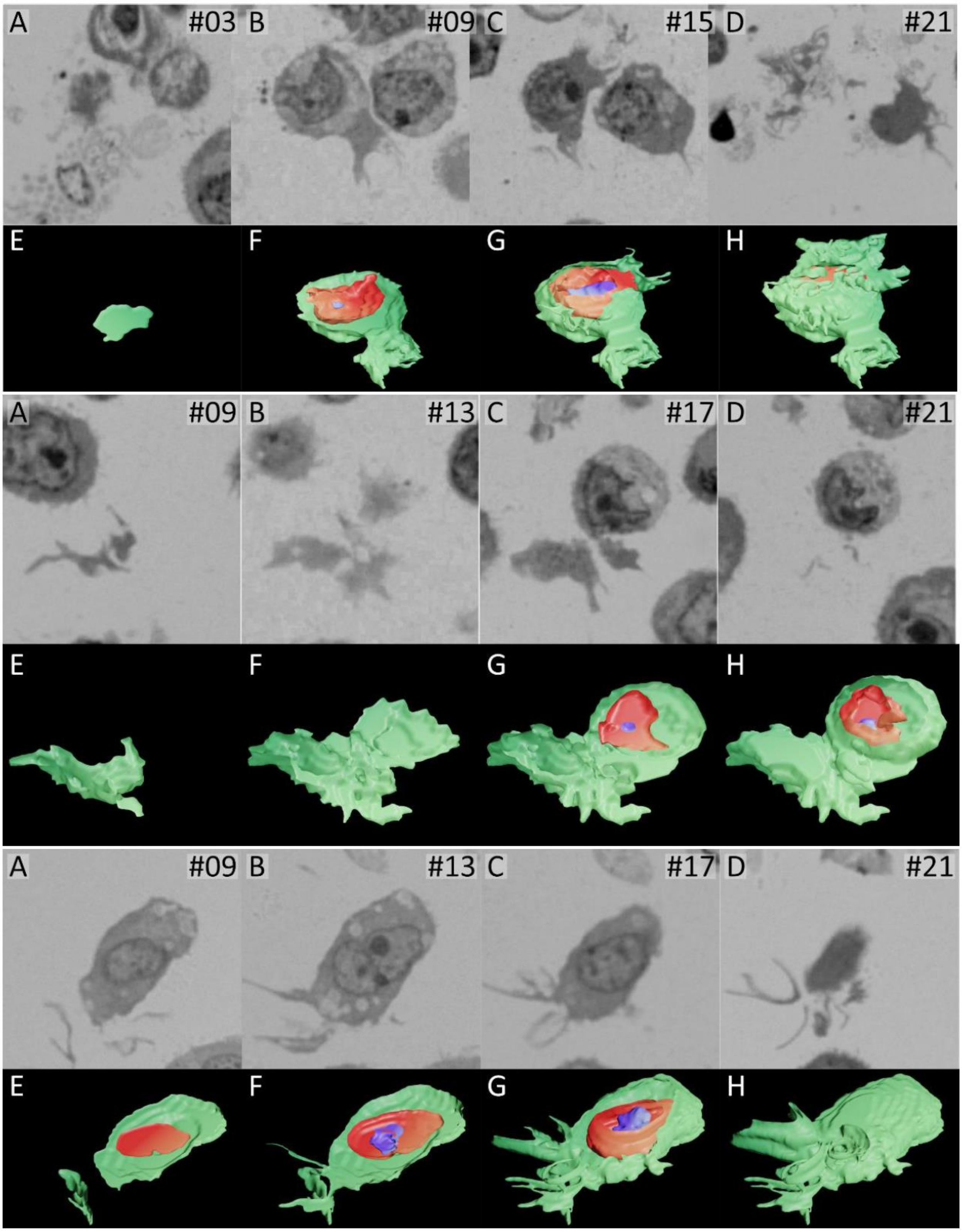

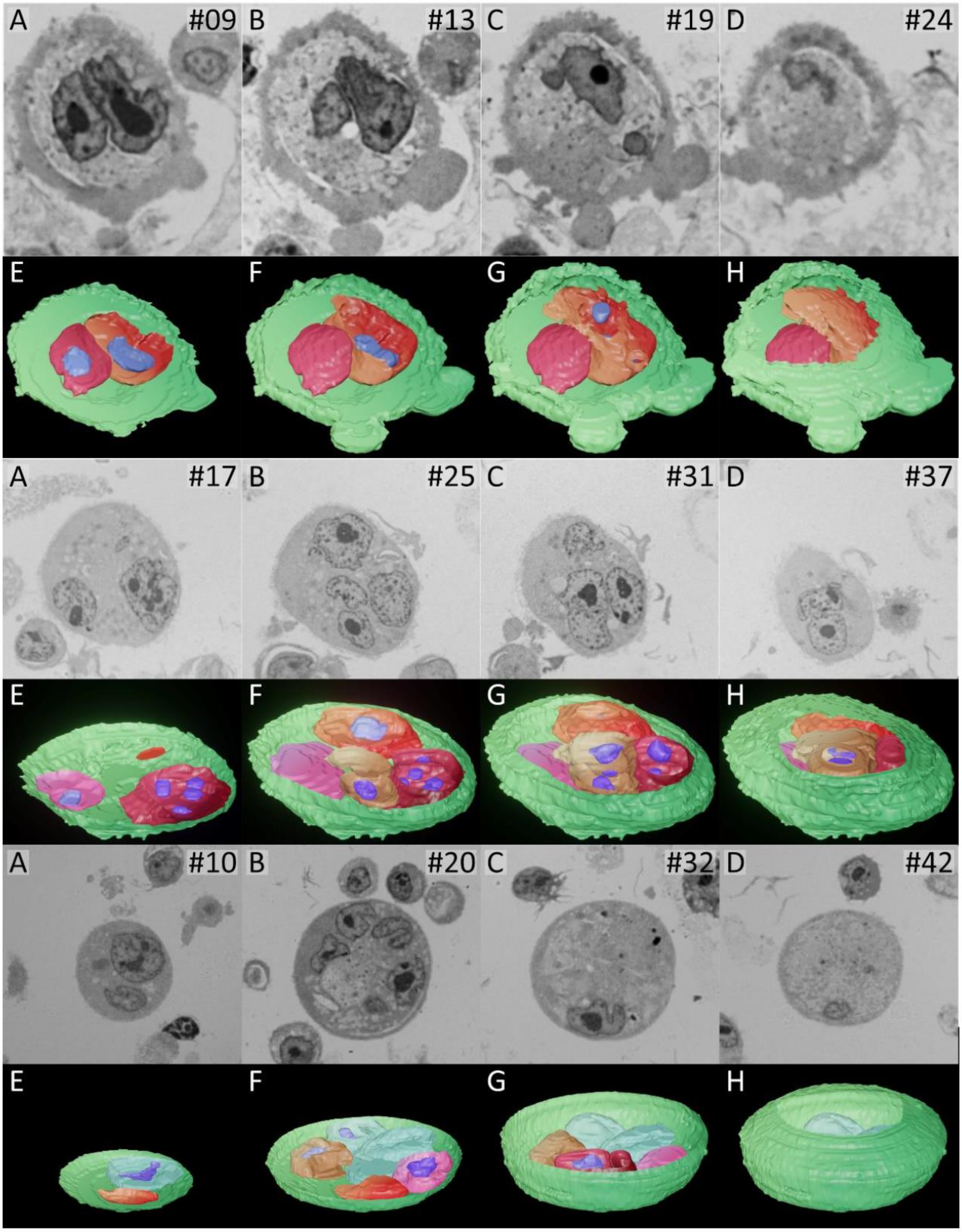
A #09
B #13
C #19
D #24
E
F
G
H
A #17
B #25
C #31
D #37
E
F
G
H
A #10
B #20
C #32
D #42
E
F
G
H

### SI: Commentaries on Videography

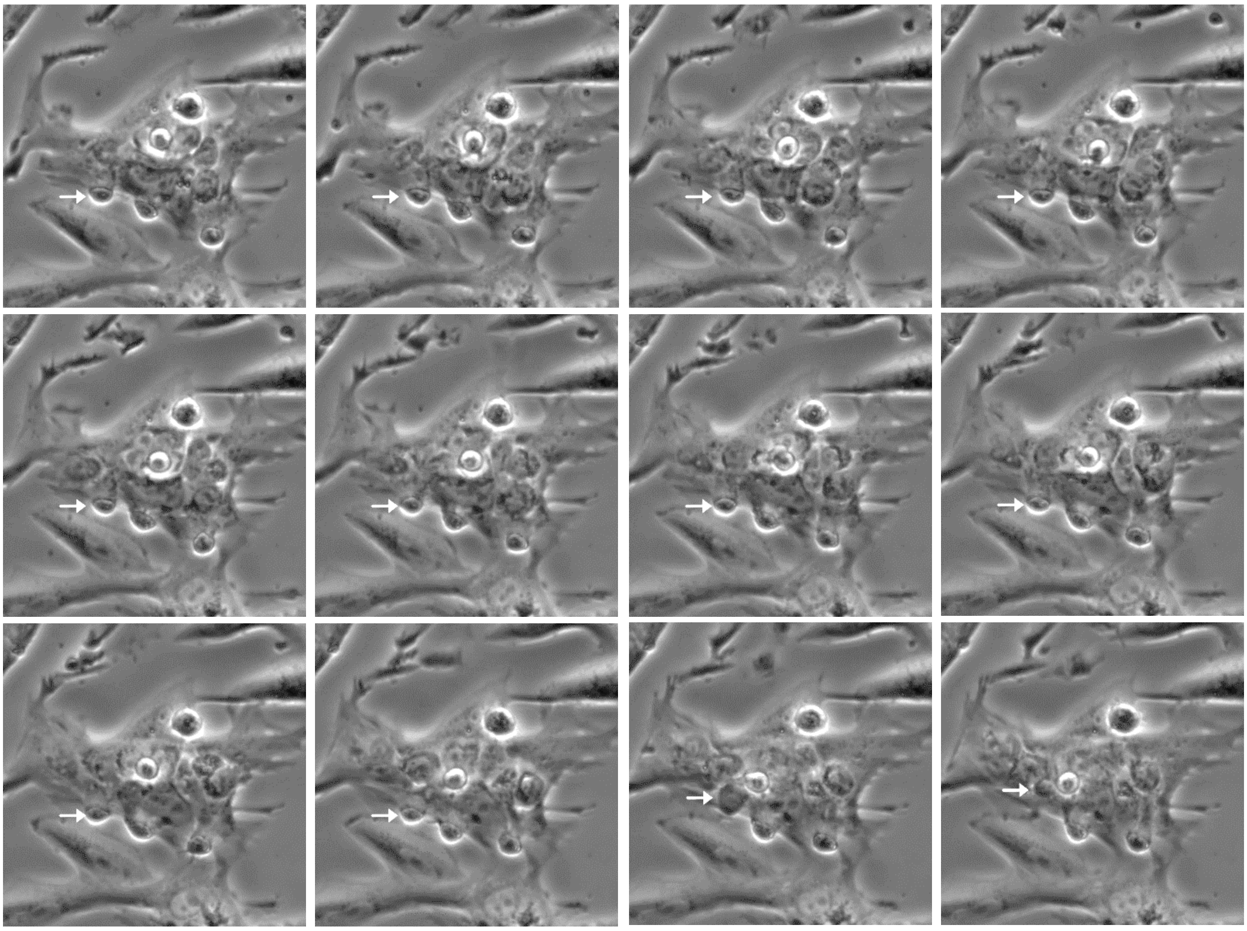

<u>Figure Video 9</u>. **Time-lapse sequence showing penetration of BLEICS cells into a cancer-associated fibroblast** (**CAF**), **corresponding to intracellular residency confirmed by TEM**. Sixteen sequential frames extracted from live-cell videography document the progressive entry of BLEICS cells into the cytoplasmic domain of a cancer-associated fibroblast (CAF). The CAF is identified by its prominent nucleus and adherent morphology, which remains structurally stable throughout the sequence. BLEICS cells, indicated by white arrows, approach the CAF from the periphery and gradually penetrate its cytoplasm without inducing visible membrane rupture, lysis, or apoptotic collapse. The spatial coherence and directional movement of the BLEICS cells across frames support a biologically regulated process rather than passive overlay or imaging artifact. The rounded morphology of both CAF and BLEICS cells reflects the fixation state following enzymatic detachment from culture substrates and should not be interpreted as representative of live-cell shape. Throughout the sequence, the CAF maintains its structural integrity, and no signs of phagocytic engulfment or vesicular enclosure are observed. This dynamic behavior corresponds directly to ultrastructural features documented in transmission electron microscopy (TEM) and 3D reconstructions, where BLEICS cells are visualized fully enclosed within CAF cytoplasm, exhibiting distinct nuclear and cytoplasmic boundaries. Together, these imaging modalities provide converging evidence for a previously undocumented cellular interaction: the active penetration and intracellular residency of BLEICS cells within non-disrupted CAFs. The underlying mechanisms remain to be elucidated.

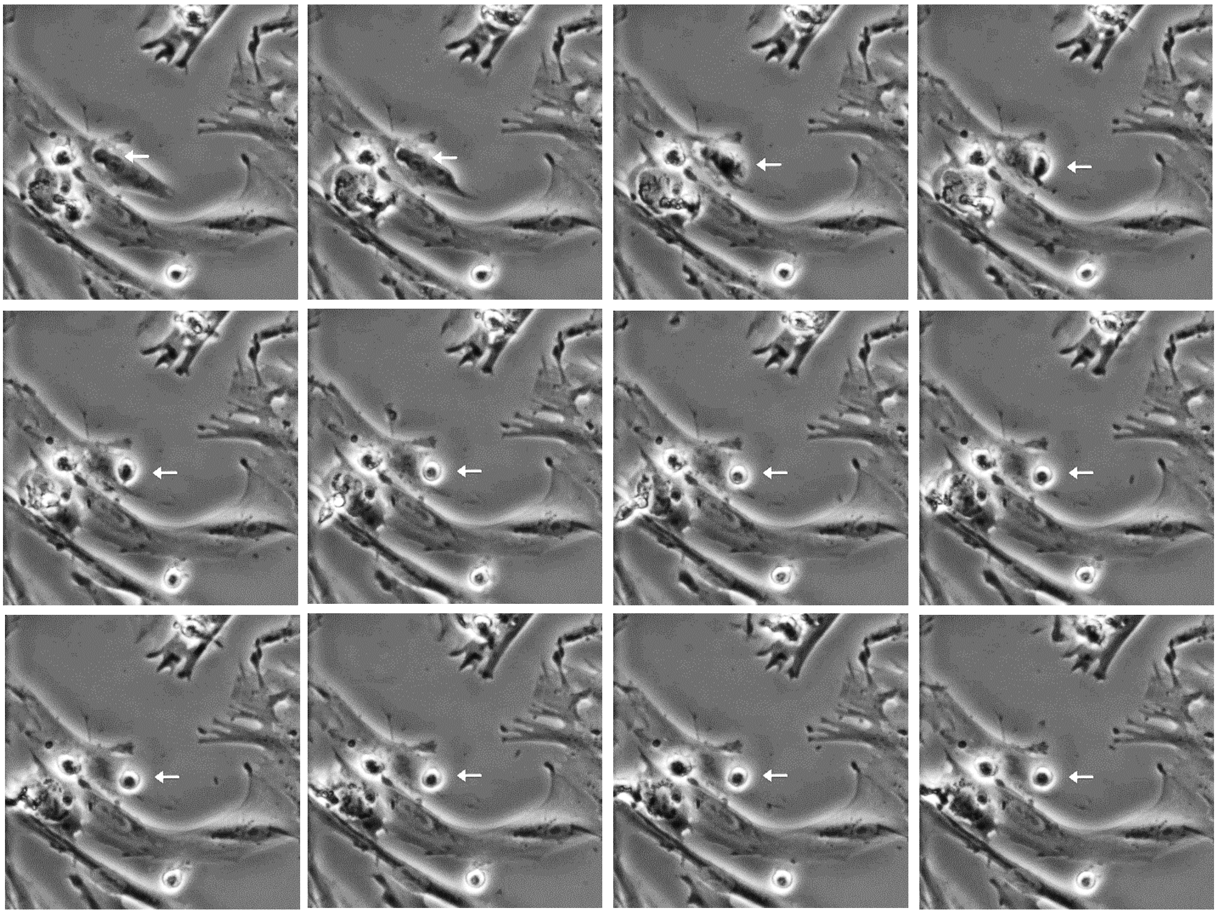

<u>Figure Video 10</u>: **Transcellular migration of a BLEICS cell through a cancer-associated fibroblast (CAF)**. Time-lapse microscopy sequence illustrating the transcellular egress of a BLEICS cell from within a CAF. The fifteen grayscale frames, extracted from live videography 10, depict a directional migration event in which the BLEICS cell transitions from an intracellular position to the extracellular space. White arrows highlight the progressive displacement of the BLEICS cell toward the CAF periphery, culminating in its exit without apparent disruption of the host cell membrane. The CAF maintains structural integrity throughout the sequence, suggesting a non-lytic, pore-mediated transcellular passage. This behavior may reflect a novel migratory phenotype of BLEICS cells.

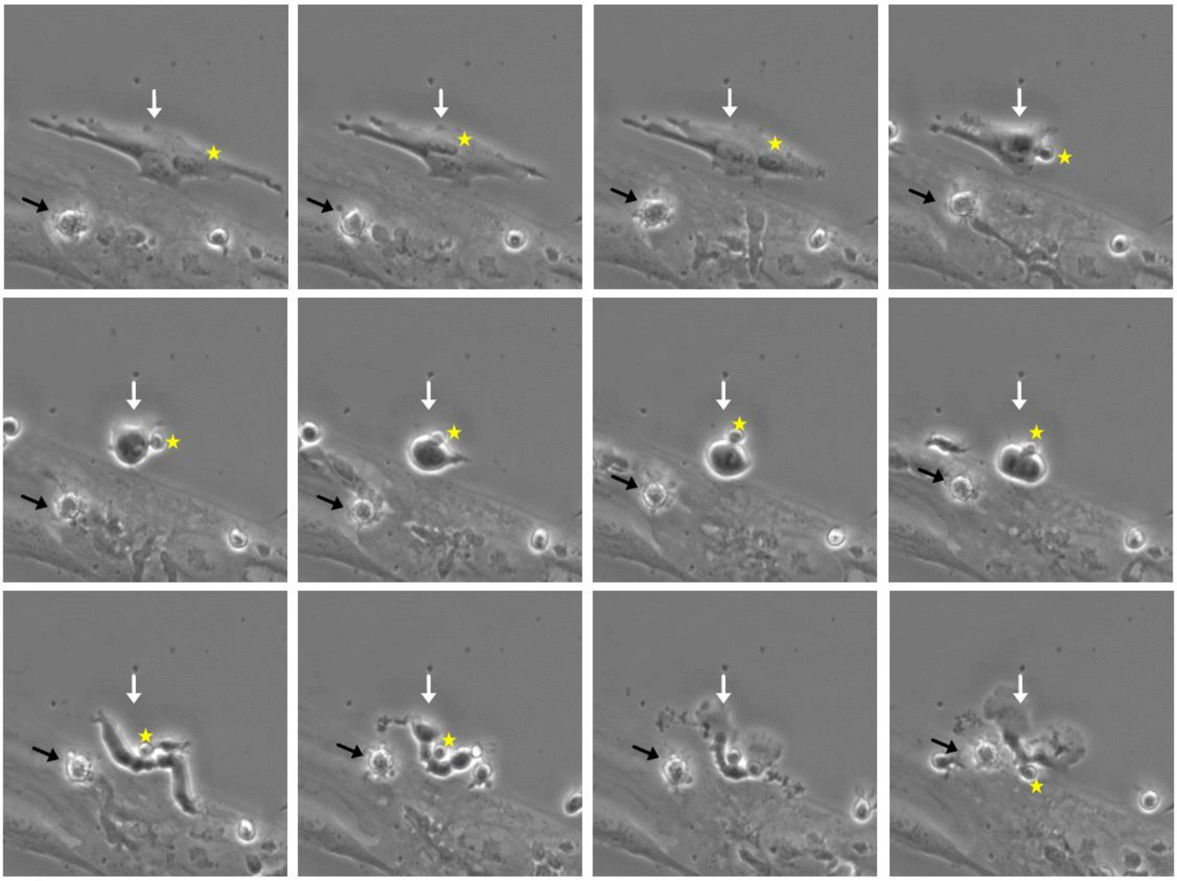

**<u>Figure Video 11</u>: BLEICS cell extrusion coinciding with CAF division: a novel dynamic interaction.** Static frames extracted from a time-lapse video sequence depict a previously uncharacterized cellular phenomenon involving BLEICS cells (yellow asterisks) and cancer-associated fibroblasts (CAFs). In this series, a BLEICS cell residing within a CAF initiates extrusion, as indicated by its progressive displacement toward the cell periphery. This transcellular exit appears temporally linked to the onset of CAF division, suggesting a possible mechanistic coupling between BLEICS cell egress and host mitotic activation. External BLEICS cells (black arrows) situated above the CAF provide spatial reference, reinforcing the interpretation of intracellular residency and clarifying the inside-outside orientation. White arrows mark the trajectory of the migrating BLEICS cell. The CAF maintains membrane integrity throughout, supporting a non-lytic, regulated passage. This sequence may represent a novel regulatory interaction between synthetic BLEICS constructs and host fibroblasts, with implications for engineered cell behavior and intercellular signaling.

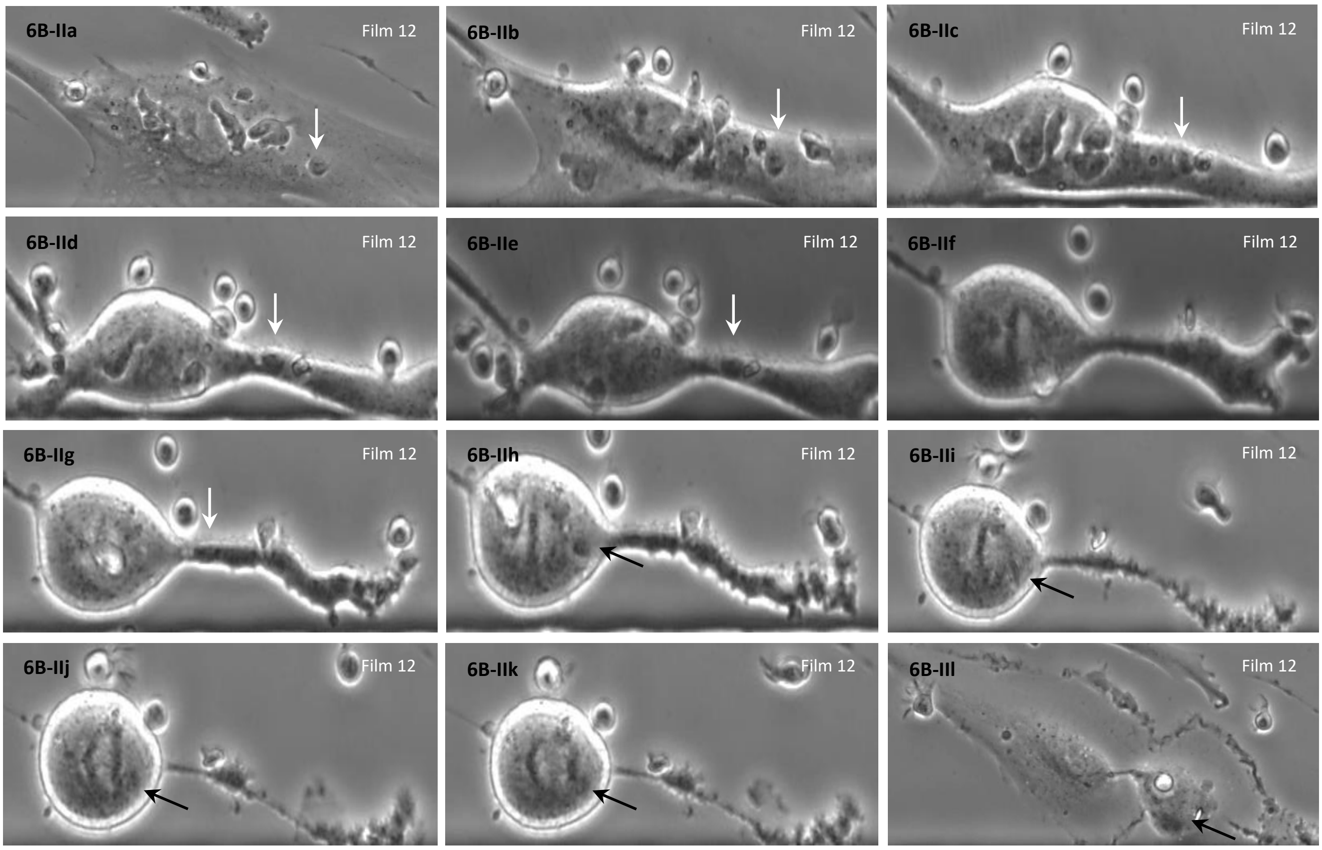


**<u>Figure Video 12</u>: Static sequence from time-lapse co-culture of BLEICS cells with CAFs.** Grayscale phase-contrast micrographs depict a 3×4 matrix of temporally ordered frames capturing the dynamic interaction between BLEICS cells and CAFs over the course of co-culture. BLEICS cells exhibit elongated morphology and progressive alignment along a horizontal substrate boundary, suggestive of directional migration and contact-mediated signaling. CAFs appear to modulate the local microenvironment, inducing morphological polarization and protrusive activity in adjacent BLEICS cells. The sequence highlights stromal-induced migratory behavior and potential paracrine or juxtacrine communication, consistent with early stages of tumor-stroma crosstalk.

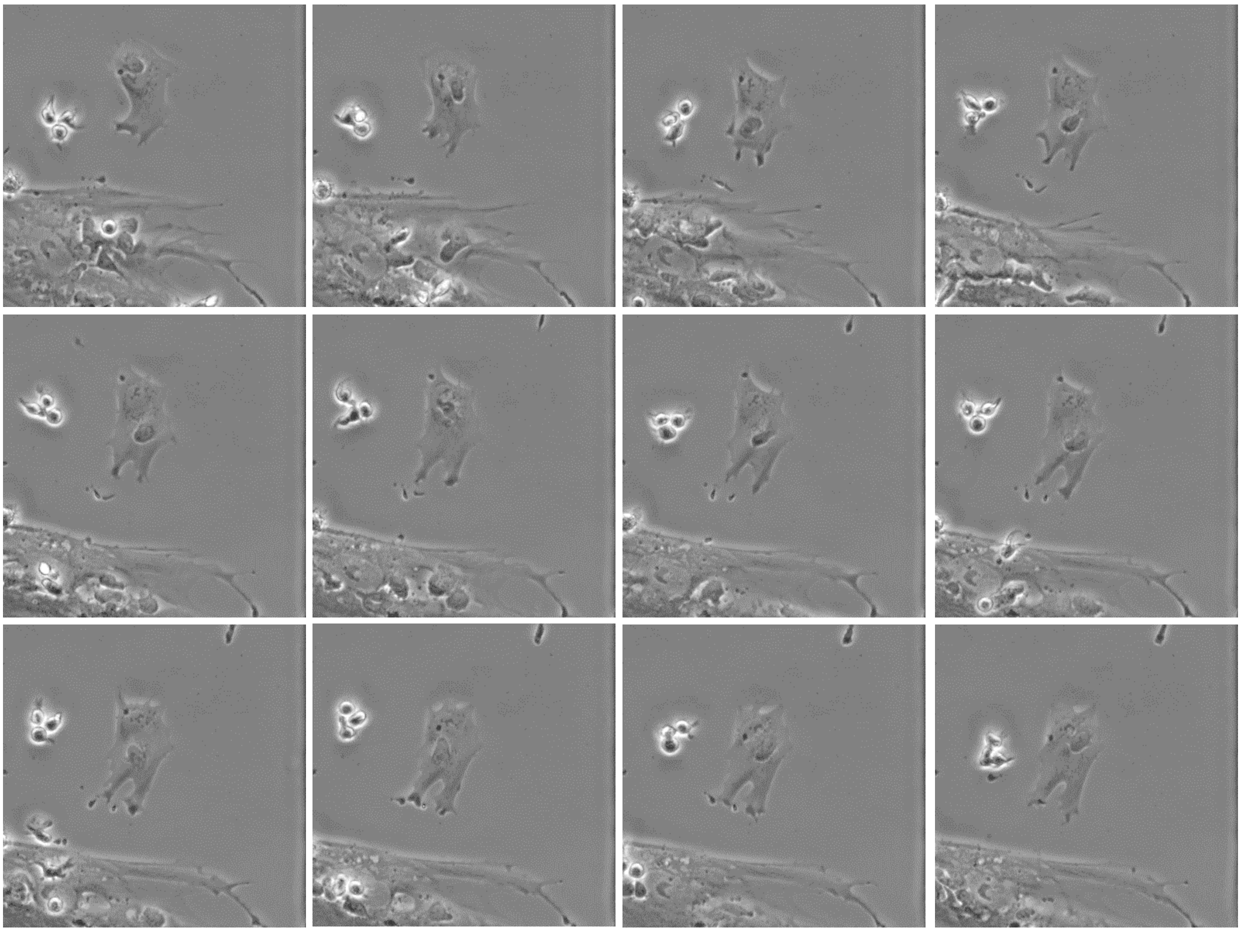

**<u>Figure Video 13 A</u>: BLEICS cell-induced nuclear displacement in co-cultured CAFs**. Sequential phase-contrast micrographs reveal a distinct biomechanical interaction wherein BLEICS cells exert upward-directed pressure on the nuclei of cancer-associated fibroblasts. The nuclei of CAFs, initially positioned centrally within the cytoplasm, appear progressively displaced toward the apical surface upon contact with BLEICS cells. This spatial reorganization suggests localized cytoskeletal remodeling and mechanical force transmission.

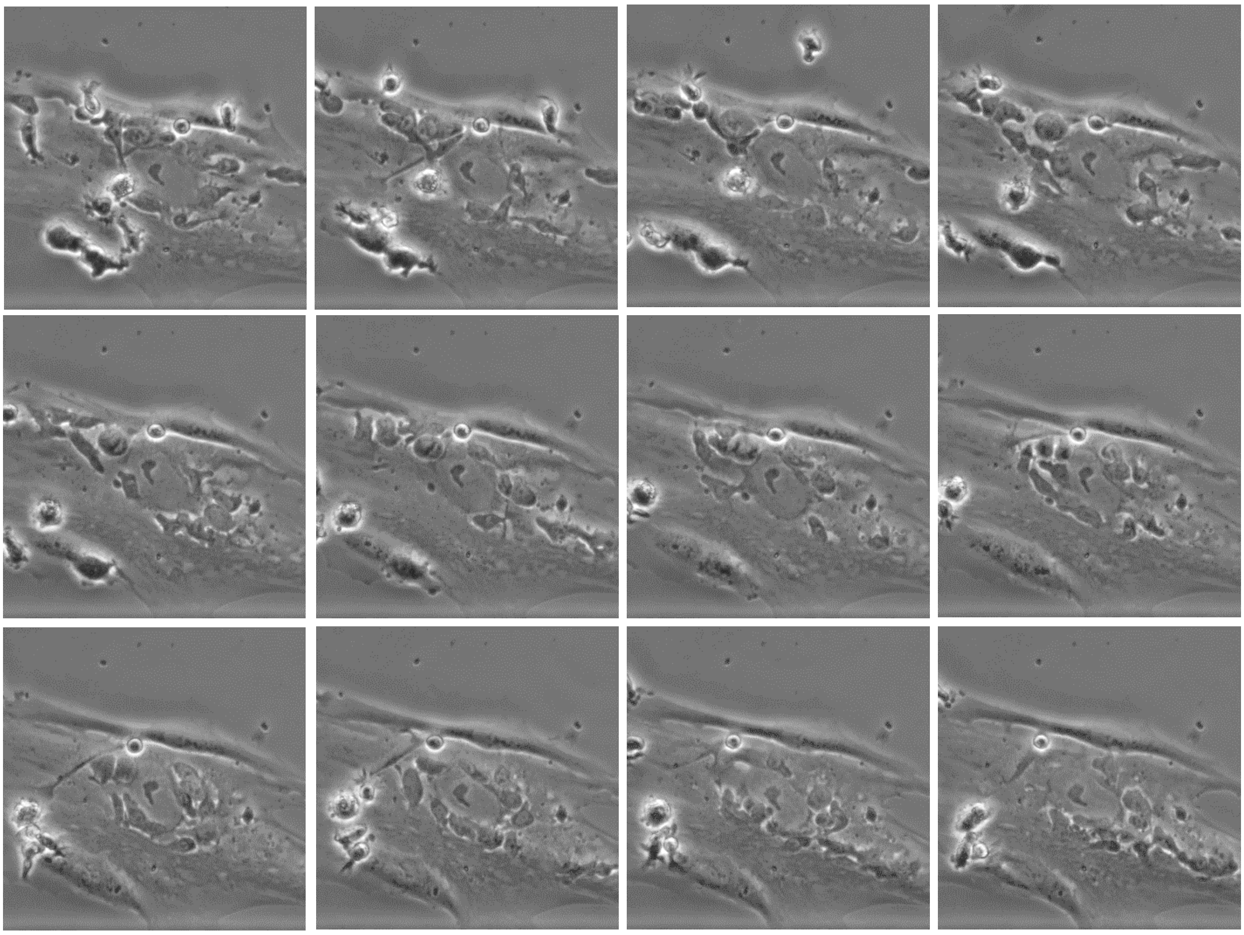

<u>**Figure Video 13 B**</u>**: BLEICS–CAF co-culture reveals dynamic intercellular interactions**. A 4×4 matrix of phase-contrast micrographs captures sequential frames from a co-culture assay involving BLEICS cells and cancer-associated fibroblasts (CAFs). BLEICS cells exhibit elongated, motile morphology and appear to engage directly with CAFs through both contact-dependent and spatially coordinated mechanisms. Notably, BLEICS cells extend protrusions toward CAFs, with several panels showing apparent physical displacement or compression of CAF nuclei, suggesting localized mechanical force transmission. CAFs respond with subtle changes in cytoplasmic architecture and nuclear positioning, indicative of reciprocal signaling or biomechanical adaptation. The consistent field of view across panels supports a time-resolved analysis of tumor-stroma crosstalk, highlighting the potential for BLEICS cells to modulate stromal behavior through direct interaction.

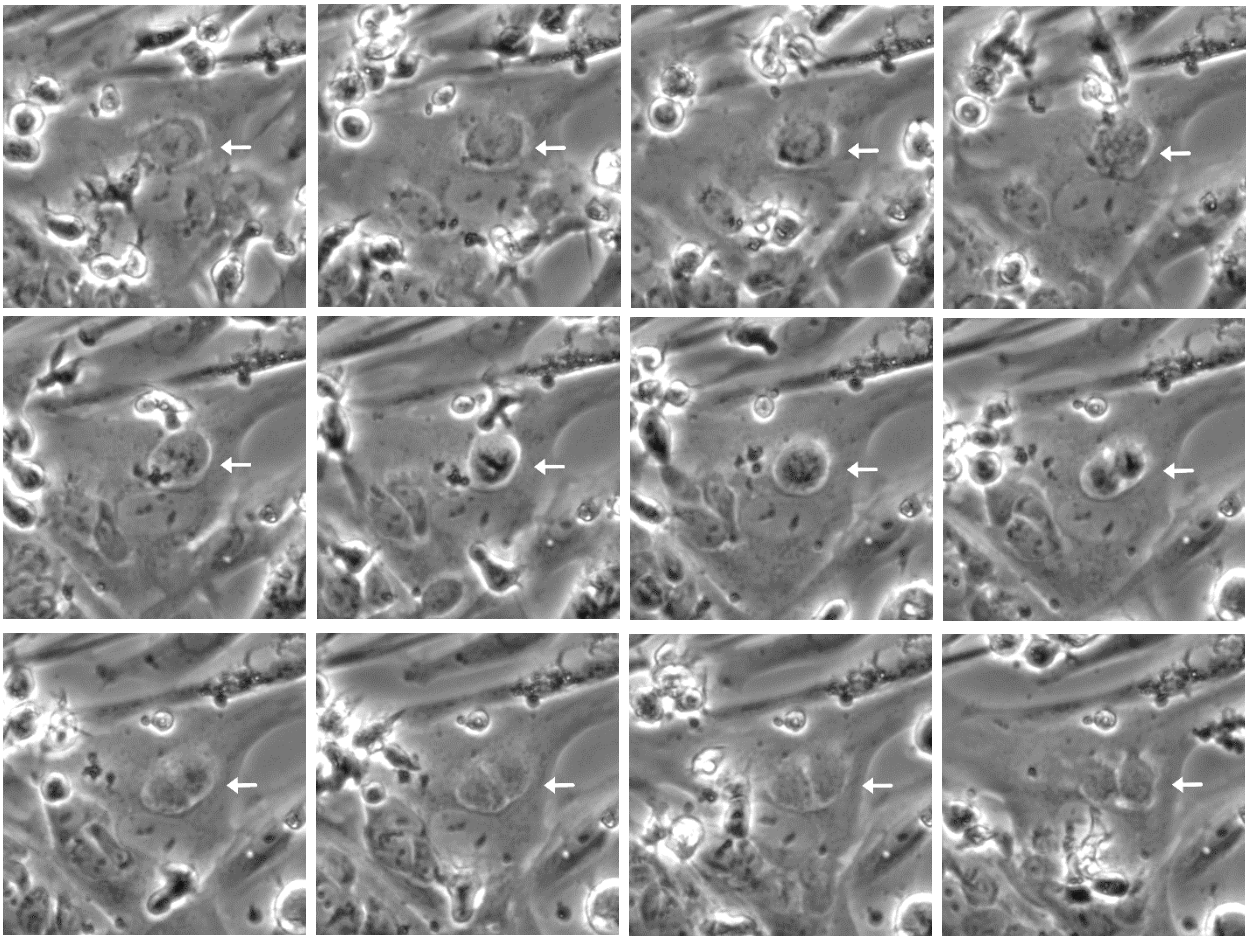

<u>Figure Video 15</u>: **Time-lapse microscopy sequence showing intracellular division of a BLEICS cell within a CAF**. Twelve sequential frames extracted from live-cell videography document a dynamic intracellular event involving a BLEICS cell residing within the cytoplasmic domain of a cancer-associated fibroblast (CAF). The CAF is identified by its prominent nucleus and adherent morphology, which remains structurally stable throughout the sequence. White arrows consistently mark the BLEICS cell positioned above the CAF nucleus. Over time, the BLEICS cell undergoes progressive morphological changes, including rounding, internal reorganization, and eventual segmentation into two adjacent compartments—consistent with mitotic division. The spatial coherence of the BLEICS cell and the absence of membrane rupture or extrusion support the interpretation of true intracellular residency during division. This behavior is corroborated by transmission electron microscopy (TEM) and 3D reconstruction data, which confirm the presence of BLEICS cells fully enclosed within CAF cytoplasm and exhibiting dual nuclear volumes. Together, these observations suggest a previously undocumented cellular phenomenon: the division of a BLEICS cell within a living CAF.

**SI: Cellularization Processes in BLEICS-Experienced CAFs**

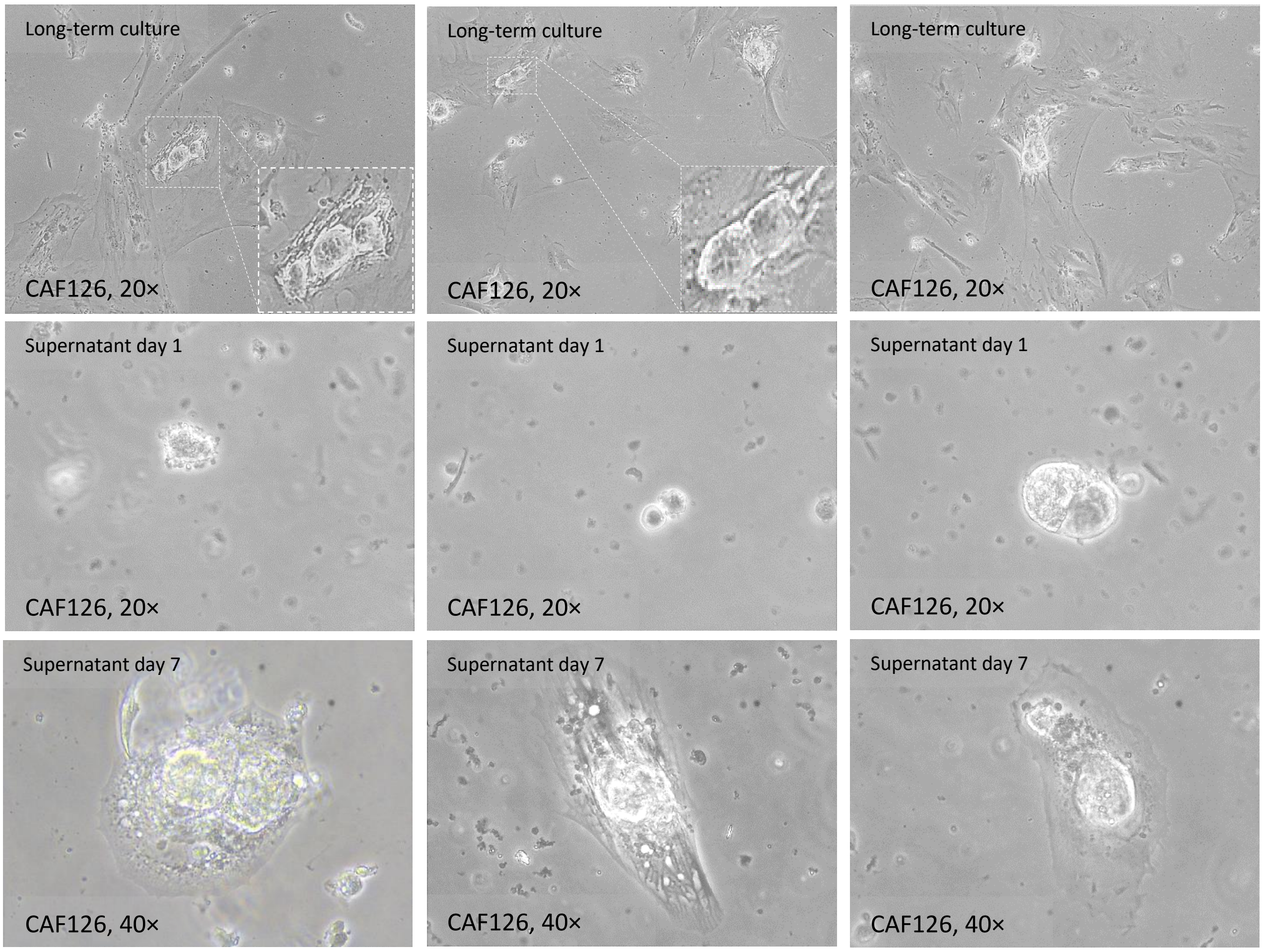


**Cellularization process in CAFs**. Tumor-derived fibroblasts undergo a cellularization process *in vitro*. Cellularization is a phenomenon observed in several tumor contexts. It begins with the separation of the nucleus from the surrounding cytoplasm and its subsequent release. The liberated nuclear bodies attach to the culture surface and initiate a new proliferative cycle. Not all nuclei retain the capacity to generate new colonies. The entire process unfolds over several days to weeks. Similar multinucleation-linked regenerative processes have been described previously (*A Distinct Oncogenerative Multinucleated Cancer Cell Serves as a Source of Stemness and Tumor Heterogeneity*. Cancer Res. 2018;78:2318–2331. doi: 10.1158/0008-5472.CAN-17-1861).

### SI: Cytogenetic I

**Panel A**: BLEICS revealed an abnormal karyotype

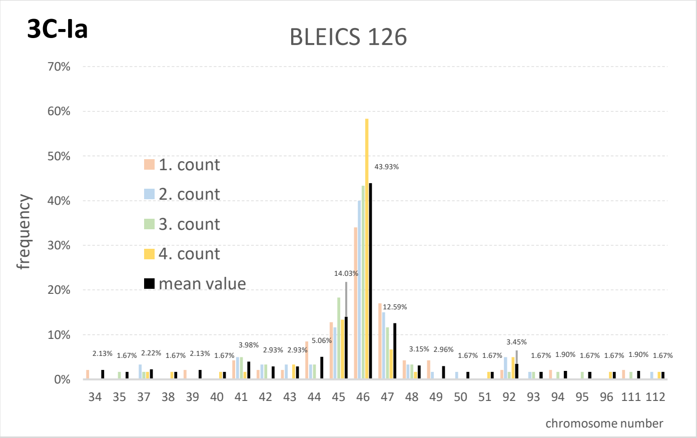


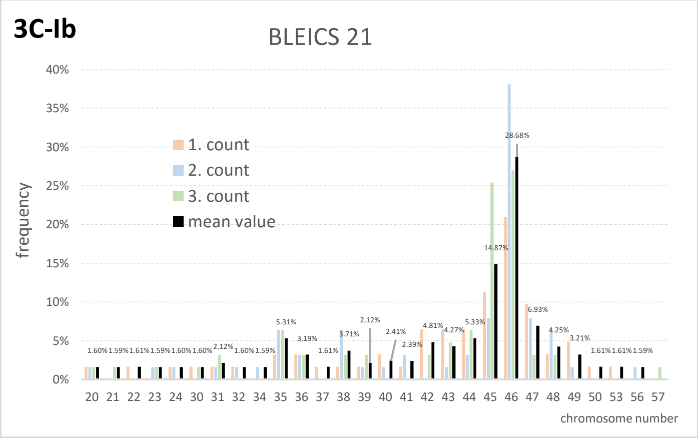


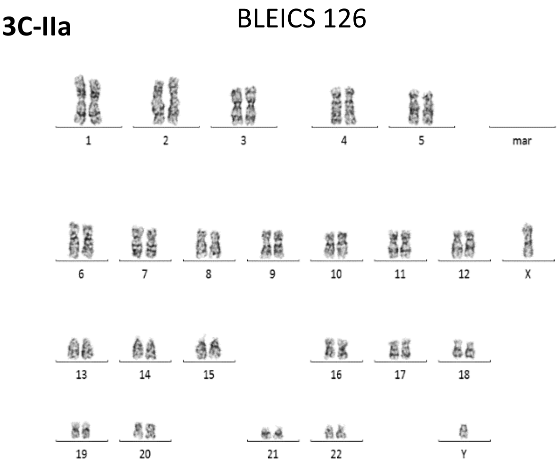


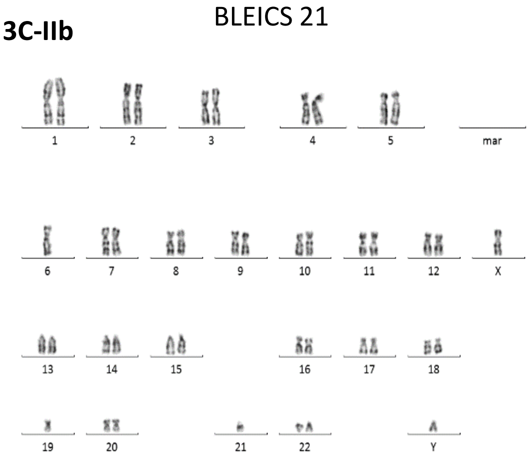


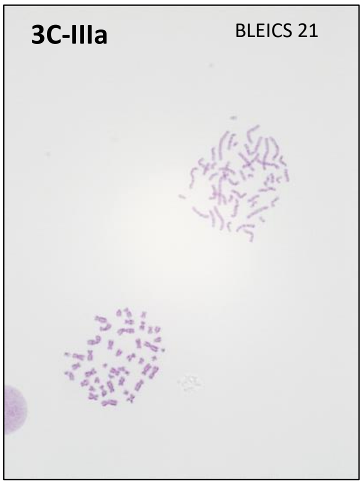


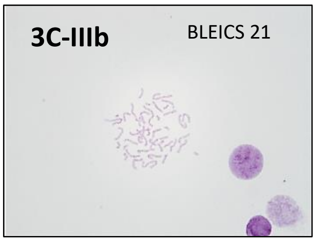


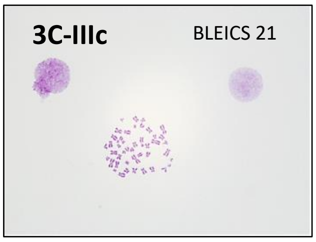


The inter-chromosomal size differences is result from the mix populations of fibroblasts and BLEICS, whereas calamari cells, with approximately 5µm of cell body and compact nuclei, are much smaller than the fibroblasts.

Cells were cultured in DMEM media and were incubated with 1 µg/ml Colcemid (Gibco, Germany) for 80 min, harvested from flask applying trypsin- EDTA (0.05/0.02 w/v)(Biochrome, Germany) and transferred to tubes. After centrifugation (170 g for 10 min), cells were incubated in hypotonic 0.56% KCl solution for 20 min and subsequently fixed and washed using a 3:1 methanol-glacial acetic acid fixative and dried at room temperature. For preparation of GTG-Bands, slides were incubated in 0.0025% Trypsin in PBS Dulbecco (Pan Biotech, Germany) for 60-90s and subsequently rinsed in PBS-Dulbecco followed by >10 min staining in Giemsa-solution (pH 6.88). Finally, slides were washed in bi-distilled water and cover slipped using Eukitt (Sigma-Aldrich, Germany). GTG bands were visualized on a Zeiss Axioscope 2 microscope. For detailed karyotyping in two patients, 100 metaphases were analyzed per sample using Ikaros software (Metasystems, Altlussheim, Germany). Karyotyping was performed at the Department of Human Genetics (Ruhr-University Bochum, Germany). For chromosome count only (performed in two patients) cells from 2 patients were stained using Giemsa solution, visualized using a Nikon microscope and counted by different operators at the Institute for Molecular Oncology, Marien-Hospital Herne, Germany.

### SI: Clonality and Ig Expression

<u>Clonal Studies of BLEICS</u>

Six separate PCRs with VH framework region 1 Primers for six VH subgroups, combined with a JH Primer mix.

| Cell line | VH amplicons | VH sequence (IGH) | Mutation frequency | productive? | comment |
|---|---|---|---|---|---|
| BLEICS 21 | VH3 | V3-11/NN/J6 | 3,0% | yes | monoclonal |
| BLEICS 105 | VH1 | V1-69/D3-10/J4 | 17,0% | yes | monoclonal |

For BLEICS 21 and 126, we obtained only one IGHV amplicon each and these were very clearly readable in Sanger direct sequencing. Somatically mutated, productive gene rearrangements were obtained. Therefore, monoclonal B cells are present in those lines. In very rare cases, B cells can also contain two productive VH rearrangements, but this violates allelic exclusion (each B cell produces only one antibody).

The occurrence of somatic mutations in the V gene allows us to conclude that the B cells are germinal center-experienced (where somatic hypermutation occurs), and are therefore germinal center B cells or memory B cells.

BLEICS 21 VH3

Productive VH rearrangement with IGHV3-11 and IGHJ6, somatically mutated with a frequency of 3,0 % in the IGHV segment. The DH gene was not clearly identifiable.

```
                                 __  CDR1 - IMGT  ______________ <--------------------------
                                                35                  40                  45
                                                     D   Y   Y   M   S   W   I   R   Q   A
BLEICS 21_VH3                   ... ... ... ... ..t gac tac tac atg agt tgg atc cgc cag gct
                                                S
X92287 Homsap IGHV3-11*03 F      ... ... ... ... ag- --- --- --- --- --c --- --- --- --- ---

                                  FR2 - IMGT  -------------------------> ______________  CDR2
                                                50                  55                  60
                                  P   G   K   G   L   E   W   V   S   Y   I   S   S   S
BLEICS 21_VH3                   cca ggg aag ggg ctg gag tgg gtt tca tat att agt agt agt ...

X92287 Homsap IGHV3-11*03 F      --- --- --- --- --- --- --- --- --- --c --- --- --- --- ...

                                  - IMGT  __________ <-------------------------------------
                                                65                  70                  75
                                      G   S   N   T   K   Y   A   D   S   V   K       G   R
BLEICS 21_VH3                   ... ggt agt aac aca aag tac gca gac tct gtg aag ... ggc cga
                                      S       Y       N
X92287 Homsap IGHV3-11*03 F      ... a-- --- t-- --- --c --- --- --- --- --- --- ... --- ---

                                 -------------------------------  FR3 - IMGT  ---------------
                                                80                  85                  90
                                  F   T   I   S   R   D   N   A   K   N   S   L   Y   L   Q
BLEICS 21_VH3                   ttc acc atc tcc aga gac aac gcc aag aac tca ctg tat ctg caa

X92287 Homsap IGHV3-11*03 F      --- --- --- --- --- --- --- --- --- --- --- --- --- --- ---

                                 -----------------------------------------------------> ____
                                                95                 100              104
                                  M   N   S   L   R   A   E   D   T   A   V   Y   Y   C   A
BLEICS 21_VH3                   atg aac agc ctg aga gcc gag gac acg gcc gta tat tac tgt gcg

X92287 Homsap IGHV3-11*03 F      --- --- --- --- --- --- --- --- --- --- --g --- --- --- ---

                                 ___________________  CDR3 -  IMGT _____________________

                                  R   D   Q   Y   S   G   Y   D   S   W   P   P   S   V   W
BLEICS 21_VH3                   aga gat caa tat agt ggc tac gat tcc tgg ccc cct agc gtc tgg

X92287 Homsap IGHV3-11*03 F      ---


                                  G   Q   G
BLEICS 21_VH3                   ggc caa ggg

X92287 Homsap IGHV3-11*03 F




BLEICS 21_VH3                   gatcaatat
J00235 Homsap IGHD2-21*01 F      ---tgc---tcc
X13972 Homsap IGHD3-10*01 F      t--t-c---ggttcggggagttattataac
X93615 Homsap IGHD3-10*02 F      t--t-c---gttcggggagttattataac
X97051 Homsap IGHD3-16*02 F      t---gt---acc
X93616 Homsap IGHD3-22*01 F      t--t-c---gatagtagtggttattactac


BLEICS 21_VH3          gatcaatatagtggctacgattcctggcccccctagcgtctggggccaaggg
X86355 Homsap IGHJ6*02 F  .........a-ta----t-c-a--acggtatgga----------------accacggtc
```

<u>BLEICS 126 VH1</u>

Productive VH rearrangement with IGHV1-69, IGHD3-10, and IGHJ4, somatically mutated with a frequency of 17,0 % in the IGHV segment.

```
                                   ------------------------------------------> ________________
                                                    20                  25                  30
                                                            S   C   K   S   S   G   I   N   F
BLEICS 105_VH1                     ... ... ... ... ... .tc tcc tgc aag tcc agt gga atc aac ttc
                                    G   S   S   V   K   V               A           G   T
Z14309 Homsap IGHV1-69*08 F         ggg tcc tcg gtg aag g-- --- --- --- g-t tc- --- gg- -c- ---

                                    __  CDR1 - IMGT  ______________ <--------------------------
                                                    35                  40                  45
                                                    F   N   Y   T   I   S   W   V   R   Q   A
BLEICS 105_VH1                     ... ... ... ... ttt aat tat acc atc agt tgg gtg cga cag gcc
                                                    S   S
Z14309 Homsap IGHV1-69*08 F         ... ... ... ... agc -gc --- --t --- --c --- --- --- --- ---

                                     FR2 - IMGT  ------------------------> ______________  CDR2
                                                    50                  55                  60
                                     P   G   Q   G   L   E   R   M   G   R   I   I   P   F
BLEICS 105_VH1                     cct gga cag ggg ctt gag cgg atg gga agg atc atc cct ttt ...
                                                            W                           I
Z14309 Homsap IGHV1-69*08 F         --- --- --a --- --- --- t-- --- --- --- --- --- --- a-c ...

                                     - IMGT  __________ <---------------------------------------
                                                    65                  70                  75
                                         L   G   T   P   K   Y   E   Q   K   Y   Q       G   R
BLEICS 105_VH1                     ... ctt ggc act cca aaa tac gaa cag aaa tat cag ... ggc aga
                                                    A   N       A           F
Z14309 Homsap IGHV1-69*08 F         ... --- --t --a g-- --c --- -c- --- --g -tc --- ... --- ---

                                    ------------------------------  FR3 - IMGT  ---------------
                                                    80                  85                  90
                                     V   T   I   T   A   D   K   S   T   G   T   V   Y   L   E
BLEICS 105_VH1                     gtc acg att acc gcg gac aag tca acg ggc aca gtt tac ttg gaa
                                                                        S       A       M
Z14309 Homsap IGHV1-69*08 F         --- --- --- --- --- --- --a --c --- a-- --- -cc --- a-- --g

                                    ------------------------------------------------------> ____
                                                    95                  100             104
                                     L   S   G   L   R   S   D   D   T   A   V   Y   Y   C   A
BLEICS 105_VH1                     ctt agt ggc ctg aga tct gat gac acg gcc gtc tat tat tgt gcg
                                             S               E
Z14309 Homsap IGHV1-69*08 F         --g --c a-- --- --- --- --g --- --- --- --g --- --c --- ---

                                    _______________________  CDR3 -  IMGT ______________________

                                     R   E   G   R   E   N   Y   Y   K   V   V   S   Y   F   D
BLEICS 105_VH1                     aga gag ggt cgg gag aat tac tat aag gtg gtt tcg tat ttt gac

Z14309 Homsap IGHV1-69*08 F         --- --


                                    ___

                                     S   W   G   P   G
BLEICS 105_VH1                     tcg tgg ggc ccg gga

Z14309 Homsap IGHV1-69*08 F


BLEICS 105_VH1                     gggtcgggagaattactataaggtggtt
X13972 Homsap IGHD3-10*01 F         t---tc--g--g---t-----c



BLEICS 105_VH1             gggtcgggagaattactataaggtggtttcgtattttgactcgtggggcccggga
X86355 Homsap IGHJ4*02 F     ............................ac--c-------ac-------a----accct
```

Secretion of Immunoglobulins by BLEICS 21 and BLEICS 126

BLEICS cells express immunoglobulins (monoclonal type) which are somehow abnormal in their molecular weights (are bigger than 150 KDa, see Pic below). Thus, we thought that those patients have some kind of B-cell lymphoma in the gut. In this regard, it is noteworthy to mention that there are scientific reports revealing that many cancer types of epithelial origin may produce immunoglobulins and express MHCII molecules. We certainly observed is a reliable MHCII expression on CRC cells after IFNγ exposure by CRCs

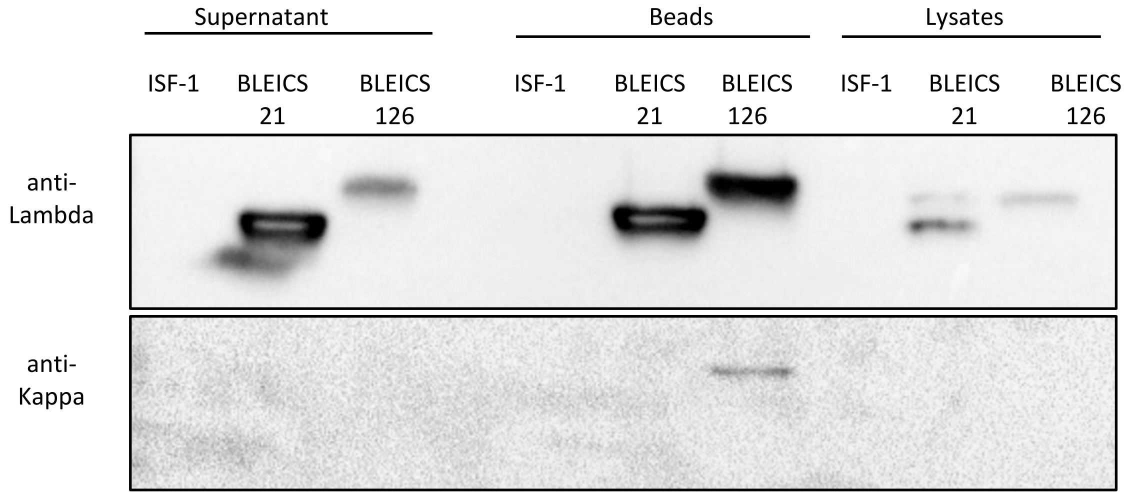


The experimental protocol involved incubating BLEICS 21 and BLEICS 126 cells overnight at 37°C in ISF-1 medium, followed by centrifugation of the cells at 100 × g for 5 min and collection of the supernatant. Magnetic beads were then used for Igs capture, wherein 100 µl of beads were initially blocked in 2 ml of pure ISF-1 medium at room temperature for 30 min. The beads were subsequently resuspended in 100 µl of BLEICS 21 and BLEICS 126 supernatant and incubated overnight at 4°C, followed by removal of the medium using a magnet. The beads were then dissolved in 25 % Laemmli sample buffer and 10 % mercaptoethanol and incubated at 95°C for 10 min. In the case of lysate samples, cell pellets were dissolved in RIPA buffer and subjected to ultrasound lysis followed by a centrifugation at 14 000 × g for 20 minutes. Supernatants were then resolved on PAGE. Gels were stained with Coomassie blue and distained in MeOH:Acetic acid 40:10 v/v. Gels were documented in a Biorad Imager Chemidoc$^{TM}$ XRS$^{+}$.

The above Figure illustrates the expression of lambda chains in both BLEICS 21 and BLEICS 126. The expression of lambda chains appears to be more pronounced in BLEICS 21 supernatant compared to BLEICS 126. After concentration with magnetic beads, the intensity of lambda chains is very similar for both cell lines. In contrast, the concentration of lambda chains obtained from the cell lysate is relatively low. In the case of kappa chains, only a low intensity was detected for BLEICS 126 even after an extended exposure time during the measurement following purification with magnetic beads. For all other samples, no signal was measurable.

According to the expected molecular weight of both chains (22.5 kDa), the lambda chains yielded two distinct bands, one around 25 kDa and the other slightly lower. However, only one band of approximately 25 kDa was detected for the Kappa chain, which was observed solely in the BLEICS 126 supernatant.

| HERV | -LOG10 (P VALUE) | ΔTPM | SUBGENES | ISD | BESTREFRV | SUPER GROUP |
|---|---|---|---|---|---|---|
| 1227 | 1.5 | 5.7 | 5LTR MA CA NC Prot RT IN TM PPT 3LTR | **LdNqiALDhLLA** | | HEPSI |
| 2636 | 2.3 | 5.0 | 5LTR PBS MA CA NC Prot RT IN TM 3LTR | **LQNRRGLDLLFiSQGGLCtALE** | Cat ECE1 endogenous retroviral element DNA, partial 57% -3795 | MLLV |
| 2858 | 1.7 | 5.7 | 5LTR PBS MA CA NC Prot RT IN TM 3LTR | **LQNRmALDiiTtAQGGTCtLtK** | | HERVFRDLIKE |
| 5106 | 1.8 | 6.2 | 5LTR PBS NC Prot RT IN SU TM PPT 3LTR | **YQNRLALDYLLA** | Homo sapiens retinoic acid-inducible endogenous re 85% -6483 | HERVERI |
| 5108 | 2.1 | 5.4 | 5LTR PBS CA DU IN SU TM PPT 3LTR | **YQNRLALDYLLA** | Human endogenous retroviral DNA (4-1), complete re 68% 1017 | HERVERI |
| 5115 | 2.2 | 8.1 | 5LTR PBS CA NC Prot RT IN SU TM PPT 3LTR | **LQNczGLYLLttEKaGLCtFLG** | Human endogenous retrovirus pHE.1 (ERV9). 90% -3615 | HERVW9 |
| 5122 | 1.5 | 6.9 | 5LTR PBS MA CA NC Prot IN SU TM PPT 3LTR | **LQNRRGLDcLTAEKGGLCtFLG** | Human endogenous retrovirus pHE.1 (ERV9). 74% -3430 | HERVW9 |
| 5208 | 1.4 | 7.6 | 5LTR PBS MA CA NC Prot RT IN SU TM PPT 3LTR | **YQNRLvLDzLLA** | Homo sapiens retinoic acid-inducible endogenous re 67% -6129 | HERVERI |
| 6128 | 1.7 | 4.2 | 5LTR PBS MA CA NC Prot RT IN SU TM 3LTR | **cQNRLALDYLLA** | Human endogenous retroviral DNA (4-1), complete re 80% 299 | HERVERI |
| 877 | ∞ | -7.6 | 5LTR PBS MA CA NC Prot RT IN SU TM PPT 3LTR | **YQNRLALyYLLA** | Human endogenous retroviral DNA (4-1), complete re 79% 261 | HERVERI |
| 906 | ∞ | -6.0 | 5LTR PBS MA CA NC Prot RT IN SU TM PPT 3LTR | **LQNRqsLDLLTAEKGGLCIFLN** | Homo sapiens RGH2 gene, retrovirus-like element. 83% -22 | HERVHF |
| 1303 | ∞ | -5.1 | 5LTR PBS MA CA NC DU Prot RT IN SU TM PPT 3LTR | **RQTVIWMGDRLMSLEyIFQLQC** | Human endogenous retrovirus HERV-K(II) DNA, complete 84% 981 | HML |
| 1716 | ∞ | -4.5 | 5LTR PBS CA NC DU Prot RT IN TM PPT 3LTR | **LRziVMWLGDRV** | Human endogenous retrovirus HERV-K10. 62% 1031 | HML |
| 1814 | ∞ | -4.0 | 5LTR MA CA IN SU TM PPT 3LTR | **YQNRLALDYLLA** | Human endogenous retroviral DNA (4-1), complete re 64% 581 | HERVERI |
| 1968 | ∞ | -4.2 | 5LTR PBS NC Prot IN SU TM PPT 3LTR | **LQNRzGLDLLTAEKGGLCIFLN** | Human endogenous retrovirus, complete genome. 92% 792 | HERVHF |
| 3698 | ∞ | -4.0 | IN SU TM PPT 3LTR | **LQNhRGLDLLTAEKGGLCIFLN** | Homo sapiens RGH2 gene, retrovirus-like element. 77% 4627 | HERVHF |
| 4352 | 1.4 | -4.8 | 5LTR MA CA NC Prot IN SU TM PPT 3LTR | **YzNRLALDYHLA** | Homo sapiens retinoic acid-inducible endogenous re 96% -4626 | HERVERI |
| 4593 | ∞ | -4.4 | 5LTR PBS MA CA NC Prot RT SU TM 3LTR | **LQNRqGLDLLTAEKGGfCtFLG** | Human endogenous retrovirus pHE.1 (ERV9). 89% -2775 | HERVW9 |

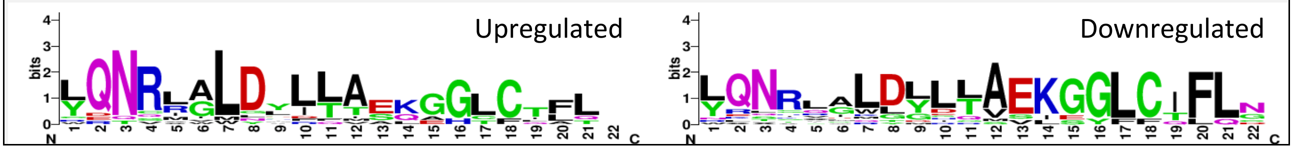


**Table 2:** Differentially expressed HERVs with ISD in BLEICS cells

### SI: CRC primary cells morphology and cellular organization

#### Col33 primary cell line (CRC)

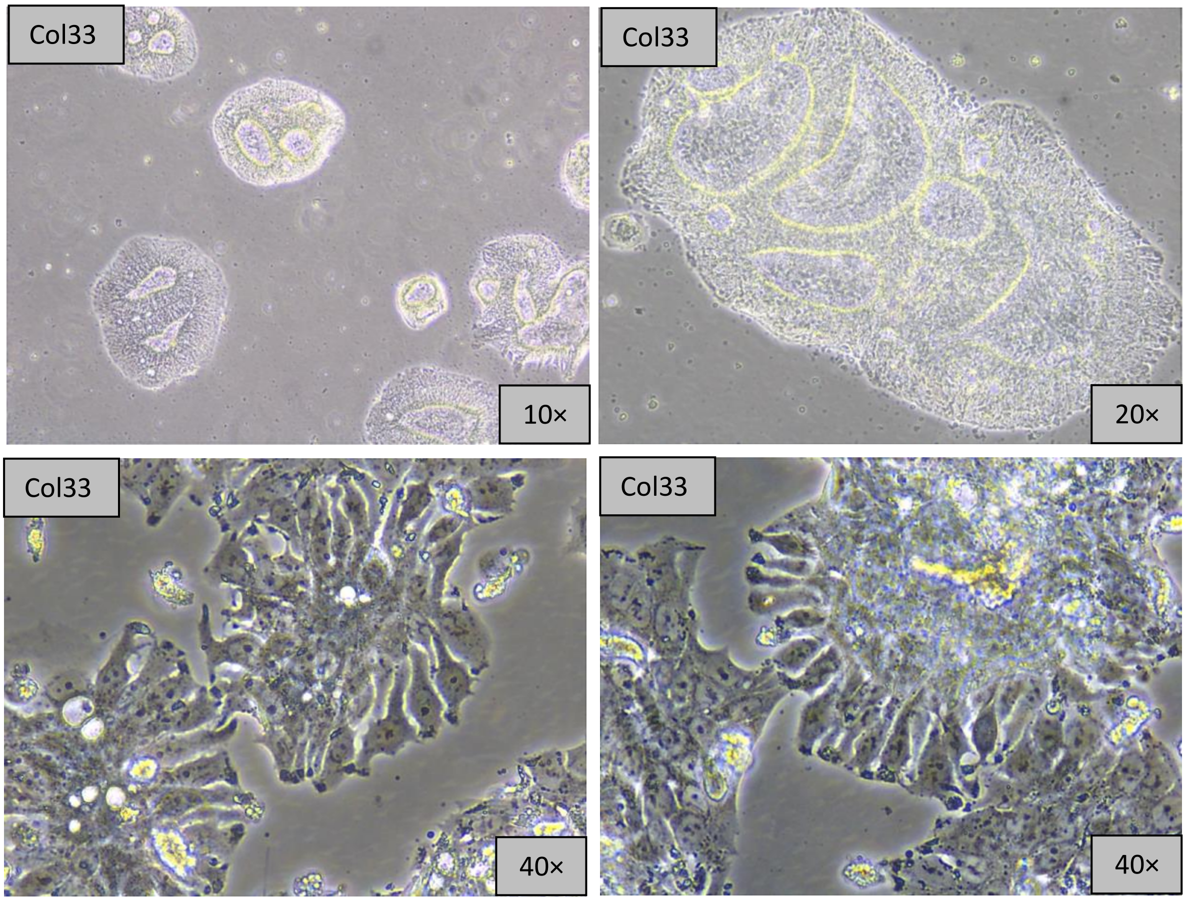


**Morphology and cellular organization of the primary CRC cell line Col33**: Cells were isolated from a colorectal carcinoma (CRC) surgical resection and cultured in DMEM/F12 supplemented with 10% fetal calf serum (FCS) and 1% penicillin-streptomycin-amphotericin B (PSA). In two-dimensional (2D) culture, the cells exhibit a radial arrangement and progressively form compact, crypt-like structures characterized by a consistent central lumen. Although these structures do not extend longitudinally as classical tubular crypts, the presence of an internal space surrounded by radially organized cells indicates that the cells retain elements of intrinsic cooperative organizational programs. Col33 cells proliferate rapidly and display high metabolic activity. Morphologically, they are bottle-shaped, with the nucleus positioned at the expanded basal end. Their spatial organization appears to be influenced by the alignment of structures located opposite the basal poles. Light microscopy reveals a pronounced accumulation of granules in the basal regions of the cells, further supporting the presence of a basal–apical polarity axis oriented toward the luminal space. The Col33 primary immortalized cell line therefore mirrors the structural paradox observed in patient crypts: despite lacking expression of canonical lineage markers, these cells spontaneously adopt crypt-like architectures with a defined lumen even under standard 2D culture conditions. This parallel suggests that crypt architecture represents a robust, self-organizing module capable of persisting independently of classical cellular identity. In both contexts -tissue and culture- the niche appears anatomically preserved yet molecularly erased, forming a “ghost crypt” scaffold that may be susceptible to reprogramming by external programs such as EBV-associated factors. The consistency of this phenomenon across systems supports the notion that architectural persistence can coexist with functional plasticity, and that niche reoccupation may follow distinct spatial and molecular logics.

**Col124 primary cell line (CRC)**

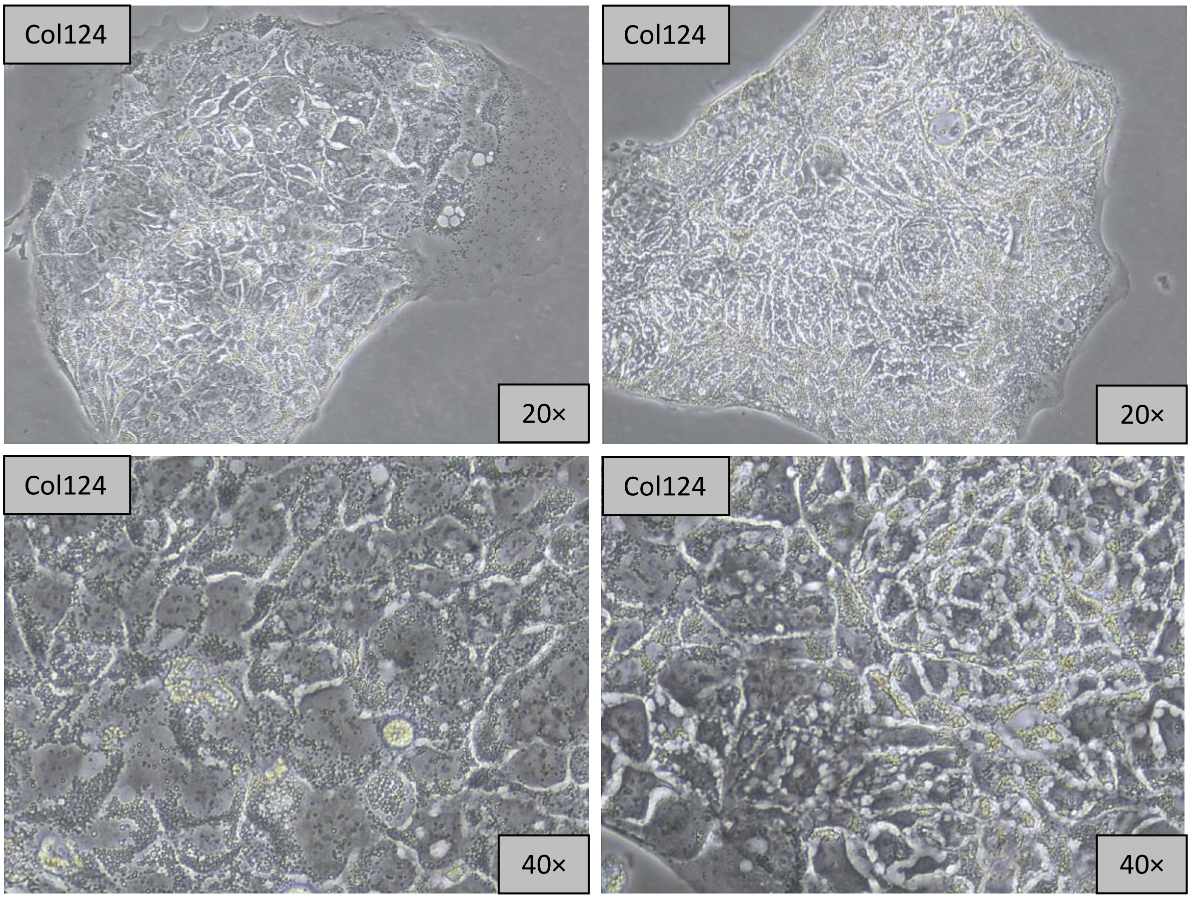


**Morphology and cellular organization of the primary CRC cell line Col124**: Cells were isolated from a colorectal carcinoma (CRC) surgical resection and maintained in DMEM/F12 medium supplemented with 10% fetal calf serum (FCS) and 1% penicillin/streptomycin/amphotericin B (PSA). In two-dimensional (2D) cultures, the cell colonies exhibit a cobblestone-like organization. Col124 cells are characterized by very slow proliferation and low metabolic activity. Notably, the colonies demonstrate strong resistance to enzymatic dissociation, which is presumed to result from high expression levels of Cadherin-17.

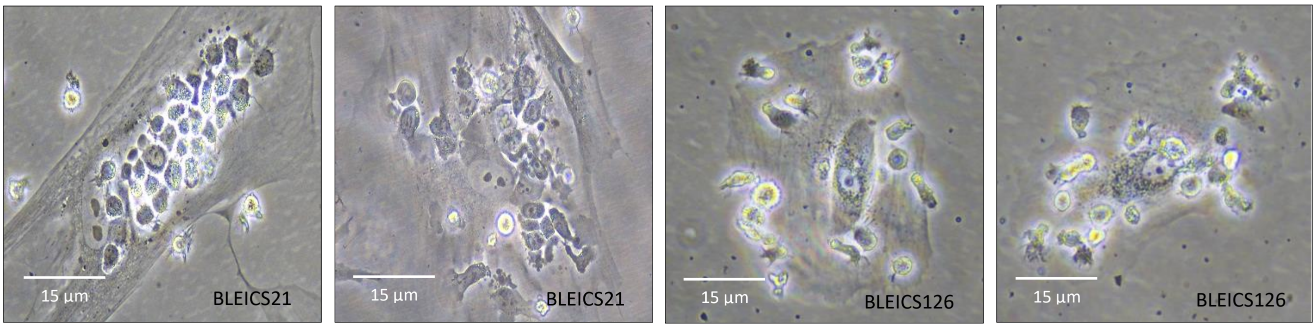


**BLEICS cells interaction with cognate CAF cells**

### SI: Antibodies and Primer

| Antibody name | Clonality | Dilution | Host | Cat No | Manufacturer |
|---|---|---|---|---|---|
| **Primary antibodies** | | | | | |
| ASCL2 | Polyclonal | 1:100 | Rabbit | 21368-1-AP | Proteintech |
| β-Catenin | Polyclonal | 1:100 | Rabbit | 9562 | Cell Signaling |
| CAR | Monoclonal | 1:100 | Rabbit | 86186-1-RR | Proteintech |
| CCR5 | Recombinant | 1:1000 | Rabbit | 82941-1-RR | Proteintech |
| CD19 | Monoclonal | 1:500 | Mouse | 66298-1-IG | Proteintech |
| CD19 | Polyclonal | 1:100 | Rabbit | 27949-1-AP | Proteintech |
| CD22 | Polyclonal | 1:100 | Rabbit | 21894-1-AP | Proteintech |
| CD22 | Monoclonal | 1:10000 | Mouse | 66103-1-IG | Proteintech |
| CD45 | Polyclonal | | Rabbit | 20103-1-AP | Proteintech |
| CD45 | Monoclonal | 1:200 | Mouse | 304002 | Biolegend |
| CD 54 ICAM | Polyclonal | 1.100 | Rabbit | 10831-1-AP | Proteintech |
| CD80 | Monoclonal | 1:100 | Mouse | 66406-1-IG | Proteintech |
| CD133 | Polyclonal | 1:200 | Rabbit | 64326 | Cell Signaling |
| Chromogranin A | Monoclonal | 1:2000 | Mouse | A3 | DAKO |
| Chromogranin A | Polyclonal | 1:1000 | Rabbit | 10529-1-AP | Proteintech |
| CLMP | Polyclonal | 1:500 | Rabbit | 16127-1-AP | Proteintech |
| CXCR4 | Monoclonal | 1:1000 | Mouse | 60042-1-IG | Proteintech |
| DCLK1 | Polyclonal | 1:200 | Rabbit | 21699-1-AP | Proteintech |
| DEFA6 | Polyclonal | 1:100 | Rabbit | 17923-1-AP | Proteintech |
| DNABJ11 | Polyclonal | 1:1000 | Rabbit | 15484-1-AP | Proteintech |
| EPHA2 | Monoclonal | 1:200 | Mouse | 66736-1-IG | Proteintech |
| GLUT3 | Polyclonal | 1:2000 | Rabbit | 20403-1-AP | Proteintech |
| GP2 | Polyclonal | 1:100 | Rabbit | CAU22808 | Biomatik |
| GP2 | Polyclonal | 1:200 | Rabbit | HPA015739 | Atlas antibodies |
| IgG LC Kappa | Monoclonal | 1:1000 | Mouse | MA1-10385 | ThermoFisher |
| IgG LC Lambda | Monoclonal | 1:1000 | Mouse | 604-611 | ThermoFisher |
| KRT80 | Polyclonal | 1:200 | Rabbit | 16835-1-AP | Proteintech |
| LBP | Monoclonal | 1:250 | Mouse | 66181-1-IG | Proteintech |
| LGR5 | Polyclonal | 1:100 | Rabbit | 30007-1-AP | Proteintech |
| MHCII Pan | Monoclonal | 1:10 -1:25 | Mouse | Sc53302 | Santa Cruz |
| NGR1 | Polyclonal | 1:300 | Rabbit | 27143-1-AP | Proteintech |
| NSE | Monoclonal | 1:25 | Mouse | 500-6754 | ThermoFisher |
| NT3 | Polyclonal | 1:1000 | Rabbit | Orb11171 | biorbyt |
| Peripherin | Monoclonal | 1:100 | Mouse | 66317-1-IG | Proteintech |
| PigR | Polyclonal | 1:100 | Rabbit | 22024-1-AP | Proteintech |
| PyY | Polyclonal | 1:300 | Rabbit | 24294-1-AP | Proteintech |
| REG4 | Polyclonal | 1:500 | Rabbit | 12268-1-AP | Proteintech |
| REST | Polyclonal | 1:200 | Rabbit | 22242-1-AP | Proteintech |
| S100A4 | Polyclonal | 1:1000 | Rabbit | 16105-1-AP | Proteintech |
| SDF1 | Polyclonal | 1:100 | Rabbit | 17402-1-AP | Proteintech |
| Somatostatin | Polyclonal | 1:100 | Rabbit | 24496-1-AP | Proteintech |
| SOX9 | Monoclonal | 1:100 | Mouse | 67439-1-IG | Proteintech |
| SpiB | Polyclonal | 1:1000 | Rabbit | 15768-1-AP | Proteintech |
| Stathmin | Monoclonal | 1:100 | Rabbit | 13655 | Cell Signaling |
| Stathmin | Polyclonal | 1:100-1:1000 | Rabbit | 11157-1-AP | Proteintech |
| TH | Polyclonal | 1:500 1:5000 | Rabbit | 25859-1-AP | Proteintech |
| TLR3 | Recombinant | 1:1000 | Rabbit | 83136-3-RR | Proteintech |
| TLR4 | Monoclonal | 1:100 | Mouse | 66350-1-IG | Proteintech |
| TNFaIP2 | Polyclonal | 1:1000 | Rabbit | 25649-1-AP | Proteintech |
| TNFaIP2 | Polyclonal | 1:100 IHC | Rabbit | STJ96055 | St Johns |
| Uromodulin | Polyclonal | 1:500 | Rabbit | 11911-1-AP | Proteintech |
| Viperin /RSAD2 | Polyclonal | 1:500 | Rabbit | 28089-1-AP | Proteintech |
| ZG16 | Polyclonal | 1:50 | Rabbit | 17397-1-AP | Proteintech |
| ZG16 | Monoclonal | 1:250 | Mouse | 67389-1-IG | Proteintech |
| **Secondary antibodies** | | | | | |
| Anti-β-Actin-HRP | Monoclonal | 1:20000 | Rabbit | A3854 | Sigma Aldrich |

| | | | | | |
|---|---|---|---|---|---|
| Anti-GAPD-HRP | Monoclonal | 1:20000 | Mouse | Sc-47724 | Santa Cruz |
| Anti-Goat-Alexa 568 | Polyclonal | 1:1000 | Donkey | Ab175704 | Abcam |
| Anti-Goat-FITC IgG | Polyclonal | 1:1000 | Rabbit | ABIN101210 | Antibodies-online |
| Anti-Goat-HRP IgG | Polyclonal | 1:1000 | Donkey | Sc-2020 | Santa Cruz |
| Anti-Mouse-HRP IgG | N/A | 1:1000 | Horse | 7076 | Cell Signaling |
| Anti-Mouse IgG Alexa 488 | Polyclonal | 1:2000 | Goat | 4408 | Cell Signaling |
| Anti-Mouse IgG Alexa 555 | Polyclonal | 1:2000 | Goat | 4409 | Cell Signaling |
| Anti-Rabbit-HRP IgG | N/A | 1:1000 | Goat | 7074 | Cell Signaling |
| Anti-Rabbit IgG Alexa 488 | Polyclonal | 1:2000 | Goat | 4412 | Cell Signaling |
| Anti-Rabbit IgG Alexa 555 | Polyclonal | 1:2000 | Goat | 4413 | Cell Signaling |
| **Coupled FACS antibodies** | | | | | |
| CCR5 (APC) | Monoclonal | 1:50 | Rat | 359121 | Biolegend |
| CD3 (PacificBlue) | Monoclonal | 1:50 | Mouse | 344824 | Biolegend |
| CD4 (Alexa488) | Monoclonal | 1:50 | Mouse | 300519 | Biolegend |
| CD8 (BrilliantViolet) | Monoclonal | 1:50 | Mouse | 301048 | Biolegend |
| CD19 (APC) | Monoclonal | 1:50 | Mouse | 302212 | Biolegend |
| CD28 (PE) | Monoclonal | 1:50 | Mouse | 302908 | Biolegend |
| CD45 (APC) | Monoclonal | 1:50 | Mouse | 368517 | Biolegend |
| CD56 (PE) | Monoclonal | 1:50 | Mouse | 362524 | Biolegend |
| CD158 (APC) | Monoclonal | 1:50 | Mouse | 339509 | Biolegend |
| CTLA-4 (PE) | Monoclonal | 1:50 | Mouse | 369604 | Biolegend |
| CXCR3 (PE) | Monoclonal | 1:50 | Mouse | 353706 | Biolegend |
| CXCR4 (BrilliantViolet) | Monoclonal | 1:50 | Mouse | 306518 | Biolegend |
| HLA-DR (BrilliantViolet) | Monoclonal | 1:50 | Mouse | 307646 | Biolegend |
| PD-1 (APC) | Monoclonal | 1:50 | Mouse | 329908 | Biolegend |

**Table 3: Antibodies -** List of antibodies used for ICC, IHC, FACS and Western blot analysis

| Name | Forward | Probe | Reverse |
|---|---|---|---|
| 18S (NR_003286) | GGACATCTAAGGCATCACAG | TGCTCAATCTCGGGTGGCTGAA | GAGACTCTGGCATGCTAACTAG |
| ACHE (M55040.1) | CTCAACGTGTGGACACCATA | CCCTGTCCTCGTCTGGATCTATGGG | CTCCTTGGACGTGTACGATG |
| ALDH1L1 (NM_001270364.2) | GAAGGATGGAGTGCCGGTAT | TTTGCCACCACATCAGGCAAAGC | GCTGCAGAAGGGCAGGA |
| ASCL1 (NM_004316.4) | AAACGCCGGCTCAACTT | TTGACCAACTTGACGCGGTTGC | CCCGAAGGGTGGCAAAG |
| β III Tubulin (NM_006086.3) | GGGCCAAGTTCTGGGAAG | TCAGTGATGAGCATGGCATCGACC | GGATCAGCGTCTACTACAACG |
| BDNF (NM_170735.6) | CTCTTTCTGCTGGAGGAATACAA | ATGCTGCAAACATGTCCATGAGGG | GCCGTTACCCACTCACTAATAC |
| c-Myc (NM_002467.6) | CTGAGGAGGAACAAGAAGATGA G | AGAGTCTGGATCACCTTCTGCTGGA | TGTGAGGAGGTTTGCTGTG |
| CD24 (NM_001291737.1) | CAAATCCAACTAATGCCACCAC | TCGTGGTCTCACTCTCTCTTCTGCA | GTTTCTTGGCCTGAGTCTCTT |
| CD133 (NM_006017.3) | GAGGCGTTGGAGAACATGAA | ACACAGCTTAGCAGCAGTCTGACC | CACAGAGGGTCATTGAGAGATG |
| Choline O-acetyltransferase (NM_001142934.1) | AAAGGCCAGCTGTCAGG | ACTATGGGCTCTTCTCCTCCTACCG | CATACCCAAGGACACGCTG |
| Chromogranin A (J03483.1) | GGTCATCTCCGACACACTTTC | AGACACTCCGAGGAGATGAACGGA | GTCTTGGAGCTCCTTCAGTAAAT |
| Doublecortin (AJ003112.1) | CTGTGTCCTCTGACCGTTT | TGACCTGACGCGATCTCTGTCTGA | CTCAGGGAGTGCGTTACATT |
| EBV EBNA1 (NC_007605.1) | CCCTGCACCCAGTACCT | CCAGCGGCCATTCTCTCTGGTAAC | CGGGCCTCTACTTCTTCTCT |
| EBV LMP1 (ON596484.1) | CTTTGGCTCCTCCTGTTTCT | ATGAACACCACCACGATGACTCCC | |
| EBV N Capsid (NC_007605.1) | AACGCGAGGAGAAGGTTATTC | TGTTTACGATCCATGCCTCCACCG | TATGCCCAATCCCAAGTACAC |

| | | | |
|---|---|---|---|
| E-cadherin (Z13009.1) | CAGAAGACAGAAGAGAGACTGG | GTTATTCCTCCCATCAGCTGCCCA | TCCAACAAAGACAAAGAAGGC |
| EIF2AK2 (NM_002759) | TGGAAAGCGAACAAGGAGTAAG | TGCGATACATGAGCCCAGAACAGATTT | CAGCAAGAATTAGCCCCAAAG |
| EpCAM (NM_002354.3) | GGTGATGAAGGCAGAAATGAATG | CCCATCATTGTTCTGGAGGGCC | TCATCGCAGTCAGGATCATAAAG |
| Ephrin B1 (NM_004429.5) | GACTGTGAACCAGGAAGAGAAG | ACCCTGATGGCTTCTTCAACTCCA | CCGTCAGGAAGATGATGATGAG |
| ERV 3.1 (NM_001007253) | CCTCCCTTGGCCAGTAATTTAT | ACATAACATGAAGCAACGTGCAGGC | TTGGTCTCCCATGTTCATTCC |
| ERVMER 34.1 SU (NM_024534.6) | GGCCTTCTGTACCAGCTATTT | TGACAGAGGCACATGGGAAATGGA | CGGTGTCCATCATGACCTTT |
| GAD67 (NM_000817.3) | CCTTCAGGGAGAGGCAATC | CTCCAAGAACCTGCTTTCCTGTGA | TTAGAGAAGTCAGTCTCTGTGC |
| GDNF (NM_000514.4) | CTTCCTAGAAGAGAGCGGAATC | TGCCAACCCAGAGAATTCCAGAGG | AGACCCAAGTCAGTGACATTTA |
| Goosecoid (NM_173849.3) | GGACTATGGCGCCTTCT | TCCCGCCTCGGCTACAACAA | CAGCTGCCCGTAGAAGTA |
| HERV Fc1 env (XM_047447119) | AATATGGCAGGCCTAGTTAATCTC | CAGCCAACCACCACTAGTAGCTGT | GGTGGGCAGTGTGGTTAC |
| HERV H env (AJ289710.2) | GGTAATCTTTCACCTTCTCGATGT | CCGAAGCCCAACTACACACATCACT | CAGTATTGATAGAGGGCTTGTCTG |
| HERV H62 SU (AJ289709.1) | CACTCCATCCTTGGCTATCTTC | TGCTCATCAGACTCTCCTCCCAGG | CGTCGAGTATCTACGAGCAATC |
| HERV K env (DQ069916.1) | CCACAATTGCTCAGGACAAAC | TGTCCAAGTGCACAAGTGAGTCCA | CCCATTCCCAAGGGTAGAAAG |
| HERV K SU (X82272) | CAATGGTGGTAAGTCTCCCTATG | CTACTGGGCCTATGTGCCTTTCCC | CCATGTGACTGCCCGAATTA |
| HERV K18 SU (Y18890.1) | ACCAGCCTCCCAAACTATAAC | AATTGGCAGCACCGTATTCT | ATGCCTTCTCTTGCTCTCAC |
| HERV T SU (AB266802) | CTTCCATAAGCACCTCAGTCTC | ACCAAGCACCCAACAATACCTGGT | GGTCCTGGTTCAGTTCCATTA |
| HES1 (Y07572.1) | GGGAAAGATTGCAAGGTGAATAAA | CGTGTCCTGAAGGAGTTCCACCAG | CTTGTGCTGCATTGCACC |
| HES5 (DQ272660.1) | CAGCATCGAGCAGCTGAA | CTGAAGCACAGCAAAGCCTTCGTC | TACAGCGAAGGCTACTCGT |
| HIF1a (NM_001530.4) | ACTCAGTTTGAACTAACTGGACA | ACTCATCCATGTGACCATGAGGAAATGAG | CAAGGCCATTTCTGTGTGTAAG |
| HLA DRA (NM_019111) | TCAGGAATCATGGGCTATCAAAG | AGGATTCAGATAGAACTCGGCCTGGA | AGTCAAACATAAACTCGCCTGA |
| KLF4 (NM_001314052.2) | AGGAGCCCAAGCCAAAG | TAATCACAAGTGTGGGTGGCGGTC | GCCTTGAGATGGGAACTCTT |
| MAP2 (NM_002374.3) | GAAGATAAATCAGGAATGTCCAAG | ACATCTGCCTTGAAAGAAGAAGCAACA | AGGCAGTGATTCTATGAACTGA |
| Nanog (NM_024865.4) | CAGGACAGCCCTGATTCTTC | CAGTCCCAAAGGCAAACAACCCAC | GTTTCTTGACCGGGACCTT |
| N-cadherin (X57548.1) | CTTCATGCCGGTACCATGT | ACAACATTCACTGCTCAGGACCCA | AAATTATCTGATCCTGCCAATTGG |
| NDF1 (AB593070.1) | GAAGAGGAGGATGACGATCAAA | AGAAGATGACTAAGGCTCGCCTGGA | ATTGAGACGCATGAAGGCT |
| Nestin (NM_006617.2) | AGCGTTGGAACAGAGGTTG | AAAGTTCCAGCTGGCTGTGGAGG | TGTAGGCCCTGTTTCTCCT |
| Neurog2 (NM_024019.4) | CTGGGTCTGGTACACGATTG | CTTCTTGATGCGCTGCACCGTCTC | GGTTGTTGGCCTTCAGTCTA |
| Neurturin (NM_004558.5) | AGAGGGCCTGCTTCTCA | CAGTACCGTGCACTCCTGCA | GCAGCTCCATCGCATCC |
| NGF (NM_002506.3) | GCAGACCCGCAACATTACT | TGCTGTTTAGCACCCAGCCTCC | GACCTCGAAGTCCAGATCCT |
| NMDAR1 (NM_007327.3) | GATGGCTCTGT CGGTGTG | AGGTCTACGCCA TCCTAGTTAGCCA | CCAACGACCAC TTCACTCC |

| | | | |
|---|---|---|---|
| NMDAR2B (U88963.1) | GATGCCCACGA GAAAGATGA | AGCCATGAATGA GACCGACCCAAA | CATCATCACC CGCATCTGT |
| Notch1 (NM_017617.5) | CAGTGTCGAGATGGCTATGA | TGTGTGTTACCTACCACAATGGCACA | CCTTCTGGACATTTGCAGTATC |
| NSE (NM_001975.3) | ACAGTGGAGGTGGATCTCTAT | TGAGGGATGGAGACAAACAGCGTT | CAGGACACCTTTGCCTAAGT |
| Nurr1 (NM_006186.3) | CCAGCTTCAGTA CCTTTATGGA | AGCCACCTTGCTTGTACCAAATGC | CAGCAGTCCTCCATTAAGGTAG |
| OAS1 (NM_016816) | GGTGGAGTTCGATGTGCTG | CCTTTGATGCCCTGGGTCAGTTGA | GATGAGCTTGACATAGATTTGGG |
| OAS2 (NM_016817) | GAAATCCAAAGTCCTCAACGAAAG | CAGCTGACCCAGTGCATTAAAGGC | CAGATCAATGAGCCCTGCATA |
| OAS3 (NM_006187) | GTGTGGACTTTGATGTGCTG | TAGACTTGAGAGCTGGGCCTGGA | CTGTAGCTGTGGATGAGGTC |
| OASL (NM_198213) | GGGTGCTGAAGGTAGTCAAG | TGCTCCTGAGAACCGTGCCATT | GTGGAAACAGCTCAGAAACG |
| Occludin (NM_002538.3) | GCAATGACATATATGGTGGAGAGA | GTTCGACCAATGCTCTCTCAGCCA | TACAAATGGACCTCTCCTCCA |
| Oct-4 / POU5F1 (NM_002701.6) | GCTTAGCTTCAAGAACATGTGTA | TGGGTGGAGGAAGCTGACAACAAT | CACGAGGGTTTCTGCTTTG |
| Pax6 (NM_000280.4) | AGGCTCAAATGCGACTTCA | GCTGAAGCGGAAGCTGCAAAGAAA | GAGAAAGAGTTTGAGAGAACCC |
| PRKRA (NM_003690) | GAAGACCAAGAACATCCCAGT | ACGTGCCCACTTTCACCTTCAGA | GCCAGCTTCTTACTTGTACCT |
| PSD95 (U83192.1) | GTGCTGCCCACAAGGAT | GGGAGGTTGCAGATTGGAGACAAGA | TGAAGATGCTGTGGCAGC |
| RNASEL (NM_021133) | AGCAGTCTTCCAGGCTTTG | TGAGTTGAGCAGGTGGAATGTCAGAAG | CAACAGAGCAGCAGTATGAAGA |
| SCG5 (NM_001144757.3) | AGGCTGCTTCATGGTGTTAT | AGCTCACCAGGCCATGAATCTTGT | GCTCCACCTTCAATGCTCT |
| SKIV2L (NM_006929) | CATCCTGCCCATCCTCAAG | CGTGGCCTGGTCAAGGTCTTGTT | TGGAGTCAAACACTACTGTACG |
| SNAIL (NM_005985.4) | GCAGGACTCTAATCCAGAGTTTAC | ATCCCACCTCCGGAGATCCTCAA | AGGACAGAGTCCCAGATGAG |
| SOX2 (NM_003106.4) | CGGACAGCGAACTGGAG | AGAGGAGAGTAAGAAACAGCATGGAGA | TTTGAGCGTACCGGGTTT |
| Stathmin (X53305.1) | GAGCACGAGAAAGAAGTGCT | GAAGGCAATAGAAGAGAACAACAACTTC | CAGAAGAGAAACTGACCCACAA |
| SYN1 SU (NM_014590.4) | GTCACTGTCTGTTGGACTTACT | TCTTGCCTGATCTTGAACTCCACCC | CGGGTGAGTTGGGAGATTAC |
| SYN2 SU (NM_207582.3) | GTAGGCACTCTTCCAAGTACAG | ACCAACAGACTTACCAAACATACACCCA | CTTGGTTGATGGCGGAATTG |
| Synaptophysin (NM_003179.2) | CAGGCTGCACCAAGTGTA | GGGCACCACCAAGGTCTTCTTAGTT | GTCAGCCGAATTCTTTGTCAC |
| TARBP2 (NM_134324) | CCTGTGTACGACCTTCTCAAAG | AGCCCACCAGCCTAATTTCACCT | CTCAGCTGCCTTGTGCTT |
| TBR1 (NM_006593.4) | ACTGGATGCGCCAAGAAA | CCACCATCTGCCCATTGTTATTTGAAGC | GGTACTTGTGCAAGGACTGTAA |
| TBR2 (NM_001278182.1) | ACCAATAACAAAGGCGCAAATA | ACAACACCCAGATGATAGTCTTACAATCC | CATATTGTTGAAGTTACAGAGGA |
| Tenascin-C (NM_002160.3) | CATCTGTGACGACGGCTT | CAATGACCAGGGCAAGTGCGTAAA | GGAGTCTGCATC TGTTTCGA |
| V1/V2 SU (NM_001191055.2) | GGATCTCACTCCACCTTTCAAT | AAACCAAACCTGTTGTCCATGCCC | CTTCCTGTCACAGTGTCCTTC |
| Vimentin (NM_003380.5) | CTCAATGTCAAGGGCCATCT | ACGAAGGTGACGAGCCATTTCCTC | GCTGCTAACTACCAAGACACTAT |

**Table 4: qPCR primers and probes -** Primer and probe sequences for qPCR designed with IDT PrimerQuest$^{TM}$ Tool. All primers and probes are reflected in 5´→3´ polarity

**SI: REST CRISPR/Cas9 ablation**

**a.** REST gene and design of gRNAs, primers and probes for analysis of gene ablation. NCBI Reference Sequence: NM_001363453.3 Homo sapiens RE1 silencing transcription factor (REST)

gRNA A

REST crRNA A CCATGTGAGCAAGTACAATTTGG neg Strang( IDT)

REST crRNA A CCAAATTGTACTTGCTCACATGG pos Strang

qPCR gRNA A

se TGGAGGTGGTTCAGGAG

as AGCTCCATGTGAGCAAGTA rev kompl TACTTGCTCACATGGAGCT

Sonde CTCAGATGGTGGGTGCCCAAATTG

T7 gRNA A TM cal NEB 65°C Annealing Temp. **848 bp**

FW: GAC AGC AAA GTG GAG GAG AAT A

REV: GGC ACT AAG CCA ACT TCA ATA AG

gRNA B

REST crRNA B CGTATTCAAGTTTTGTCCAGAGG neg Strang (IDT)

REST crRNA B CCTCTGGACAAAACTTGAATACG pos Strang

qPCR gRNA B

se CTACCTGGTCTTGCTGCTAATA

as CTGTCAGTCTGATGTTTACCATTT rev kompl AAATGGTAAACATCAGACTGACAG

Sonde ACTTGAATACGCCAGAGGGTGAAACT

T7 gRNA B TM cal NEB 65°C Annealing Temp. **915 bp**

FW:: CCTGTTCAGATGGAGTTGTCTC

REV**:** CGATTGAGGTGTTTGCTGTAATC

```
ATGGCCACCCAGGTAATGGGGCAGTCTTCTGGAGGAGGAGGGCTGTTTACCAGCAGTGGCAACATTGGAATGGCC
CTGCCTAACGACATGTATGACTTGCATGACCTTTCCAAAGCTGAACTGGCCGCACCTCAGCTTATTATGCTGGCA
AATGTGGCCTTAACTGGGGAAGTAAATGGCAGCTGCTGTGATTACCTGGTCGGTGAAGAAAGACAGATGGCAGAA
CTGATGCCGGTTGGGGATAACAACTTTTCAGATAGTGAAGAAGGAGAAGGACTTGAAGAGTCTGCTGATATAAAA
```

GGTGAACCTCATGGACTGGAAAACATGGAACTGAGAAGTTTGGAACTCAGCGTCGTAGAACCTCAGCCTGTATTT
GAGGCATCAGGTGCTCCAGATATTTACAGTTCAAATAAAGATCTTCCCCCTGAAACACCTGGAGCGGAGGACAAA
GGCAAGAGCTCGAAGACCAAACCCTTTCGCTGTAAGCCATGCCAATATGAAGCAGAATCTGAAGAACAGTTTGTG
CATCACATCAGAGTTCACAGTGCTAAGAAATTTTTTGTGGAAGAGAGTGCAGAGAAGCAGGCAAAAGCCAGGGAA
TCTGGCTCTTCCACTGCAGAAGAGGGAGATTTCTCCAAGGGCCCCATTCGCTGTGACCGCTGCGGCTACAATACT
AATCGATATGATCACTATACAGCACACCTGAAACACCACACCAGAGCTGGGGATAATGAGCGAGTCTACAAGTGT
ATCATTTGCACATACACAACAGTGAGCGAGTATCACTGGAGGAAACATTTAAGAAACCATTTTCCAAGGAAAGTA
TACACATGTGGAAAATGCAACTATTTTTCAGACAGAAAAAACAATTATGTTCAGCATGTTAGAACTCATACAGGA
GAACGCCCATATAAATGTGAACTTTGTCCTTACTCAAGTTCTCAGAAGACTCATCTAACTAGACATATGCGTACT
CATTCAGGTGAGAAGCCATTTAAATGTGATCAGTGCAGTTATGTGGCCTCTAATCAACATGAAGTAACCCGCCAT
GCAAGACAGGTTCACAATGGGCCTAAACCTCTTAATTGCCCACACTGTGATTACAAAACAGCAGATAGAAGCAAC
TTCAAAAAACATGTAGAGCTACATGTGAACCCACGGCAGTTCAATTGCCCTGTATGTGACTATGCAGCTTCCAAG
AAGTGTAATCTACAGTATCACTTCAAATCTAAGCATCCTACTTGTCCTAATAAAACAATGGATGTCTCAAAAGTG
AAACTAAAGAAAACCAAAAAACGAGAGGCTGACTTGCCTGATAATATTACCAATGAAAAAACAGAAATAGAACAA
ACAAAAATAAAAGGGGATGTGGCTGGAAAGAAAAATGAAAAGTCCGTCAAAGCAGAGAAAAGAGATGTCTCAAAA
GAGAAAAAGCCTTCTAATAATGTGTCAGTGATCCAGGTGACTACCAGAACTCGAAAATCAGTAACAGAGGTGAAA
GAGATGGATGTGCATACAGGAAGCAATTCAGAAAAATTCAGTAAAACTAAGAAAAGCAAAAGGAAGCTGGAAGTT
GACAGCCATTCTTTACATGGTCCTGTGAATGATGAGGAATCTTCAACAAAAAAGAAAAAGAAGGTAGAAAGCAAA
TCCAAAAATAATAGTCAGGAAGTGCCAAAGGGTGACAGCAAAGTGGAGGAGAATAAAAAGCAAAATACTTGCATG
AAAAAAAGTACAAAGAAGAAAACTCTGAAAAATAAATCAAGTAAGAAAAGCAGTAAGCCTCCTCAGAAGGAACCT
GTTGAGAAGGGATCTGCTCAGATGGACCCTCCTCAGATGGGGCCTGCTCCCACAGAGGCGGTTCAGAAGGGGCCC
GTTCAGGTGGAGCCGCCACCTCCCATGGAGCATGCTCAGATGGAGGGTGCCCAGATACGGCCTGCTCCTGACGAG
CCTGTTCAGA**TGGAGGTGGTTCAGGAG**GGGCCTGCTCAGAAGGAGCTGCTGCCTCCCGTGGAGCCTG**CTCAGATG**
**GTGGGTGCCCAAATTGTACTTGCTCACATGGAGCT**GCCTCCTCCCATGGAGACTGCTCAGACGGAG
GTTGCCCAAATGGGGCCTGCTCCCATGGAACCTGCTCAGATGGAGGTTGCCCAGGTAGAATCTGCTCCCATGCAG
GTGGTCCAGAAGGAGCCTGTTCAGATGGAGCTGTCTCCTCCCATGGAGGTGGTCCAGAAGGAGCCTGTTCAGATA
GAGCTGTCTCCTCCCATGGAGGTGGTCCAGAAGGAACCTGTTAAGATAGAGCTGTCTCCTCCCATAGAGGTGGTC
CAGAAGGAGCCTGTTCAGATGGAGTTGTCTCCTCCCATGGGGGTGGTTCAGAAGGAGCCTGCTCAGAGGGAGCCA
CCTCCTCCCAGAGAGCCTCCCCTTCACATGGAGCCAATTTCCAAAAAGCCTCCTCTCCGAAAAGATAAAAAGGAA
AAGTCTAACATGCAGAGTGAAAGGGCACGGAAGGAGCAAGTCCTTATTGAAGTTGGCTTAGTGCCTGTTAAAGAT
AGCTGGCTTCTAAAGGAAAGTGTAAGCACAGAGGATCTCTCACCACCATCACCACCACTGCCAAAGGAAAATTTA
AGAGAAGAGGCATCAGGAGACCAAAAATTACTCAACACAGGTGAAGGAAATAAAGAAGCCCCTCTTCAGAAAGTA
GGAGCAGAAGAGGCAGATGAGAGC**CTACCTGGTCTTGCTGCTAATA**TCAACGAATCTACCCATATTTCAT**CCT**
**CTGGACAAAACTTGAATACG**CCAGAGGGTGAAACTTT**AAATGGTAAACATCAGACTGACAG**TATAGT
TTGTGAAATGAAAATGGACACTGATCAGAACACAAGAGAGAATCTCACTGGTATAAATTCAACAGTTGAAGAACC
AGTTTCACCAATGCTTCCCCCTTCAGCAGTAGAAGAACGTGAAGCAGTGTCCAAAACTGCACTGGCATCACCTCC
TGCTACAATGGCAGCAAATGAGTCTCAGGAAATTGATGAAGATGAAGGCATCCACAGCCATGAAGGAAGTGACCT
AAGTGACAACATGTCAGAGGGTAGTGATGATTCTGGATTGCATGGGGCTCGGCCAGTTCCACAAGAATCTAGCAG
AAAAAATGCAAAGGAAGCCTTGGCAGTCAAAGCGGCTAAGGGAGATTTTGTTTGTATCTTCTGTGATCGTTCTTT
CAGAAAGGGAAAAGATTACAGCAAACACCTCAATCGCCATTTGGTTAATGTGTACTATCTTGAAGAAGCAGCTCA
AGGGCAGGAGTAA

**b.** Results of the CRISPR/Cas9 T7 digestion products in HCT8$^{\Delta REST}$ cells separated by 2% agarose gel.

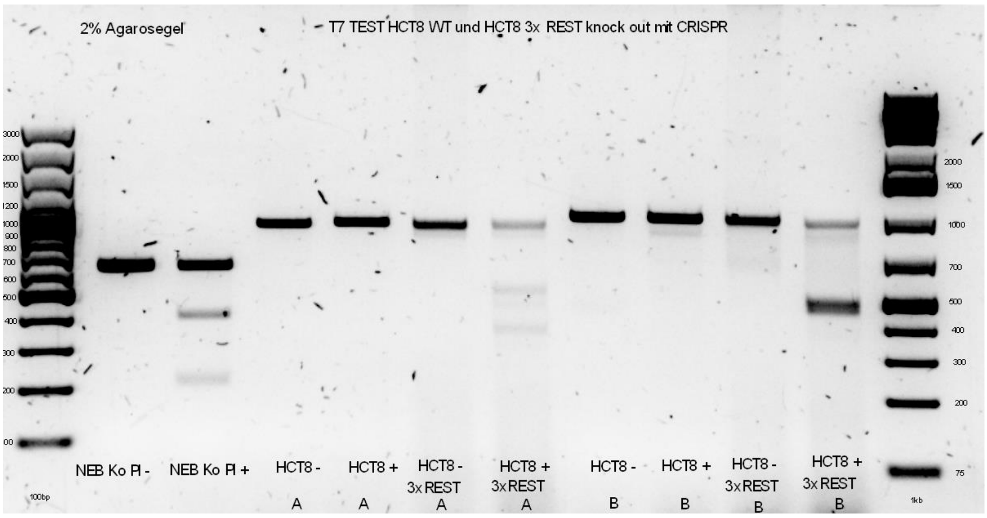


**Lanes**: 1: NEB control plasmid Ø T7 digestion; 2: NEB control plasmid T7 digestion (full length 669, cut 1: 198 bp, cut 2: 471); 3: HCT8$^{WT}$ (gRNA-A) Ø T7 digestion; 4: HCT8$^{WT}$ (gRNA-A) T7 digestion; 5: HCT8$^{\Delta REST}$ 3× treated with gRNA-A Ø T7 digestion; 6: HCT8$^{\Delta REST}$ 3× treated with gRNA-A T7 digestion; 7: HCT8$^{WT}$ (gRNA-B) Ø T7 digestion; 8: HCT8$^{WT}$ (gRNA-B) T7 digestion; 9: HCT8$^{\Delta REST}$ 3× treated with gRNA-B Ø T7 digestion; 10: HCT8$^{\Delta REST}$ 3× treated with gRNA-B T7 digestion.

**c.** Successful knock out of REST gene as assessed by qPCR in CRISPR/Cas9-modified HCT8 cells.

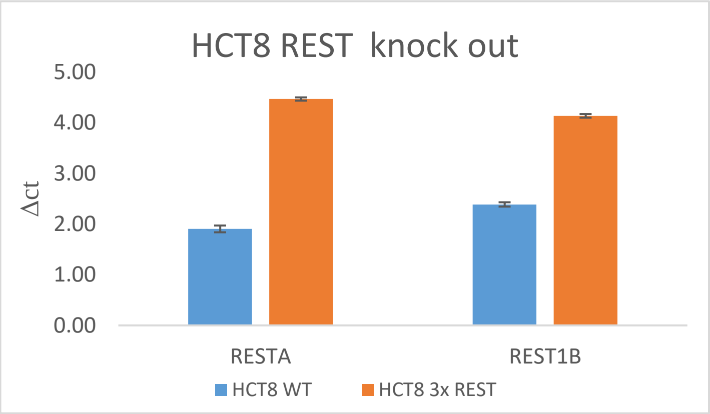

**d.** Westernblot with HCT8 wt and 10 HCT8 ΔREST clones tested for REST protein

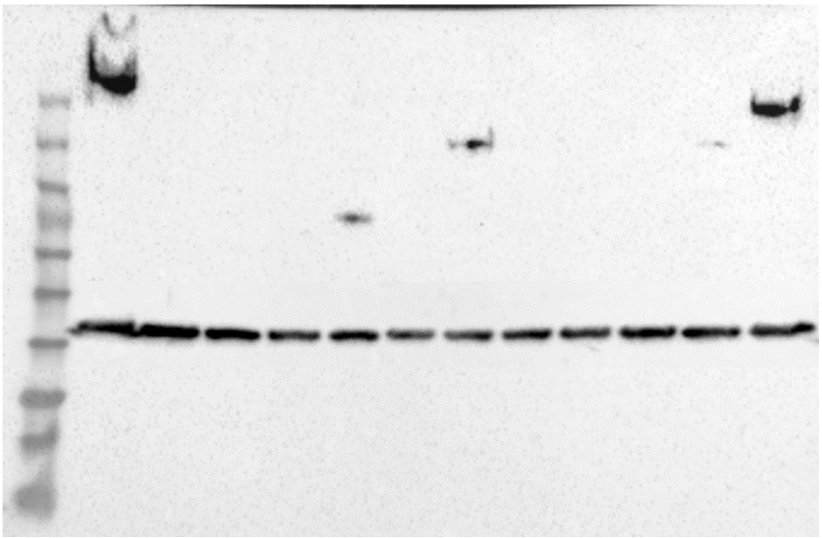


**a**. TLR7 gene and design of gRNAs, primers and probes for analysis of gene ablation. NCBI Reference Sequence: NM_001363453.3. Green: Primers for T7 (848 bp); REST gRNA-A. Petrol: Primers for T7 (915 bp); REST gRNA-B. Yellow: Primers for the detection of mutation using gRNA-A, including S, AS and probe. Grey: Primers for the detection of mutation using gRNA-B, including S, AS and probe. **b**. Results of the CRISPR/Cas9 T7 digestion products in SKOV3TLR7-KO cells separated by 2% agarose gel. **c**. Successful knock out of TLR7 gene as assessed by qPCR with gDNA in CRISPR/Cas9-modified SKOV3 cells. **d**. Ten different clones of $HCT8^{\Delta REST}$ were tested on western blot with antibody α REST (200 kDA). $HCT8^{WT}$ protein lysates served as positive controls and GAPDH was employed as loading control.

### SI: Colonic organoid medium

| **Substance** | **final concentration** |
| --- | --- |
| A83-01 | 0.5 µM |
| Albumin | 1% |
| EGF | 100 ng/ml |
| Gastrin | 10 nM |
| HEPES | 10 mM |
| L-glutamine | 1 x |
| N2 | 1 x |
| N-acetylcisteine | 1.25 mM |
| NC21 supplement | 1 x |
| Nicotinamide | 10 mM |
| Noggin | 100 ng/ml |
| Prostaglandin E2 | 10 nM |
| PSA | 1 x |
| R spondin 1 | 100 ng/ml |
| SB202190 | 10 µM |
| Wnt | 10 ng/ml |
| Y27632 | 10 µM |

**Table 5: Colonic organoid medium**- The formulation is designed to support the growth, maintenance, and phenotypic fidelity of tumor epithelial cells in vitro, mimicking the in vivo tumor microenvironment. The medium typically consists of a basal formulation (Advanced DMEM/F12) supplemented with a defined set of growth factors, pathway modulators, and supportive additives

**SI: Toxicity of REST inhibitors X5050 and GSK126**

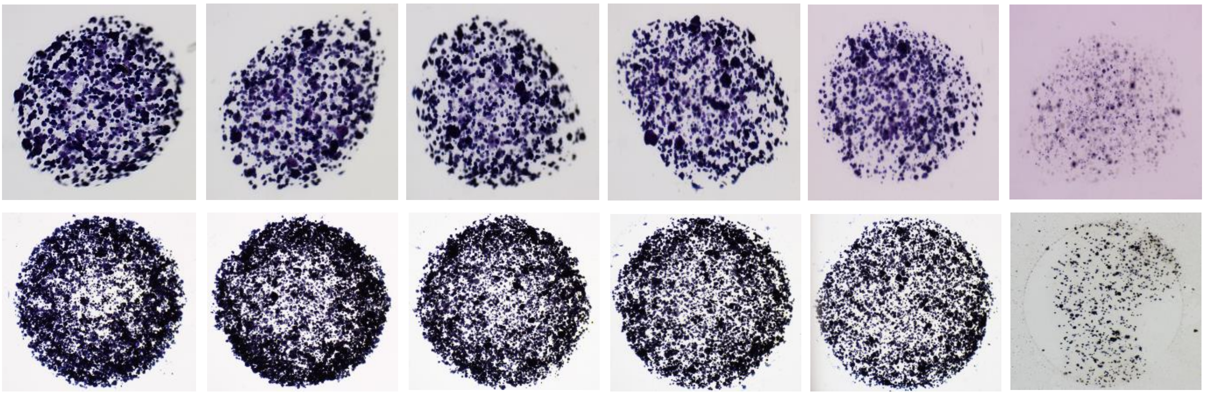


**Cytotoxicity of REST inhibitors in colon organoids.** To more accurately assess the toxicity of two REST inhibitors *in vitro*, organoids were generated from Col33, a primary culture of colorectal carcinoma. For each dome in a 12-well chamber slide, 5,000 cells were suspended in 10 µL of Geltrex matrix. The organoids were cultured in neuronal medium for three days prior to drug treatment. They were then exposed to the REST inhibitors X5050 and GSK126 at concentrations ranging from 50 to 3.12 µg/mL for 72 hours. Following drug exposure, MTT solution was added to achieve a final concentration of 0.5 mg/mL. After incubation, the supernatant was removed, and the organoids were imaged using a stereoscope. Because the drugs were dissolved in DMSO, a vehicle control containing the same DMSO volume as in the highest drug concentration was included. The results presented are representative of three independent experiments.
